# Early defects in lysosomal storage diseases disrupt excitatory synaptic transmission

**DOI:** 10.1101/2020.07.06.186809

**Authors:** Camila de Britto Pará de Aragão, Luigi Bruno, Poulomee Bose, Xuefang Pan, Chanshuai Han, Peter S. McPherson, Erika Freemantle, Jean-Claude Lacaille, Éric Bonneil, Pierre Thibault, Claire O’Leary, Brian Bigger, Carlos R. Morales, Graziella DiCristo, Alexey V. Pshezhetsky

**Affiliations:** CHU Sainte-Justine Research Center, University of Montreal, Montreal, H3T 1C5, QC, Canada; Department of Anatomy and Cell Biology, McGill University, Montreal, H3A 0C7, QC, Canada; Department of Neurology and Neurosurgery, Montreal Neurological Institute, McGill University, Montréal, H3A 2B4, QC, Canada; Department of Neurosciences, Faculty of Medicine, University of Montreal, H3T 1J4, QC, Canada; Proteomic Platform, Institute for Research in Immunology and Cancer, University of Montreal, Montreal, H3T 1J4, QC, Canada; Stem Cell & Neurotherapies, Division of Cell Matrix Biology and Regenerative Medicine, School of Biological Sciences, Faculty of Biology Medicine and Health, University of Manchester, Manchester, M13 9PT, United Kingdom

**Keywords:** synaptic transmission, mucopolysaccharidosis, heparan sulfate, acetyl-CoA: α-glucosaminide N-acetyltransferase

## Abstract

At least two thirds of patients affected with lysosomal storage disorders (LSD) exhibit neurological symptoms. For mucopolysaccharidosis type IIIC (MPS IIIC, Sanfilippo disease type C) caused by mutations in the *HGSNAT* gene and lysosomal storage of heparan sulfate the major burden is progressive and severe neuropsychiatric problems, mental retardation, and dementia though to be mainly mediated by neurodegeneration. HGSNAT knockout mice match human clinical phenotype and develop hyperactivity followed by memory impairment and death.

In order to understand whether early clinical symptoms in MPS IIIC mice occurring before the onset of massive neurodegeneration are caused by neuronal dysfunction we studied synaptic transmission and morphology in cultured hippocampal and CA1 pyramidal neurons of MPSIIIC mice. Synaptic spines were also studied in other mouse LSD models and postsynaptic densities in post-mortem cortices of human neurological MPS patients.

Cultured hippocampal and CA1 pyramidal neurons of MPS IIIC mice showed a drastic decrease or abnormal distribution of multiple pre- and postsynaptic proteins that could be rescued in vitro and in vivo by virus-mediated gene correction. Dendritic spine densities were immature in cultured hippocampal MPS IIIC mouse neurons and reduced in pyramidal neurons of mouse models of MPS IIIC and other (Tay-Sachs, sialidosis) LSD starting from postnatal day 10. MPS IIIC neurons presented alterations in frequency and amplitude of miniature excitatory and inhibitory postsynaptic currents, sparse synaptic vesicles, reduced postsynaptic densities, disorganised microtubule network and partially impaired axonal transport of synaptic proteins. Postsynaptic densities were also reduced in post-mortem cotrees of human MPS I, II, IIIA, C and D patients suggesting that the pathology is common for these neurological LSD.

Together, our results demonstrate that lysosomal storage causes alterations in synaptic structure and abnormalities in neurotransmission originating from disrupted vesicular transport and preceding the first cognitive symptoms and suggest drugs known to affect synaptic transmission can be potentially applied to treat behavioral and cognitive defects in neurological LSD patients.

## Background

Lysosomal storage diseases (LSD) are progressive, pediatric multi-systemic disorders, with a typical cellular landmark of “storage bodies” caused by lysosomal accumulation of undigested macromolecules. More than two-thirds of the LSD present with central nervous system (CNS) involvement causing cognitive or motor impairment (reviewed in (1)). Neurological manifestations are particularly common among mucopolysaccharidoses (MPS), which comprise approximately 30% of cases of LSD (2). Progressive and severe neurological decline is the major burden for Mucopolysaccharidosis type III (MPS III), also known as Sanfilippo syndrome, a rare genetic neurogenerative disease manifesting with neuropsychiatric problems, such as hyperactivity, aggressiveness and autistic features, followed by mental retardation, speech and hearing loss and dementia (3). Most patients develop gait disorders, become wheelchair-bound during adolescence and die before adulthood. In some cases, however, the survival may exceed the fourth decade of life (3) with progressive dementia and retinitis pigmentosa (4–7). MPS III is the most prevalent MPS disorder occurring with a combined frequency of 0.28–4.1 per 100,000 live births (2, 8). Four subtypes, A to D, of the disease are associated with deficiencies of specific enzymes catalyzing subsequent steps of heparan sulphate (HS) catabolism (4, 5): sulfamidase (MPS IIIA, OMIM 252900)(9), α-N-acetylglucosaminidase (MPSIIIB, OMIM 252920)(10), heparan acetyl CoA: α-glucosaminide N-acetyltransferase (HGSNAT, MPSIIIC, OMIM 252930)(11) and N-acetylglucosamine-6-sulfatase (MPSIIID, OMIM 252940)(12).

Our previous studies of the *Hgsnat^Geo^* mouse model of MPS IIIC revealed that these animals develop hyperactivity and learning impairment at 5-6 months, followed by urinary retention and death, at 10-11 months of age (13). Analysis of pathological changes in the brain demonstrated that primary accumulation of HS in the microglial cells and neurons causes impaired autophagy, secondary neuronal storage of G_M2_/G_M3_ gangliosides and misfolded proteins (13), neuroinflammation, as well as abnormalities in mitochondrial energy metabolism, leading eventually to neuronal death (13). Importantly, the neuronal loss in the MPS IIIC mice becomes significant only after 10 months of age whereas the behavioural abnormalities manifest starting from 4 months. In human MPS IIIA and MPS IIIB patients imaging studies have shown a correlation between gray matter deterioration and decrease of development quotient scores (14–16), however the degree of atrophic changes does not always match clinical severity. Some reported patients with a severe clinical outcome demonstrated moderate MRI alterations (17), others showed an extensive neuronal degradation but had relatively mild neuropsychiatric signs (18). Together, these data suggest that neurobehavioral and cognitive manifestations in animal MPS III models and human patients are also caused by functional pathological changes within the neurons.

Synaptic pathology has been previously reported in several animal models of LSD including feline models of GM1-gangliosidosis (19, 20), mouse model of Niemann-Pick C disease (21) and the Twitcher mouse, the natural mouse model of Krabbe disease (22). Recently, reduced excitatory synaptic strength on the somatosensory cortex was shown in the mouse model of MPS IIIA (23). For the same MPS IIIA mouse model another study reported inhibition of soluble NSF attachment receptor (SNARE) complex assembly and synaptic vesicle recycling which authors proposed to be caused by perikaryal accumulation of insoluble α-synuclein and increased proteasomal degradation of cysteine string protein α (CSPα) resulting in low availability of these proteins at the nerve terminal (24). However, both α-synuclein accumulation and CSPα deficiency manifest in 8-month-old MPS IIIA mice (24), when these animals are at the advanced/terminal stage of the disease concomitant with the massive neuronal death. It is, therefore, implausible that these changes are the underlying cause of the early symptoms of disease such as hyperactivity and deficits of working memory that appear in this model as early as at 3-4 months likely marking presence of defects in synaptic neurotransmission and/or long-term-potentiation (25).

In the current study, we demonstrate that synaptic pathology in hippocampal neurons of the MPS IIIC mouse, including deficits in synaptic spines, synaptic vesicles and abnormalities in neurotransmission, are very early events appearing during the postnatal period, long before the emergence of the first noticeable symptoms. Moreover, our data present evidence that defects in glutaminergic neurotransmission originating from defects in axonal transport is a recurring feature of multiple neurological LSD affecting in particular human patients with wide range of MPS disorders and should be considered as an essential part of pathological CNS changes and a potential therapeutic target for these conditions.

## Methods

### Animals

The mouse models of MPSIIIC (*Hgsnat^Geo^*), sialidosis (*Neu1^-/-^*) and Tay-Sachs (*Hexa^-/^*^-^) in C57BL/6J genetic background have been previously described (13, 26–28). The Thy1-EGFP transgene mice expressing enhanced green fluorescent protein (EGFP) under the control of a modified regulatory region of the mouse thy1.2 gene promoter (containing the sequences required for neuronal expression but lacking the sequences required for expression in non-neural cells) were obtained from The Jackson Laboratory (JAX stock #007788)(29). All mice were bred and maintained in compliance with the Canadian Council on Animal Care (CCAC) in the accredited animal facility of the CHU Sainte-Justine. Mice were housed in an enriched environment with continuous access to food and water, under constant and controlled temperature and humidity, on a 12 h light/dark cycle. All the experiments performed on mice have been approved by the Animal Care and User Committee of the Ste-Justine Hospital Research Center. The experiments were conducted for both male and female mice and the data analyzed to determine the differences between sexes. Because no differences between sexes were observed in the experiments conducted in this study the data for males and females were combined. The neuronal cultures were established from pooled hippocampi of mouse embryos of both sexes.

### Neuronal cultures and transduction

Primary hippocampal neuronal cultures were established from mouse brain tissues at embryonic day 16. The hippocampi were dissected, treated with 2.5% trypsin (Sigma-Aldrich, T4674) for 15 minutes at 37°C, washed 3 times with Hank’s Balanced Salt Solution (HBSS, Gibco, 14025-092) and mechanically dissociated using borosilicate pipets with opening sizes of 3, 2 and 1 mm. The cells were then counted with the viability dye trypan blue (Thermo Fisher Scientific, 15250061) in a hemocytometer and resuspended in Neurobasal media (Gibco, 21103-049) supplemented with B27, N2, penicillin and streptomycin. The hippocampal cells were plated at a density of 60,000 cells per well in a 12-well plate, on Poly-L-Lysine hydrobromide (Sigma Aldrich, P9155) coated coverslips. Cells were cultured for 21 days and 50% of media was changed on days 3, 10 and 17. Primary hippocampal neurons were transduced at day *in vitro* (DIV) 3 with lentivirus (LV) encoding for HGSNAT fused to green fluorescent protein (GFP) under a cytomegalovirus (CMV) promoter, for the rescue experiments, or with LV encoding for synapsin 1 (Syn1) fused to GFP (GeneCopoeia, Inc., cat.# LPP-Z5062-Lv103-100), for live-cell imaging. Transduction was performed in the presence of 8 µg/ml of protamine sulfate for 15 h.

### Analyses of postsynaptic densities (PSD) in human brain tissues

Frozen or fixed with paraformaldehyde (PFA), cerebral cortices from clinically confirmed MPS patients (1 case of MPS I, 1 case of MPS II, 2 cases of MPSIIIA, 1 case of MPSIIIC and 2 cases of MPSIIID) and age-matched controls with no pathological changes in the central nervous system were provided by NIH NeuroBioBank (project 1071, MPS Synapse). Upon arrival to the laboratory, the samples were embedded in Tissue-Tek® optimum cutting temperature (OCT) Compound and stored in - 80°C. Brains were cut in 40 µm sections and stored in cryopreservation buffer (0.05 M sodium phosphate buffer pH 7.4, 15% sucrose, 40% ethylene glycol) at −20°C until labelling for immunofluorescence.

### Immunofluorescence

Cultured neurons at DIV 21 were fixed in 4% PFA/4% sucrose solution in PBS, pH 7.4, washed 3x with PBS and stored at 4°C for posterior analysis. Mouse brains were collected from animals perfused with PBS followed by perfusion with 4% PFA and left in 4% PFA overnight. Brains were placed in 30% sucrose for 2 days and then they were embedded in Tissue-Tek® OCT Compound and stored in −80°C. Brains were cut in 40 µm sections and stored in cryopreservation buffer (0.05M sodium phosphate buffer pH 7.4, 15% sucrose, 40% ethylene glycol) at −20°C until labelling. The cultured neurons were permeabilized with 0.1% Triton X-100 and blocked with 5% goat serum in PBS for 1 h, and then incubated overnight at 4°C with primary antibodies in 5% goat serum. Brain sections (mouse or human) were washed 3 times with PBS, and permeabilized/blocked by incubating in 5% bovine serum albumin (BSA) in 1% Triton X-100 for 2 h. Hybridization with primary antibodies dissolved in 1% BSA in 0.1% Triton X-100 was performed overnight at 4°C. The following antibodies were used: rabbit anti-mouse synapsin 1 (1:200, Abcam, catalog ab64581), rabbit anti-mouse synaptophysin (1:300, Millipore Sigma, catalog SAB4502906), rat anti-mouse lysosomal-associated membrane protein 1 (LAMP1, 1:200, DSHB, catalog 1D4B-s), mouse anti-mouse heparan sulfate (10E4 epitope, 1:100, AMSBIO, catalog F58-10E4), rabbit anti-mouse NeuN (1:250, Millipore Sigma, catalog MABN140), mouse anti-mouse β-tubulin III (1:400, Millipore Sigma, catalog T2200), mouse humanized anti-G_M2_ (1:400, KM966), chicken anti-mouse microtubule-associated protein 2 (MAP2, 1:2000, Abcam, catalog ab5392) rabbit anti-mouse vesicular glutamate transporter 1 (vGlut1, 1:1000 for cells, 1:200 for tissues, Abcam, catalog ab104898), mouse anti-mouse postsynaptic density protein 95 (PSD-95, 1:1000 for cells, 1:200 for tissues, Abcam, catalog ab99009), rabbit anti-mouse vesicular gamma aminobutyric acid (GABA) transporter (vGAT, 1:1000, Synaptic Systems, catalog #131003), mouse anti-mouse Gephyrin (1:1000, Synaptic Systems, catalog #147021), Neuroligin-1 (1:300, Abcam, catalog ab26305), mouse anti-mouse neurofilament medium chain (1:200, DSHB, catalog 2H3-s). The cells and tissues were washed 3x with PBS and counterstained with Alexa Fluor 488-, Alexa Fluor 555- or Alexa Fluor 633- conjugated goat anti-rabbit, anti-mouse, anti-rat or anti-chicken IgG (1:1000 for cells, 1:400 for tissues, all from Thermo Fisher Scientific) for 1 h for cells or 2 h for tissues at room temperature and the nuclei were stained with Draq5 (1:1000, Thermo Fisher Scientific) for 40 min or 4′,6- diamidino-2-phenylindole (DAPI) within the mounting media.

To visualize the synaptic spines in cultured neurons, on DIV 21 the culture media was replaced by 2.5 µM 1,1’-Dioctadecyl-3,3,3’,3’-Tetramethylindocarbocyanine Perchlorate (Dil, Thermo Fisher Scientific) dissolved in fresh media and the cells were incubated for 15 min at 37 °C, washed 3x with media preheated to 37 °C, and fixed with 1.5% PFA in PBS for 15 min. The slides were mounted with Prolong Gold Antifade mounting reagent (Thermo Fisher Scientific) and analyzed using Leica DM 5500 Q upright confocal microscope (63x oil objective, N.A. 1.4, zoom 1.5). Images were processed and quantified using ImageJ 1.50i software (National Institutes of Health, Bethesda, MD, USA). Quantification was double-blinded and performed for at least 3 different experiments. Quantifications of synaptic spine density and puncta were performed in 20 µm of dendrite length, respectively, 30 µm away from the neuronal soma.

### Transmission electron microscopy

At 3 and 6 months, 3 mice per genotype were anesthetized with sodium pentobarbital and transcardiacally perfused with PBS followed by 2.5% glutaraldehyde in 0.2 M phosphate buffer (pH 7.2). The brains were extracted and post fixed in the same fixative for 24 h at 4°C. The hippocampi were dissected in blocks of 1 mm^3^ and sections of 1 µm thickness were cut, mounted on glass slides, stained with toluidine blue and examined on a Leica DMS light microscope to select the CA1 region of the hippocampus for the electron microcopy. For the neuronal cultures, cells at DIV 21 were washed with PBS and fixed overnight in 2.5% glutaraldehyde in 0.1 M sodium cacodylate buffer (pH 7.4, Electron Microscopy Sciences) at 4°C. Both tissue and cultured cell samples were stained with 1% osmium tetroxide (Mecalab) and 1.5 % potassium ferrocyanide (Thermo Fisher Scientific) followed by dehydration in a graded series of ethanol (30%-90%) and embedded in Epon. The polymerized blocks were trimmed, and 100 nm ultrathin sections were cut with an Ultracut E ultramicrotome (Reichert Jung), mounted on 200-mesh copper grids (Electron Microscopy Sciences), stained with uranyl acetate (Electron Microscopy Sciences) and lead citrate (Thermo Fisher Scientific) and examined on a FEI Tecnai 12 transmission electron microscope (FEI Company) operating at an accelerating voltage of 120 kV equipped with an XR-80C AMT 8 megapixel CCD camera. For quantification, the micrographs were taken with 13,000 x and 49,000 x magnification.

### Whole-cell patch clamp recordings in dissociated hippocampal neuronal cultures

Experiments were performed after DIV 22-25 on dissociated hippocampal neuronal culture. Cultures were maintained in a humidified atmosphere of 5% CO_2_ and 95% O_2_ at 37°C. Coverslips with cultured hippocampal neurons were placed in a recording chamber mounted on an inverted microscope (Nikon Eclipse Ti-S). Whole-cell recordings were obtained from neurons, selected by visual identification in phase contrast, using borosilicate pipettes (3–6 MΩ). The intracellular solution for recording miniature excitatory postsynaptic currents (mEPSCs) and inhibitory postsynaptic current (mIPSCs) contained (in mM) 132 CsMeSO_3_, 8 CsCl, 0.6 EGTA, 10 diNa-phosphocreatine, 10 HEPES, 4 ATP-Mg^2+^, 0.4 GTP-Na_3_. It was filtered, and adjusted to pH 7.25-7.30 with 275-280 mOsmol CsOH. mEPSCs and mIPSCs were recorded in the presence of tetrodotoxin (TTX) (1 µM; Abcam) in artificial cerebrospinal fluid (ACSF) containing (in mM) 132.3 NaCl, 3 KCl, 15 HEPES, 1.25 NaH_2_PO_4_, 2 CaCl_2_, 2 MgSO_4_, 10 D-Glucose. ACSF was adjusted to pH 7.37-7.41 with 295-305 mOsm NaOH and perfused at 1 ml/min. Recordings were obtained using a Multiclamp 700B amplifier and a 1440A Digidata acquisition board (Molecular Devices). Signals were low-pass-filtered at 2 kHz, digitized at 20 kHz and stored on a PC. Upon whole cell configuration, holding potential was initially maintained at −70 mV for mEPSCs and subsequently reduced to 0 mV for mIPSCs for 5 min recordings of each in the same cell. Signal quality was routinely monitored and recordings were only included if access resistance was less than 30 MΩ and varied for less than 25%. For analysis, mEPSCs and mIPSCs were Bessel filtered at 2.8 kHz using pClamp10 software (Molecular Devices). Miniature events were analyzed using MiniAnalysis (Synaptosoft). Unpaired-sample t-tests (p<0.05; two-sided) were used for statistical comparison of mEPSC and mIPSC frequencies, amplitudes, rise and decay times.

### Whole cell recordings in acute hippocampal slices

Acute hippocampal slices were prepared as described (30). Briefly, animals (P45-60) were anaesthetized deeply with isoflurane and decapitated. The brain was dissected carefully and transferred rapidly into an ice-cold (0–4°C) solution containing the following (in mM): 250 sucrose, 2 KCl, 1.25 NaH_2_PO_4_, 26 NaHCO_3_, 7 MgSO_4_, 0.5 CaCl_2_ and 10 glucose, pH 7.4. The solution was oxygenated continuously with 95% O_2_ and 5% CO_2_, 330–340 mOsm/L. Transverse hippocampal slices (thickness, 300 µm) were cut using a vibratome (VT1000S; Leica Microsystems), transferred to a bath at room temperature (23°C) with standard ACSF at pH 7.4 containing the following (in mM): 126 NaCl, 3 KCl, 1 NaH_2_PO_4_, 25 NaHCO_3_, 2 MgSO_4_, 2 CaCl_2_, 10 glucose, continuously saturated with 95% O_2_ and 5% CO_2_ and allowed to recover for 1 h. During the experiments, slices were transferred to the recording chamber at physiological temperature (30–33 °C) continuously perfused with standard ACSF, as described above, at 2 ml/min. Pyramidal CA1 neurons from the hippocampus were identified visually using a 40X water immersion objective. Whole-cell patch-clamp recordings were obtained from single cells in voltage- or current-clamp mode and only 1 cell per slice was recorded to enable post-hoc identification and immunohistochemical processing. Recording pipettes (4–6 MΩ) were filled with a K-gluconate based solution for voltage-clamp recordings (in mM): 130 K-gluconate, 10 KCl, 5 diNa-phosphocreatine, 10 HEPES, 2.5 MgCl_2_, 0.5CaCl_2_,1 EGTA, 3 ATP-Tris, 0.4 GTP-Li, 0.3% biocytin, pH 7.2–7.4, 280–290 mOsm/L.

After obtaining whole cell configuration, passive membrane properties were monitored for 5 min and current clamp recordings were done to measure action potential characteristics. Slices were then perfused with 0.5 μM TTX (to isolate miniature events) for 3 mins before commencing voltage clamp recordings. Cells were voltage clamped at −70 mV for mEPSCs recording and then held at 0 mV (calculated from the reversal potential of Cl) for mIPSCs recording. Data acquisition (filtered at 2–3 kHz and digitized at 15 kHz; Digidata 1440A, Molecular Devices, CA, USA) was performed using the Axopatch 200B amplifier and the Clampex 10.6 software (Molecular Devices). Both mEPSCs and mIPSCs were recorded for 7 min and a running template on a stable baseline (minimum of 30 events) was used for the analysis of miniature events on MiniAnalysis. Clampfit 10.2 software was used for analysis of action potential characteristics and other passive membrane properties.

For some experiments, to verify that all mEPSCs are blocked, slices were perfused with 5 μM DNQX (6,7-dinitroquinoxaline-2,3-dione) and 50 μM AP5 in addition to the TTX while recording mEPSCs at −70 mV after addition of α-amino-3-hydroxy-5-methyl-4-isoxazolepropionic acid (AMPA) and N-methyl-D-aspartate receptor (NMDAR) blockers. Similarly, for some experiments, slices were perfused with 100 μM BMI (bicuculline methiodide) and 50 μM AP5 in addition to the TTX in the ACSF to verify that all mIPSCS are blocked at 0 mV.

### Isolation of synaptosomes

At 3 and 6 months of age, mice (3 per genotype) were anesthetized with sodium pentobarbital and sacrificed by cranial dislodgement. Brains were removed, weighted and placed in a cold Dounce tissue grinder with 2 mL of Syn-PER Synaptic Protein Extraction Reagent (Thermo Scientific) per 200 mg of brain tissue in the presence of protease and phosphatase inhibitor cocktails (cOMplete Tablets EDTA-free and PhosSTOP EASYpack tablets, Roche). The brains were homogenized on ice with 10 slow strokes, transferred to centrifuge tubes and centrifuged at 1,200 *g* for 10 min at 4°C. The supernatants were collected and centrifuged at 15,000 *g* for 20 min at 4°C. The supernatant, corresponding to the cytosolic fraction was collected and the pellet (synaptosomes) was resuspended with the Syn-PER reagent (500 µL per 200-400 mg of brain tissue). The concentration of proteins in the synaptosomes was measured using the Bio-Rad Bradford kit.

### Liquid Chromatography / Mass Spectrometry

Each sample was diluted with 630 µL of 50 mM ammonium bicarbonate containing 5 mM TCEP [Tris(2-carboxyethyl) phosphine hydrochloride; Thermo Fisher Scientific], 20 mM 2-chloroactamide and vortexed for 1 h at 37°C. One μg of trypsin was added, and digestion was performed for 8 h at 37°C. Digestion was stopped with the addition of 6.3 μL of trifluoroacetic acid (TFA). The extracted peptide samples were dried down and solubilized in 5% ACN-0.2% formic acid (FA). The samples were loaded on a home-made C18 precolumn (0.3-mm inside diameter [i.d.] by 5 mm) connected directly to the switching valve. They were separated on a home-made reversed-phase column (150-μm i.d. by 150 mm) with a 56-min gradient from 10 to 30% ACN-0.2% FA and a 600-nL/min flow rate on a Ultimate 3000 nano-LC connected to an Q-Exactive Plus (Thermo Fisher Scientific, San Jose, CA). Each full MS spectrum acquired at a resolution of 70,000 was followed by 12 tandem-MS (MS-MS) spectra on the most abundant multiply charged precursor ions. Tandem-MS experiments were performed using collision-induced dissociation (HCD) at a collision energy of 27%. The data were processed using PEAKS 8.5 (Bioinformatics Solutions, Waterloo, ON) and a mouse database. Mass tolerances on precursor and fragment ions were 10 ppm and 0.01 Da, respectively. Variable selected posttranslational modifications were carbamidomethyl (C), oxidation (M), deamidation (NQ), and phosphorylation (STY). The data were visualized with Scaffold 4.8.7 (protein threshold, 99%, with at least 2 peptides identified and a false-discovery rate [FDR] of 1% for peptides).

### Live-cell imaging

Primary hippocampal neurons were plated on poly-L-lysine coated 4- well chambered glass slides at a density of 6 x10^4^ cells. Neurons were transduced with LV-Syn1-GFP at DIV 3 and were imaged at DIV 21 using an inverted spinning disk confocal microscope (Leica DMi8, 63x objective). During image recording the cells were maintained in a humidified atmosphere of 5% CO_2_ at 37°C. Digital images were acquired with an EM-CCD camera. Cells were recorded every 2 s for a total of 600 s for 5 z-stacks of 0.4 µm each. The videos were played with 10 frames per second, all stacks combined. Kymographs were generated by analyzing the traffic of synaptic vesicles using ImageJ 1.50i software (National Institutes of Health, Bethesda, MD, USA).

### Western blots

Brain tissues (frontal part of a hemisphere, approximately 25% of the brain) were homogenized in RIPA buffer (50 mM Tris-HCl pH 7.4, 150 mM NaCl, 1% NP-40, 0.25% sodium deoxycholate, 0.1% SDS, 2 mM EDTA, 1 mM PMSF, Roche protease and phosphate inhibitor cocktails, 2.5 ml per 1 g of tissue). Neuronal cells grown in culture were scraped in the same buffer (0.1 ml per million cells). The homogenates were kept on ice for 30 min and centrifuged at 13,000 RPM at 4°C for 25 minutes. The supernatant was collected and centrifuged again for 15 min. The resulting lysates were separated by SDS-PAGE on 8% gels. Western blot analyses were performed according to standard protocols using the following antibodies: synapsin 1 (rabbit, 1:2000, Abcam, catalog ab64581), PSD-95 (rabbit, 1:2000, Abcam, catalog ab18258), Munc18-1 (rabbit, 1:3000, Abcam, catalog ab3451), calcium-calmodulin kinase II (CaMKII, mouse, 1:2000, Abcam, catalog ab22609), clathrin heavy chain (rabbit, 1:12000, Abcam, catalog ab21679) and α-Tubulin (1:2000, mouse, DSHB, catalog 12G10). Equal protein loading was confirmed by Ponceau S staining and the bands were quantified using ImageJ software.

### Production of the Lentivirus

Human *HGSNAT* codon-optimized cDNA Tordo (31) fused to a GFP (*HGSNAT-GFP*) was cloned into a pENTR1A vector (Invitrogen, 11813-011) and then transferred to a 3^rd^ generation lentiviral vector plasmid (pLenti PGK Blast DEST (w524-1), Addgene, Plasmid #19065, (32)), using Gateway^TM^ technology according to manufacturer’s instructions. The LV was produced in HEK293T cells cultured in RPMI media (Gibco) to 70-90% confluency and co-transfected with REV 6 µg per plate, VSVG 7.8 µg per plate, pMDL (gag-pol), 15 µg per plate, and *Hgsnat-*GFP, 9 µg per plate plasmids. Polyethylenimine 40 µg/ml (PEI, Linear, MW 25000, Transfection Grade, Polysciences Inc.) was used as a transfection reagent and incubated overnight at 37°C, 5% CO_2_. On the following day, the medium was replaced by Dulbecco’s Modified Eagle’s Medium (DMEM, Gibco^TM^) supplemented with 10% fetal bovine serum (FBS, Wisent Inc.) and 1% Penicillin-Streptomycin (Gibco^TM^) and the cells cultured for 30 hours at 37°C, 5% CO_2_. Medium was collected, filtered using a 0.22 µm low protein binding filter and then centrifuged at 50,000 g for 2 h, at 10°C with maximum acceleration and slow brake. The pellet with the virus was resuspended in 100 µl PBS, aliquoted and stored at −80°C.

### Statistics

All data in the figures represent the mean values ± SD or SEM from multiple experiments. Data were analyzed using the GraphPad Prism version 5.00 software (GraphPad Software, San Diego California USA). The numbers of experiments and replicates analyzed in each experiment are indicated in the figure legends. The statistical significance of differences between two conditions was calculated by Student’s t test or ANOVA with level of significance of p<0.05. LC/MS data were analyzed using the Scaffold software v. 4.8.1.

## Results

### 1. Hippocampal MPSIIIC neurons present immature and scarce synaptic spines in vitro and in vivo

To test if defects in synaptic neurotransmission in hippocampal neurons can be responsible for early-onset deficits in memory and learning in MPSIIIC mice (13), we first analyzed and quantified synaptic spines on the dendrites of hippocampal pyramidal neurons, which play an important role in learning and memory (33). The density and morphology of spines on the dendrites of hippocampal cultured neurons were studied at DIV 21 by confocal microscopy after staining the cell membranes with the lipophilic fluorescent dye Dil. Although total spine density (number of spines per 20 μM of the dendrite) was similar for the WT and MPSIIIC neurons, the cells from MPSIIIC mice had ∼2-fold higher density of immature filopodia spines and similarly decreased density of mature “mushroom” spines (Fig. 1A and B). Importantly, similarly to hippocampal and cortical neurons from the brain of MPSIIIC mice (13), cultured MPSIIIC neurons were storing primary (HS) and secondary (GM2 ganglioside) metabolites (Fig. S1A and B, respectively), while no storage was detected in neurons from WT mice. HS colocalized with LAMP1 (lysosomal-associated membrane protein 1) positive organelles, while GM2 showed only partial colocalization with LAMP1, suggesting that the ganglioside also accumulates in non-lysosomal compartments. The accumulation of both metabolites could also be visualized in the lysosomes of NeuN-negative glial cells present in the neuronal cultures (Fig. S1A and B). Similar to microglia *in vivo*, these cells presented more intense HS/LAMP1 staining than neurons (13). Analysis of primary cultures by transmission electron microscopy (TEM) confirmed the lysosomal storage in the MPSIIIC neurons and microglia (Fig. S1C). Both types of cells showed electron-dense lamellar bodies compatible with secondary storage of gangliosides and misfolded proteins as well as electron-lucent vacuoles that can be attributed to the primary storage of HS (Fig. S1D).

**Figure 1.**
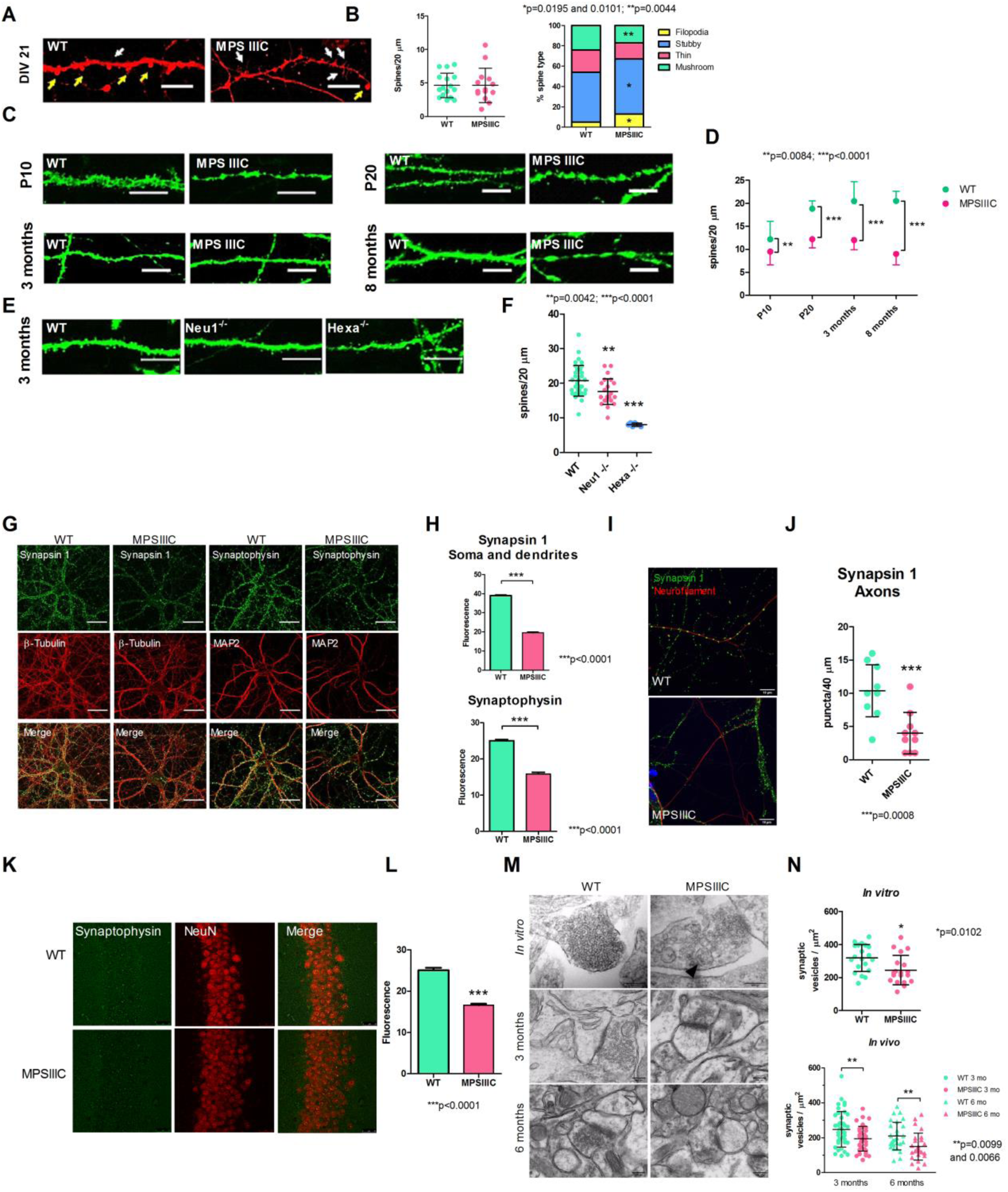
MPS IIIC hippocampal neurons present decreased density of mature synaptic spines, reduced densities of Syn1 and synaptophysin-positive puncta in the axons and scarcity of synaptic vesicles in the synaptic terminals. **(A)** Majority of dendritic spines in MPS IIIC cultured neurons are immature. Dissociated cultured hippocampal neurons at DIV 21 were stained with Dil dye. Mature (mushroom shape) spines are marked with yellow arrows while immature spines (filipodia), with white arrows. **(B)** Quantification of total number of dendritic spines (left) and distribution of different types of spines (right) in hippocampal cultured neurons at DIV 21. **(C)** Representative images and **(D)** quantification of synaptic spines in pyramidal neurons expressing EGFP in the CA1 hippocampus region of mice at the age of P10, P20, 3 months and 8 months. Dendrites in the MPS IIIC neurons have uneven thickness with thin areas alienating with the wider areas resembling spheroids or knots. **(E)** Representative images and **(F)** quantification of synaptic spines in pyramidal neurons from the CA1 region of the hippocampus of 3-month-old *Neu1* KO (mouse model of sialidosis) and *Hexa* KO mice (mouse model of Tay-Sachs disease) and their respective WT littermates heterozygous for the Thy1-EGFP allele. The analysis of spine subtypes and quantification of total spines was performed in a double-blinded manner for 20 µm-long dendrite segments at 30 µm distance from the soma. Data on the graphs show mean values, and SD. P-value was calculated by t-test. Scale bar equals 25 µm for all panels and 10 µm for all enlagrements. At least 30 cells from 3 different mice for each age and genotype were analyzed. **(G)** Representative images of cultured hippocampal neurons at DIV 21 from WT and MPS IIIC mice stained with antibodies against Syn1 and synaptophysin. **(H)** When compared to the WT, MPS IIIC neurons have lower density of Syn1 and synaptophysin-positive puncta. **(I)** Representative images of cultured hippocampal neurons at DIV 21 from WT and MPS IIIC mice stained with antibodies against Syn1 and an axonal marker, neurofilament medium chain protein. **(J)** Quantification of Syn1-positive puncta in the axons. **(K)** Representative images of CA1 pyramidal layer of the hippocampus of 2-month-old MPS IIIC and WT mice stained with antibodies against synaptophysin and a neuronal marker NeuN. **(L)** Reduced density of synaptophysin-positive puncta in MPS IIIC brains. P-value was calculated by t-test, data show mean value and SD of 3 independent experiments or 3 different animals. Scale bar equals 25 µm in **(G)** and **(K)** and 10 µm in **(I)**. (**M**) Representative TEM images of synaptic terminals of cultured hippocampal neurons at DIV 21 and of pyramidal neurons from the CA1 hippocampus region of 3- and 6-month-old mice. An autophagosome in the synaptic terminal is marked with an arrowhead. Scale bar equals 200 nm. **(N)** The density of synaptic vesicles is reduced in MPS IIIC neurons both *in vitro* and *in vivo*. At least 30 cells from 3 mice per each age and genotype were analyzed.

The synaptic spines were further studied *in vivo* on pyramidal neurons in the CA1 region of the hippocampus of *Hgsnat-Geo* mice heterozygous for the *Thy1-EGFP* transgene that expresses the enhanced green fluorescent protein (EGFP) under the control of a modified Thy1 promoter containing the sequences required for neuronal expression but lacking the sequences required for expression in non-neural cells. Mice were compared with double-heterozygous littermates. Synaptic spines were quantified at post-natal day 10 (P10), P20, 3 months and 8 months. As early as at P10, the density of synaptic spines was reduced by ∼22% (WT=12.2±3.9, MPSIIIC=9.5±2.9). The difference between the control and MPSIII groups increased further with age reaching 56% at 8 months (WT=20.5±2.2, MPSIIIC=9±2.4) (Fig. 1C and D).

To assess whether reduced density of synaptic spines is a cellular pathology shared among different types of neurological lysosomal storage disorders, we quantified spines on pyramidal CA1 neurons in the mouse models of Tay-Sachs disease and sialidosis. Human Tay-Sachs patients have mutations in the *HexA* gene encoding for lysosomal β-hexosaminidase A resulting in accumulation of GM2 ganglioside (34), while sialidosis patients have mutations in the neuraminidase 1 (*NEU1*) gene, leading to lysosomal storage of sialylated glycoproteins (35, 36). Previously described *Hexa* KO and *Neu1* KO mice were crossed with the *Thy-1-EGFP* strain, and GFP-positive pyramidal neurons from the CA1 region of the hippocampus were studied by confocal microscopy. Our data show that neurons from 3-month-old sialidosis and Tay-Sachs mice also present reduction of synaptic spine density (WT=20.7±4.4; *Neu1* KO=17.6±3.7; *Hexa* KO =8±0.5; Fig. 1E and F), suggesting that this effect is not exclusively related to heparan sulfate storage.

### 2. Synaptic vesicles are reduced in the terminals of MPSIIIC hippocampal neurons

To test whether the levels of synaptic vesicles are changed in cultured MPSIIIC neurons we studied the cellular distribution and density of synapsin 1 (Syn1) and synaptophysin by confocal immunofluorescent microscopy. In neurons from MPSIIIC mice at DIV 21, the densities of puncta positive for both proteins were significantly reduced (Fig. 1G and H). Density of Syn1-positive puncta was reduced by 50% (WT=39±0.5, MPSIIIC=19.5±0.5) and synaptophysin-positive, by 37% (WT=25±0.7, MPSIIIC=15.9±0.9). We further measured density of Syn1-positive puncta specifically associated with the axonal marker, neurofilament medium chain (NF-M), and also found it to be significantly reduced (Fig. 1I and J). In similar fashion, density of synaptophysin puncta was significantly reduced in pyramidal neurons of the CA1 region of the hippocampus of 3-month-old MPSIIIC mice (Fig. 1K and L).

The density of synaptic vesicles at the synaptic terminals of hippocampal neurons in culture and in pyramidal neurons of the CA1 region of the hippocampus from 3- and 6-month-old mice was further analyzed by TEM. In cultured neurons the density of synaptic vesicles (total number of synaptic vesicles in the axonal terminal divided by the area of the terminal in µm^2^) in the MPSIIIC neurons was decreased by 23% (WT=319.1±81.6, MPSIIIC=245.3±88.7) (Fig. 1M and N, upper panels). A similar phenotype was also observed *in vivo.* In 3-month-old MPSIIIC mice, the density of synaptic vesicles was decreased by 21.7% as compared with WT (WT=248.3±101.7, MPSIIIC=194.3±71.7). This reduction was even more prominent by the age of 6 months (Fig. 1M and N, middle and lower panels), when the difference between MPSIIIC and the WT reached 28.9% (WT=209.6±79, MPSIIIC=149.1±78). Besides, around 10-20% of terminals in the MPSIIIC neurons *in vivo* and *in vitro* contained multivesicular vacuoles with double limiting membrane (Fig. 2G, arrowhead) resembling autophagosomes found in the neurons of patients with adult neurodegenerative disorders (reviewed in (37)). Overall, our TEM results directly confirm the reduction of synaptic vesicles *in vivo* and *in vitro* predicted from decrease in Syn1 and synaptophysin-positive puncta.

**Figure 2.**
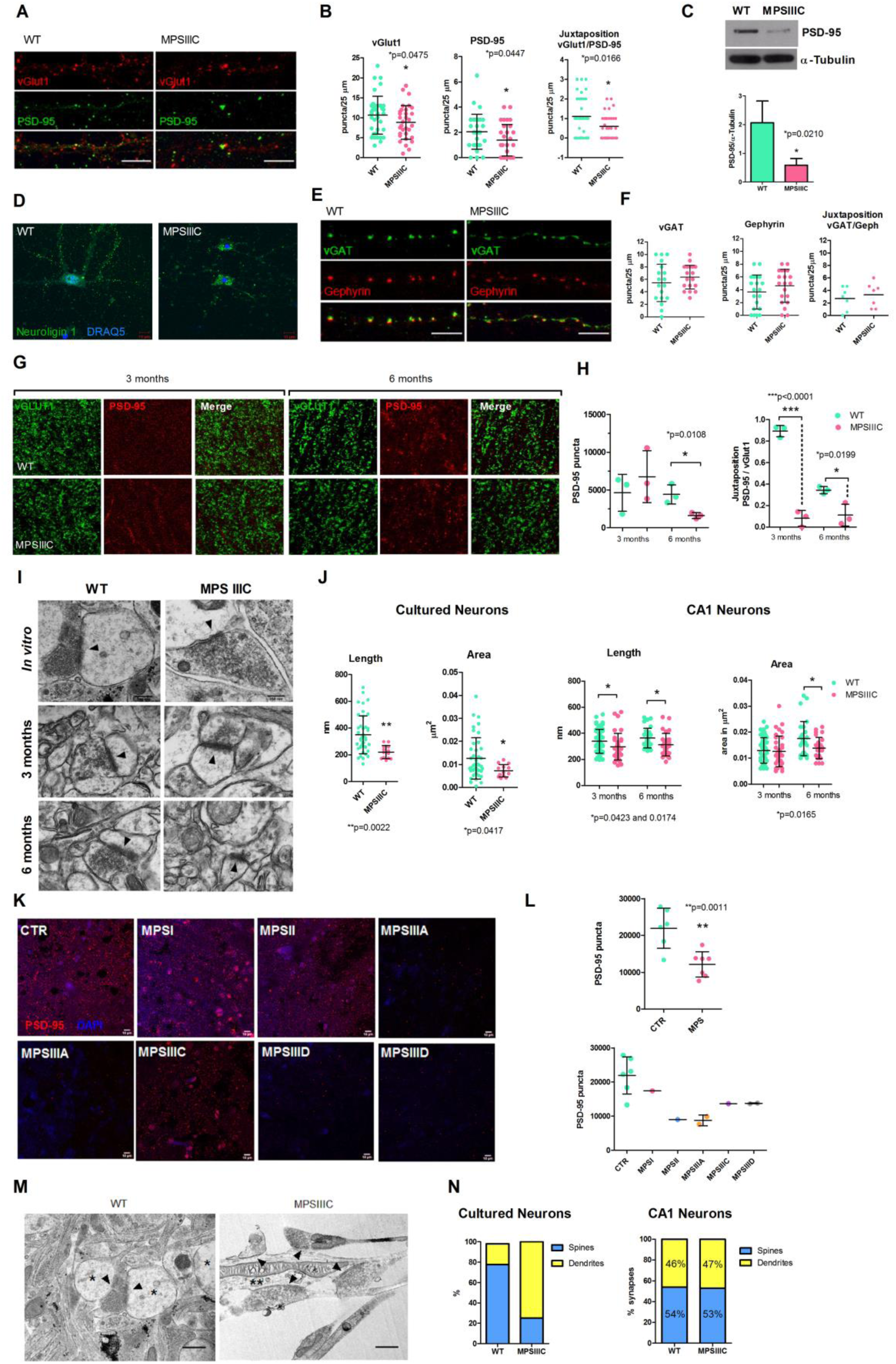
MPS IIIC hippocampal neurons present alterations in the distribution of protein markers of the excitatory synapse and reduction of postsynaptic densities. **(A)** Representative images of hippocampal cultured neuron from at DIV 21 labeled with antibodies against vGlut1 and PSD-95 showing characteristic punctate pattern. **(B)** Quantification of vGlut1-positive puncta, PSD-95-positive puncta and their juxtaposition demonstrates significantly lower densities of PSD-95-positive puncta and PSD-95-positive puncta in juxtaposition with vGLUT1-positive puncta. **(C)** Western blot of protein extracts from cultured hippocampal neurons confirms the reduction of PSD-95 in MPS IIIC cells. **(D)** Representative images of cultured hippocampal neurons stained with antibodies against neuroligin-1 and nuclear marker DRAQ5. In MPS IIIC neurons, neuroligin-1 shows perinuclear accumulation instead of fine puncta observed in the neurites of WT cells. **(E)** Representative images of hippocampal cultured neuron at DIV 21 labeled with antibodies against vGAT and Gephyrin. The insert shows the juxtaposition between the vGAT-positive and Gephyrin-positive puncta indicating functional synapses. **(F)** Quantification of vGAT-positive puncta, Gephyrin-positive puncta and their juxtaposition. The quantification of puncta was performed within 25 µm segments of dendrites at 30 µm distance from the soma in a double-blinded manner using cultures from 3 independent experiments with a total of 10 cells being analyzed for each experiment. P-value was calculated by t-test. Scale bars equal 10 µm. **(G)** Representative confocal images of CA1 hippocampal regions of 3- and 6-month-old WT and MPS IIIC mice stained for the markers of excitatory synapse, PSD-95 and vGLUT1. **(H)** Density of PSD-95-positive puncta and those in juxtaposition with vGLUT1-positive puncta were quantified using ImageJ software. The quantification of puncta was performed in a double-blinded manner using 3 mice per age per genotype. P-value was calculated by t-test. Scale bar = 10 µm. **(I)** Representative TEM images of synapses in cultured hippocampal neurons at DIV 21 and in pyramidal neurons from the CA1 region of the hippocampus of 3- and 6-month-old mice. PSDs are marked with arrowheads. **(J)** Quantification of length (nm) and area (µm^2^) of PSDs in hippocampal cultured neurons and in pyramidal neurons from the CA1 region of the hippocampus. Data show mean values and SD of 3 different sets of experiments with at least 10 images per experiment. Only asymmetric (excitatory) PSDs were considered for analysis. P-value was calculated by t-test. Scale bar equals 250 nm. **(K)** Representative confocal images of post-mortem human cortices from controls and MPS patients stained with antibodies against PSD-95 **(L)** Density of PSD-95-positive puncta in human cortices. **(M)** Representative TEM images of synapses in WT and MPS IIIC cultured hippocampal neurons at DIV 21 showing alteration in the distribution of synapses. In MPSIIIC cultured neurons, there is a shift from axospinous to an axodendritic pattern. Spine (*), synaptic cleft (arrowheads) and dendrite (**). Scale bar equals 500 nm. **(N)** Quantification of axospinous and axodendritic synaptic connections in cultured hippocampal neurons and in the pyramidal neurons in the CA1 region of the hippocampus of 6- month-old WT and MPS IIIC mice. Data show total values of 3 independent experiments with 3 mice per genotype.

### 3. MPSIIIC neurons show alterations in distribution of excitatory synaptic markers

To test whether both excitatory and inhibitory synapses are affected in cultured hippocampal MPS IIIC neurons we studied by confocal immunofluorescent microscopy distribution and density of pre- and postsynaptic markers associated with either glutamate-mediated excitatory or GABA-mediated inhibitory neurotransmission. Puncta were quantified in 25 µm-long dendrite segments, 40 µm away from the soma. We counted separately isolated pre- and postsynaptic puncta and when they were in juxtaposition, indicating the presence of a functional synapse at the moment of fixation. The analysis of the excitatory markers, vGlut1 (the presynaptic transporter of glutamate in the synaptic vesicles) and PSD-95 (postsynaptic density protein 95, a scaffold protein present on the postsynaptic density that interacts with NMDA receptors (38)) demonstrated a significant difference between MPS IIIC and WT neurons (Fig. 2A). Quantification of puncta revealed decreased density of PSD-95 in MSPS IIIC neurons (WT=1.8±1.5 and MPSIIIC=1.4±1.3) and reduced number of PSD-95-positive puncta in juxtaposition with vGLUT1-positive puncta (WT=1.1±1 and MPSIIIC=0.6±0.6), meaning that fewer functional excitatory synapses were occurring in these neurons at the moment of fixation (Fig. 2B). The reduction of PSD-95 in MPS IIIC cultured neurons was confirmed by Western blot (Fig. 2C). Altered cellular localization was also detected for another key protein in the excitatory PSD, neuroligin-1, that binds to PSD-95 and NMDA-R1 receptor (39) and mediates cell-cell interaction in the synapses by binding to neurexins (40). In MPSIIIC cultured hippocampal neurons, the majority of neuroligin-1 accumulated in perinuclear structures in the cell body instead of being localized in the fine puncta in the dendrites as observed in WT neurons (Fig. 2D). Conversely, both density and juxtaposition of puncta for the markers of the inhibitory synapse, vGAT (a transporter involved in the uptake of GABA into the synaptic vesicles) and Gephyrin (a protein that anchors GABA-A receptors) were similar for MPS IIIC and the WT neurons (Fig. 2E, F). We further analyzed whether PSD95-positive puncta and its juxtaposition with vGLUT1-positive puncta were also reduced *in vivo* in the brains of MPS IIIC mice. Brain slices from 3 month and 6-month-old WT and MPSIIIC animals were stained with the corresponding antibodies and the CA1 area of the hippocampus analyzed by confocal microscopy (Fig. 2G). We found that densities of PSD-95-positive puncta and PSD-95-positive puncta in juxtaposition with GLUT1-positive puncta were significantly reduced in the brains of 6-month-old mice (Fig. 2H).

To test if the levels of excitatory PSD are also reduced in the human MPS patients we have analyzed the PFA-fixed somatosensory cortex post-mortem tissues collected at autopsy and donated to the NIH NeuroBioBank. Samples of eight MPS patients (one MPS I, one MPS II, two MPS IIIA, one MPS IIIC, and two MPS IIID) and 7 non-MPS, controls matched for age and sex, were analyzed (project 1071, MPS Synapse). The age and the cause of death, sex, race and available clinical and neuropathological information for the patients and controls are shown in Supplementary Table S1. All MPS patients had complications from their primary disease and died in the first-third decades of life except MPS II patient 902, who died at 42 years of age from MPS II-related pneumonia. None of the patients had received enzyme replacement therapy (ERT) or hematopoietic stem cell transplantation (HSCT). Among non-MPS controls, two died of accident-related injury, two of asthma complications, two of atherosclerotic cardiovascular disease, and one of smoke inhalation (Table S1). This analysis confirmed that densities of PSD-95-positive puncta were significantly reduced in human cortices from MPSIII patients (Fig. 2K-L), thus matching the results obtained in mice.

The length and the area of PSD of asymmetric (excitatory) and symmetric (inhibitory) synapses of cultured hippocampal neurons and pyramidal neurons in the CA1 region of hippocampus from 3 and 6-month-old mice were further studied by electron microscopy (Fig. 2I). The length of excitatory PSD in MPS IIIC hippocampal neurons was generally smaller than in the WT cells (Fig.4D-E). In cultured neurons, the length of PSD for MPS IIIC mice was 37% smaller than WT (WT=349.3±141.1 and MPS IIIC=220.3±48.6 nm). *In vivo*, excitatory PSD length was reduced by 12.5% (WT=339.3±91.5 and MPS IIIC=296.9±101.6) in 3-month-old mice, and by 14% in 6-month-old ones (WT=363.6±73.6 and MPS IIIC=312.2±85.8). The area of excitatory PSD was smaller in cultured neurons and *in vivo* at 6 months but not at 3 months (Fig. 2I-J). In cultured neurons, the area of PSD in µm^2^ of MPSIIIC was reduced by 42% as compared with WT cells (WT=0.013±0.009 and MPS IIIC=0.007±0.003). In 3-month-old mice, the area of excitatory PSD was similar between mouse strains (WT=0.013±0.005; MPS IIIC=0.013±0.006) but by the age of 6 months, it became 21% smaller in MPS IIIC mice (WT=0.018±0.006; MPS IIIC=0.014±0.004). In contrast, PSD length in symmetrical inhibitory synapses was similar between MPS IIIC and WT neurons *in vivo* and *in vitro* (Fig. S2), consistent with immunolabeling results obtained for GABAergic markers (Fig. 2E).

Interestingly, TEM analysis also demonstrated that synapse localisation in cultured MPS IIIC neurons is significantly altered as compared with the WT cells. While in the WT neurons the majority (∼80%) of synapses were located on dendritic spines (axospinous synapses), MPS IIIC cells showed a shift towards axodendritic synapses (78% axodendritic; 22% axospinous, Fig. 2M-N). On the other hand, we did not detect any difference in synaptic localisation on dendrites of CA1 pyramidal cells between 6-month-old WT (54% spines; 46% dendrites) and MPS IIIC (53% spines; 47% dendrites) mice (Fig. 2N).

### 4. MPSIIIC neurons show alterations in neurotransmission

To test if alterations in the distribution of synaptic markers in MPSIIIC mice translate into changes in synaptic transmission we conducted electrophysiological recordings of miniature inhibitory (mIPSCs) and excitatory (mEPSCs) postsynaptic currents. To characterize the synaptic transmission *in vivo*, we performed whole cell patch clamp recordings on acute slices, which showed that both mEPSCs and mIPSCs were altered in MPS IIIC animals as compared to WT. mEPSC amplitude (WT= 26.6±2.52 pA; MPS IIIC= 16.94 ±1.53 pA) and frequency were reduced (WT= 0.75±0.028; MPS IIIC= 0.37±0.04) in MPS IIIC mice (Fig. 3A-B). No significant difference in mEPSC kinetics between the two animal groups was observed (decay τ: MPS IIIC =2.288±0.23; WT=2.095±0.35; rise time: MPS IIIC =1.492±0.23; WT=1.552±0.19) (Fig. 3C).

**Figure 3.**
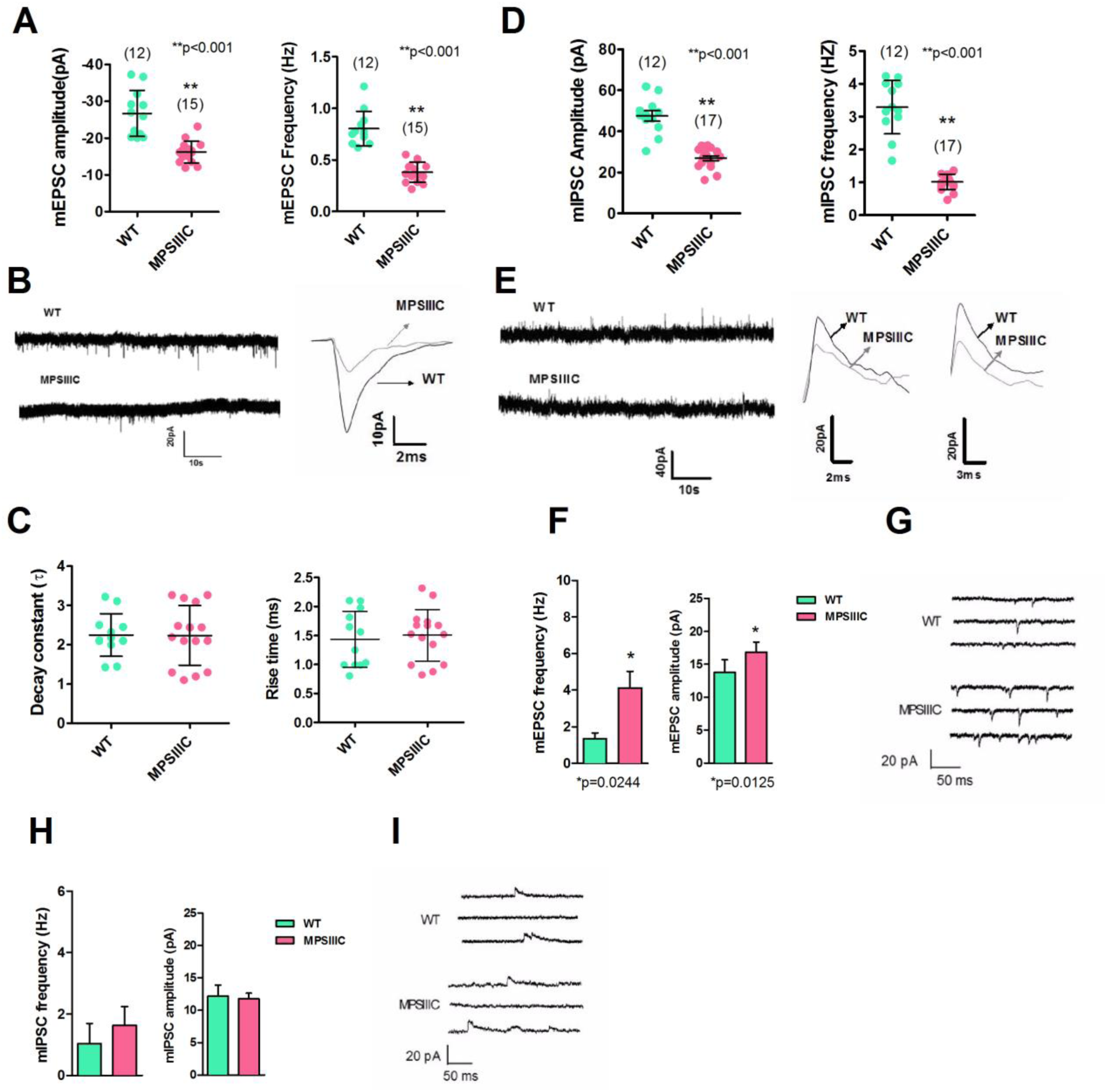
Alteration of miniature inhibitory and excitatory postsynaptic currents in MPS IIIC neurons. **(A)** Significant decrease in the amplitude and frequency (left and right panels, respectively) of mEPSCs as compared to age matched WT controls. Number in parentheses indicate sample size. **(B)** Representative recording of mEPSCs (left) and overlay of representative individual events from MPS IIIC and WT mice (right). **(C)** No significant differences in the decay constant (left) and the rise time (right). **(D)** Significant decrease in the amplitude (left) and frequency (right) of mIPSCs in MPS IIIC animals as compared to age matched WT controls. Number in parentheses indicate sample size. **(E)** Representative recording of mIPSCs from WT and MPS IIIC animals (left panel). Overlay of representative individual mIPSCs from MPS IIIC and WT animals showing events with fast (middle) and slow (right) kinetics. **(F)** Excitatory neurotransmission is enhanced in MPS IIIC cultured hippocampal neurons. Frequency (left) and amplitude (right) of the mEPSCs in WT and MPS IIIC cultured hippocampal neurons (n=11 cells from WT and 17 cells from MPSIIIC) **(G)** Representative traces of the mEPSCs in WT and MPS IIIC cultured hippocampal neurons at DIV19-22. **(H)** Inhibitory neurotransmission mediated by GABA is not affected in MPSIIIC cultured hippocampal neurons. Frequency (left) and amplitude (right) of the mIPSCs in WT and MPS IIIC cultured hippocampal neurons (n=6 cells from WT and 9 cells from MPSIIIC). **(I)** Representative traces of the mIPSCs in WT and MPS IIIC cultured hippocampal neurons at DIV19-22. For mE/IPSCs, the cells originated from at least 8 different neuronal cultures. For all data P-value was calculated by t-test.

Both mIPSC amplitude and frequency were also reduced in MPS IIIC mice as compared with WT (Fig. 3D-E: amplitude, MPS IIIC=26.71±1.96 pA and WT=44.94±3.8 pA; frequency, MPS IIIC=0.82±0.08 Hz and WT= 3.12±0.03 Hz). Upon assessing the kinetics of the mIPSCs, we found two subpopulations of events with decay τ and rise times (Fig. 3E, right panels). Previous studies suggest that different interneuron populations mediate cell–type specific kinetic classes of IPSCs (for review see (41, 42)). However, for both slow and fast mIPSCs the kinetics of the events was not significantly different between MPSIIIC and WT animals (Fig S3).

Upon assessing the membrane properties of hippocampal CA1 neurons from MPS IIIC and WT mice, we found no significant differences in their resting potential, input resistance, action potential threshold, half width of action potential or firing frequency (Supplementary Table S2). This suggests that the intrinsic excitability of CA1 neurons was not significantly different in MPS IIIC mice.

Miniature synaptic events were also recorded in cultured hippocampal neurons from MPS IIIC and WT mice (DIV19-DIV21). We found that mEPSCs of cultured MPS IIIC neurons (Fig. 3F-G) displayed a 3-fold increase in frequency (WT=1.3±1.1 Hz and MPSIIIC=4.1±3.7 Hz) and a significant increase in the amplitude (WT=13.8±6.2 pA and MPSIIIC=16.9±6.3 pA). In contrast, the frequency and amplitude of the inhibitory events in cultured MPS IIIC hippocampal neurons (Fig. 3 H-I) were not significantly different from those in the WT cells (frequency, WT=1±1.6 Hz and MPS IIIC=1.6±1.8 Hz; amplitude, WT=12.2±4.2 pA and MPSIIIC=11.7±2.7 pA).

### 5. PSD-95 deficiency in MPS IIIC neurons can be rescued by correcting primary genetic defect

To establish causative relations between the deficits of protein markers of excitatory synapses and HGSNAT deficiency we attempted to rescue this phenotype by a virus overexpressing the WT human enzyme. Human codon-optimized HGSNAT cDNA fused with that of GFP was cloned into a third-generation lentiviral vector (LV) under the control of a CMV promoter. The LV was tested in HEK 293 cells transduced with a multiplicity of infection (MOI) of 10. The transduced cells expressed HGSNAT-GFP fusion protein correctly targeted to the lysosomes and had 120-fold increased HGSNAT activity as compared with non-transduced cells (Fig. S4).

We further transduced MPS IIIC primary hippocampal neurons at DIV 3 with either LV-GFP or LV-HGSNAT-GFP (Fig. 4A), kept them in culture until DIV 21, fixed and analyzed by confocal fluorescent microscopy to measure density of PSD-95 in the dendrites. Our data (Fig. 4B) show that density of PSD-95 puncta in MPS IIIC hippocampal neurons transduced with LV-HGSNAT-GFP was similar to WT neurons. The density of Syn1-positive puncta associated with NF-stained axons was quantified (Fig. 4C). The number of puncta calculated per 40 µm of the axon was significantly reduced in MPS IIIC neurons as compared with the WT cells and significantly increased in the MPS IIIC cells transduced with LV-HGSNAT-GFP as compared with the LV-GFP-transduced cells (Fig. 4D).

**Figure 4.**
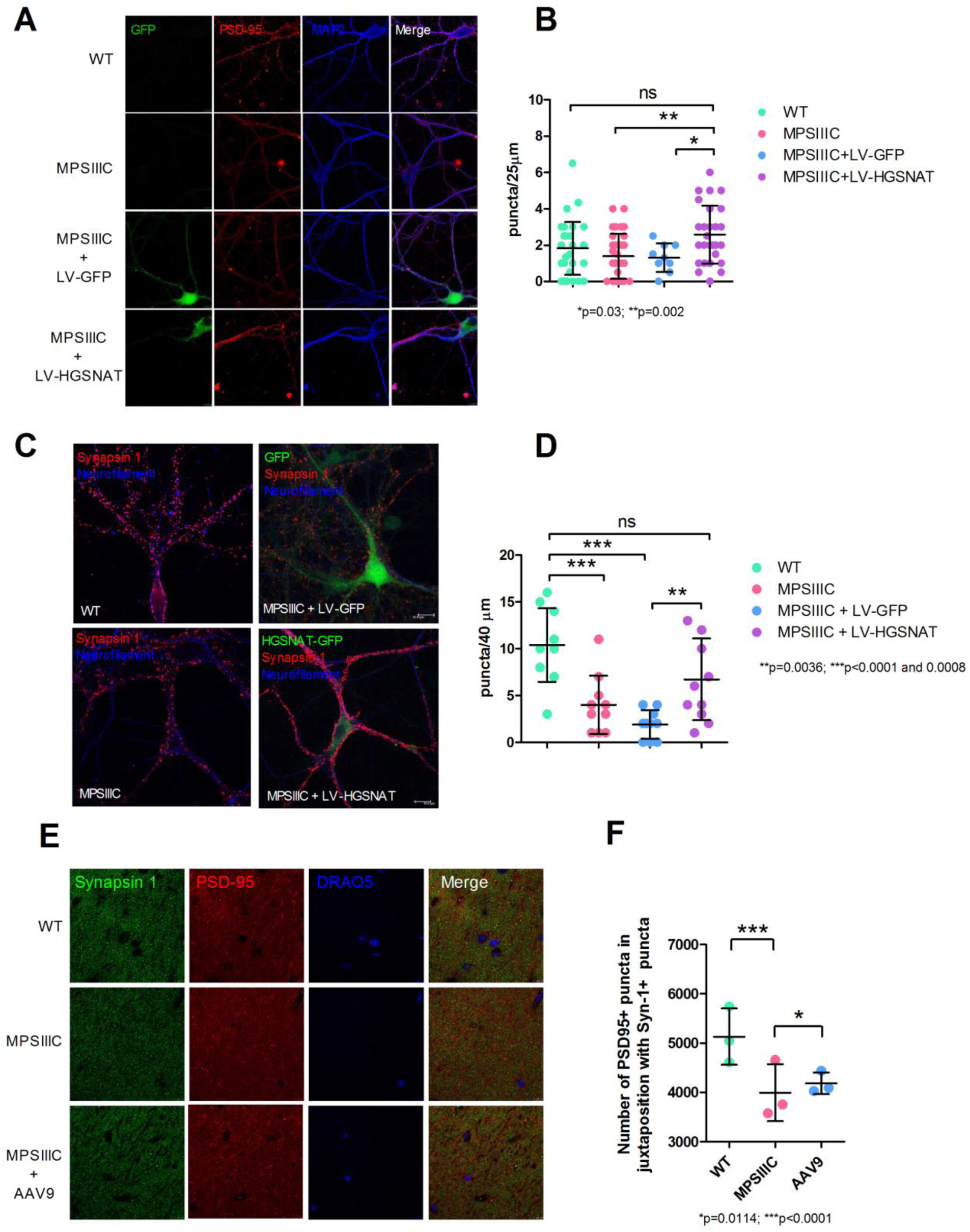
Deficit of PSD-95 and Syn1 in MPSIIIC neurons is rescued *in vitro* and *in vivo* by transduction with viral vectors encoding for WT human HGSNAT. **(A)** Representative images of cultured hippocampal WT, and MPS IIIC neurons and MPS IIIC neurons transduced with either LV-GFP or LV-HGSNAT-GFP stained with anti-PSD-95 antibodies. **(B)** Quantification of PSD-95 puncta in WT, and MPS IIIC cells as well as in MPS IIIC cells treated with LV-GFP, and LV-HGSNAT-GFP. **(C)** Representative images of cultured WT, and MPS IIIC neurons and MPS IIIC neurons transduced with either LV-GFP or LV-HGSNAT-GFP stained with anti-Syn1 antibodies. **(D)** Quantification of Syn1 puncta in the axons of cultured neurons. **(E)** CA1 region of the hippocampus from WT and MPS IIIC mice non-treated or treated with AAV9-HGSNAT stained with anti-Syn1 and anti-PSD-95 antibodies. **(F)** Quantification of Syn1-positive and PSD-95-positive puncta in juxtaposition. Graphs show data from 3 different sets of experiments/mice with 10 images analyzed per experiment. P-value was calculated by one-way ANOVA. Scale bar equals 5 µm in **(A)** and 10 µm in **(C)** and **(E)**.

Previously, we have demonstrated a rescue of the behavioral defects, primary and secondary lysosomal storage and neuroinflammation in MPS IIIC KO mice receiving intracranial injections of AAV vectors (AAV9 and AAV2 true type (TT)) encoding for human untagged HGSNAT (31). Female 8-week-old MPSIIIC and WT mice were injected into each striatum (2 mm lateral and 3 mm deep to bregma) with 2.6 × 10^9^ vg/hemisphere. Control mice received sham injections of saline. After behavioral assessment at the age of 6 and 8 months, mice were sacrificed at 6 months post-treatment and their brains fixed with PFA and cryopreserved. We stained brain slices of AVV9-HGSNAT treated and sham-treated MPS IIIC mice with antibodies against PSD-95 and Syn1 and assessed the density of PSD-95-positive puncta in hippocampal CA1 area (Fig. 4E). We found that the treatment significantly increased the density of the puncta and the density of PSD-95-positive puncta in juxtaposition with Syn1-positive puncta (Fig. 4F). There was also a trend for the increase of Syn1-positive puncta density but the results were not statistically significant perhaps due to high difference between the individual animals in the groups (data not shown). Thus, by restoring the primary defect of HGSNAT we could rescue PSD-95 defects both *in vivo* and *in vitro*.

### 6. Proteomic analyses of synaptosomes reveal a dramatic decrease in synaptic, vesicle trafficking-associated proteins and mitochondrial proteins in MPSIIIC mouse brains

To get insight into the molecular mechanisms underlying synaptic deficits in MPS IIIC neurons, we performed proteomic analyses of mouse brain synaptosomes (43). Synaptosomes were isolated from three MPS IIIC and three WT mice at 3 and 6 months of age, and their protein content analyzed by quantitative proteomics using label-free liquid chromatography tandem mass spectrometry (LC-MS/MS). In synaptosomes from 3-month-old mice, the LC/MS profile identified 1120 proteins in the WT and 1248 proteins in the MPS IIIC (FDR ≤ 1%). In 6-month-old, 809 proteins were identified in the WT and 1246 in the MPS IIIC (Fig. S5A). A volcano plot distribution of synaptosome proteins from 3-month-old mice revealed that 133 proteins were reduced in abundance in MPS IIIC as compared to WT, whereas 395 proteins were reduced in MPSIIIC synaptosomes at 6 months (Fig. S5B, Supplementary Table S3). These proteins were then classified according to their biological function and linked to a particular metabolic or signaling pathway using automated GO (gene ontology terms) annotation (Fig. S5C-D) (44). The 3 protein groups that have shown major reduction in MPS IIIC synaptosomes both at 3 and 6 months were mitochondrial proteins (27.3% of all proteins) synaptic proteins (14%), and proteins involved in vesicle trafficking (5.5 %). In particular, synaptic proteins that were reduced in MPS IIIC mice at both ages (Fig. 5A) and/or showed further reduction with age included: syntaxin-binding protein 1 (Stxbp1/Munc18-1), responsible for docking and fusion of synaptic vesicles in the synaptic terminal; Syn1 and synapsin 2 (Syn2), that coat synaptic vesicles and function in the regulation of neurotransmitter release; calcium/calmodulin-dependent protein kinase II α (Camk2a), a member of the NMDAR signaling complex in excitatory synapses that functions in long-term potentiation and neurotransmitter release; and neuroligin-2 (Nlgn2), a transmembrane protein that acts on the recruitment and clustering of synaptic proteins. In contrast to neuroligin-1 specific to glutaminergic synapse and mislocalized in MPS IIIC cultured neurons (Fig. 2D), Nlgn2 is exclusively localized to inhibitory synapses (45). Specifically, Stxbp1 was reduced by 43.1% in MPS IIIC synaptosomes from 3-month-old mice (WT=36.3±3; MPS IIIC=20.7±5.1) and by 50% by the age of 6 months (WT=44.7±3.2; MPS IIIC=22.3±2). Syn1 was 29.8% lower at 3 months (WT=22.3±7.2; MPS IIIC=15.7±5.5) and reached 41.5% reduction by the age of 6 months (WT=39.3±0.6; MPS IIIC=23±1.7). Syn2 was reduced 51.6% at the age of 6 months (WT=20.7±1.1; MPS IIIC=10±1). Camk2a was reduced by 22.4% at 3 months (WT=16.3±1.1, MPS IIIC=12.7±1.5). Nlgn2 was reduced by 83.3% at 3 months (WT=4±1; MPS IIIC=0.7±0.6). In general, deficiency of synaptic proteins detected by proteomics well correlated with the described above results of confocal immunofluorescent analysis (Syn1, PSD-95 and Nlgn1). At the same time majority of abundant neuronal proteins, for example such as spectrin beta chain, or myelin basic protein were similar in both groups (See Supplementary Table S4). Significant reduction of Syn1, Stxbp1 and Camk2a in the brain (frontal part) homogenates from 6-month-old MPS IIIC mice has been also independently confirmed by Western blots (Fig. 5B).

**Figure 5.**
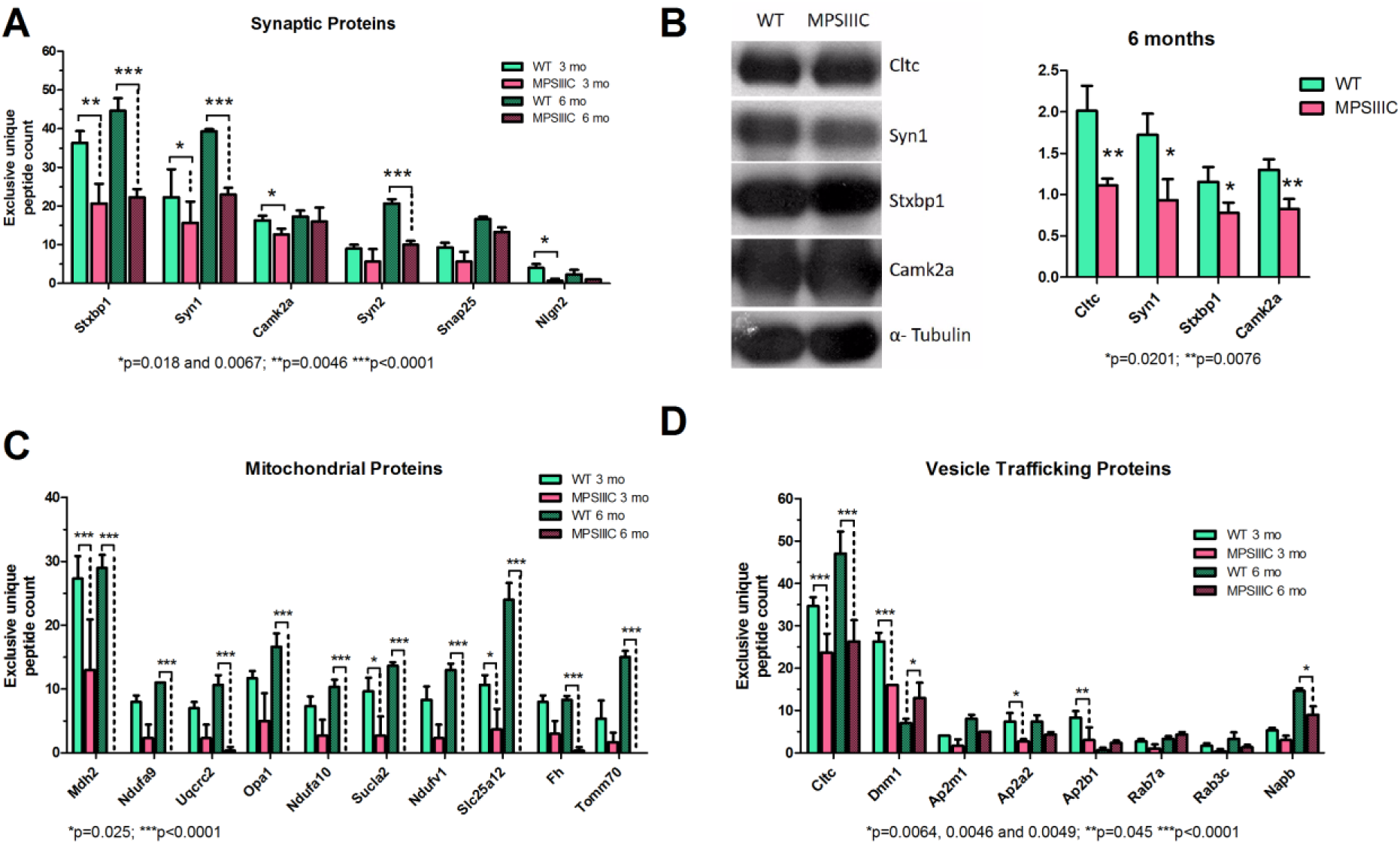
Semiquantitative LC-MS/MS analysis of proteins present in synaptosomes from brains of 3- and 6-month-old mice reveals deficiencies of synaptic, mitochondrial and trafficking vesicle-associated proteins in MPS IIIC mice. **(A)** Exclusive unique peptide counts of synaptic proteins. **(B)** Western blots of total protein extracts from brains of 6-month-old mice and their respective quantifications confirming changes in protein abundance identified by proteomic analysis. **(C)** Exclusive unique peptide counts of mitochondrial proteins. **(D)** Exclusive unique peptide counts of proteins associated with intracellular vesicle trafficking and endocytosis. Data obtained from synaptosomes extracted from 3 different animals per genotype. P-value was calculated in the exclusive unique peptide count and area of LC peptides with 2-way ANOVA using Bonferroni as a post-test.

The second group significantly changed in MPS IIIC neurons consisted of mitochondrial proteins and enzymes. Multiple proteins were reduced in MPS IIIC at the age of 3 months (Fig. 5C), with further drastic decrease by the age of 6 months when many of them were diminished to below the detection levels for LC-MS/MS technique. This confirmed progressive deficiency of mitochondrial function previously described by us for the neurons of MPS IIIC mice (13).

The third major group of proteins deficient or reduced in the synaptosomes from MPS IIIC mouse brains contained those involved in vesicle trafficking and endocytosis (Fig. 5D). In particular, we observed major alterations in the levels of clathrin heavy chain 1 (Cltc/CHC) and dynamin-1 (Dnm1). Cltc was reduced by 31.7% at 3 months (WT=34.7±2; MPS IIIC=23.7±4.5) and by 43.9% by 6 months (WT=47±5.1; MPS IIIC=26.3±5) in the MPS IIIC synaptosomes. Its reduction in the MPS IIIC mouse brain homogenates has been also confirmed by Western blot (Fig. 5B). Dnm1 was reduced by 39% at 3 months (WT=26.3±2; MPS IIIC=16±0), but increased by 46.1% at the age of 6 months (WT=7±1; MPS IIIC=13±3.6). Subunits of AP2a2 and AP2b1 of the adaptor protein complex 2 (AP-2) involved in recruiting clathrin for assembly of synaptic vesicles and selecting cargo for clathrin-mediated endocytosis(46) were also reduced in the MPSIIIC brain at the age of 3 months. The NSF attachment protein beta (Napb) was significantly reduced in MPS IIIC synaptosomes at the age of 6 months. Napb belongs to the group of SNAP proteins that play a role in SNARE complex dissociation and recycling (synaptic vesicle docking) and its deficiency has been associated with the emerging of seizures in human patients (47) and in mouse models (48). Together, the deficiency of proteins involved in vesicle trafficking and endocytosis suggested that these processes could be affected in the MPS IIIC neurons.

### 7. MPS IIIC hippocampal neurons show partial impairment of synaptic vesicle trafficking and turnover

Besides reduced density of synaptic vesicles and smaller areas/lengths of PSD, TEM analysis also identified defects of microtubules in the MPS IIIC cultured hippocampal neurons. The majority of microtubules were disorganized, sparse and non-parallel, with multiple storage bodies present between the microtubule filaments (Fig. 6A). This observation together with the reduction of proteins involved in vesicle targeting and turnover observed by proteomic analysis of synaptosomes suggested that the low density of synaptic vesicles at the axonal terminals could be caused by impaired vesicular trafficking of synaptic vesicle precursor organelles transported from the cell body. To test this hypothesis, we studied the axonal trafficking of synaptic vesicle precursors by live imaging in hippocampal neuronal cultures. On DIV 3 cells were transduced with LV encoding for Syn1-GFP (GeneCopoeia^TM^). On DIV 21, 10-min videos were recorded using a spinning disk microscope to visualize trafficking of GFP-positive vesicles (Fig. 6B). In MPS IIIC neurons, the majority of Syn1-GFP-positive vesicles remained stationary or wiggled back and forth instead of showing a constant motion in one direction (Fig. 6B, Supplementary video 2). Conversely, in WT neurons, the moving vesicles were progressing in a stable direction (Fig. 8B, Supplementary video 1) and their speed was faster as compared with that of vesicles in MPS IIIC cells (Fig. 6C). In addition, while majority of Syn1-GFP in WT neurons were associated with fine punctate distributed over the axons, in the MPS IIIC neurons they were localized in coarse intensely-stained spheroid granules (Fig. 6B). This type of staining was consistent with the hypothesis that Syn1-GFP is accumulated along the axon. These bodies, however, were negative for the lysosomal marker LAMP2, or autophagosomal marker LC3 suggesting that the inclusions containing Syn1-GFP were different from lysosomal storage bodies or autophagosomes (Fig. 6D).

**Figure 6.**
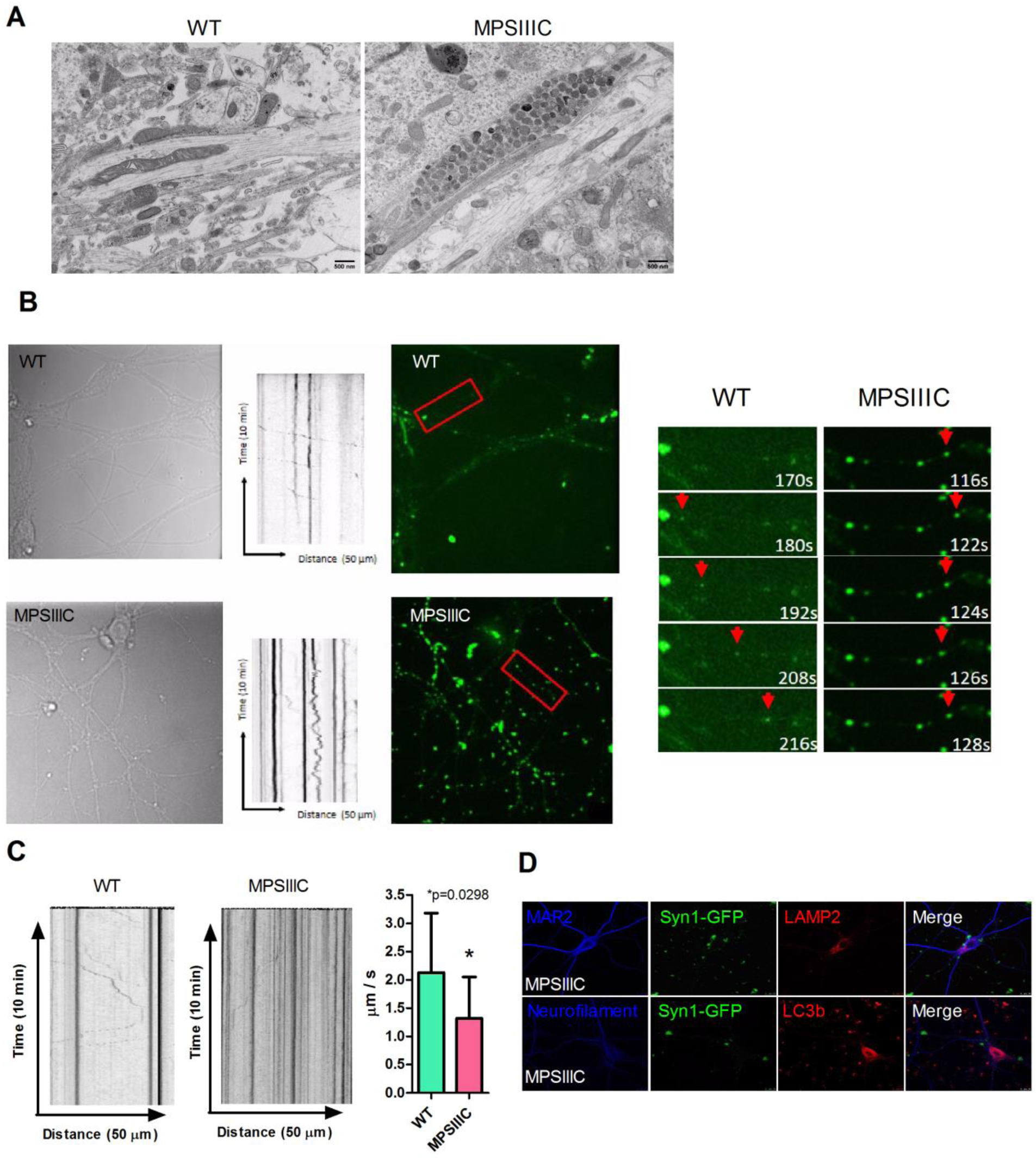
Vesicle transport defects in MPS IIIC neurons. **(A)** Microtubule network in neurites of MPS IIIC cultured hippocampal neurons is disorganized, sparse and non-parallel, and multiple electron-lucent and electron-dense storage vacuoles are present between the microtubule filaments. Panels show representative images of at least 30 from 3 different experiments or 3 different animals per genotype. **(B)** Live imaging of Syn1-GFP positive vesicles in cultured hippocampal neurons. Brightfield images, fluorescent images and kymographs of WT and MPS IIIC hippocampal neurons transduced with LV-Syn1-GFP. In MPS IIIC neurons, moving GFP-positive vesicles show a wiggling pattern, in contrast with the WT cells where the majority of moving vesicles travel in one direction. **(C)** Kymographs reveal that GFP-positive vesicles in the MPS IIIC neurons move at a slower speed than those in the WT cells. Videos were recorded for 10 min, with 1 recording every 2 s with a 63x objective. The bar graph shows the average speed on Syn1-positive vesicles and SD P-value was calculated by t-test. **(D)** The granules observed in the MPS IIIC neurons transduced with LV-Syn1-GFP do not co-localize with neither the lysosomal marker LAMP2, nor with autophagosomal marker LC3. For the quantification of vesicle velocity, 17 WT and 12 MPS IIIC cells originating from 3 different sets of experiments for each genotype were analyzed.

## Discussion

Neurodegeneration and neuroinflammation caused by metabolite storage, defects in autophagy and microgliosis were recognized as major causes of behavioural alterations and developmental delay in patients with neurological LSD. Our current data demonstrate, however, that both excitatory and inhibitory synaptic inputs to the pyramidal CA1 hippocampal neurons are drastically reduced in the mouse model of Sanfilippo C syndrome already at P45-60 i.e. at least 2-3 months before the development of other neuronal pathologies such as neuroinflammation, neuronal storage of gangliosides, misfolded proteins and mitochondrial damage (13). We observe that amplitudes and frequencies of both mEPSCs and mIPSC are reduced at P45-60. For comparison, in the mouse model of MPS IIIA slightly reduced mEPSCs amplitude but no changes in frequency was detected in the somatosensory cortex (23) while in the mouse model of Niemann-Pick C disease, neurons presented enhanced glutamatergic neurotransmission in hippocampus leading to hyperexcitability (49). Our electrophysiological data closely parallel the marked reduction of the vGlut1/PSD-95 colocalization as observed in cultured MPS IIIC neurones and in the hippocampal neurones of 6-month-old MPS III mice, together suggesting an overall synaptic deficit that aggravates with age. It is interesting to note that, we do not observe a similarly reduced vGAT/Gephyrin colocalization in cultures as we would have expected to see based on our electrophysiological observations. We, thus, speculate that a reduced GABAergic input results from a homeostatic compensatory mechanism triggered by an intrinsic glutamatergic deficit. Moreover, density of dendritic synaptic spines of pyramidal CA1 hippocampal neurons is reduced already at P10 and never reaches the levels observed in the WT mice. Drastically reduced levels of synaptic vesicles in the terminals and smaller areas of postsynaptic densities were found both in pyramidal CA1 hippocampal neurons at 3 and 6 months of age and in cultured hippocampal neurons at DIV 21. These changes affect mainly excitatory circuits, and PSD-95 seems to be one of the most severely reduced biomarkers not only in the mouse model but also in all studied post-mortem cortical tissues of neurological MPS human patients. Importantly, the levels of this protein can be rescued *in vivo* and *in vitro* by correcting the primary genetic defect in neurons with the viral vectors expressing the WT human HGSNAT enzyme. This implicates the genetic defects in HGSNAT as a primary cause of synaptic deficits but also suggests that improvements in the behaviour of MPSIIIC mice treated by striatal administration of AAV-HGSNAT vectors (31) could be at least partially explained by the rescue of synaptic defects.

Major changes in the synaptic morphology and the levels and distribution of synaptic markers found in the hippocampus i.e. scarcity of synaptic vesicles in the excitatory terminals, reduced PSD length, reduced density of mature mushroom-shaped spines were mainly recapitulated in dissociated neuronal cultures. In contrast, the alterations in excitatory or inhibitory neurotransmission had different features *in vivo* and *in vitro*. Miniature synaptic currents from hippocampal cultured neurons revealed no significant changes in amplitude or frequency of mIPSCs whereas an almost 3-fold increased frequency was detected in mEPSCs. In contrast, our acute hippocampal slice recordings showed reduction of both frequency and amplitude of mEPSCs and mIPSCs, suggesting that both excitatory and inhibitory neurotransmission are deficient in MPSIIIC mice. Previously reported studies of pluripotent stem cells-derived neurons generated from human MPS IIIC patients’ fibroblasts also revealed reduced amplitudes of firing, defects in neuronal activity, network-wide degradation and altered effective connectivity (50). It is tempting to speculate that changes in GABAergic synaptic transmission occurring *in vivo* but not in neuronal cultures, appear due to an attempt to compensate defects in excitatory transmission. Furthermore, the change in frequency and lack of a significant change in the kinetics of the events suggests a possible presynaptic locus of deficit. This latter observation is also in agreement with reduced density of synaptic vesicles in the axonal terminals in MPS IIIC mice as compared with WT.

We hypothesize that the increase in mEPSCs observed in cultured MPS IIIC neurons is linked to the major changes in the positioning of synapses on dendrites detected by both confocal and transmission electron microscopy. While the overwhelming majority of synapses in WT cultures were axospinous, in MPS IIIC neurons >75% of the synapses were axodendritic. Since MPS IIIC cultured neuron dendrites present mainly immature filipodia spines that lack the proper postsynaptic machinery for synaptic function (51), we speculate that they reallocate their excitatory synapses to the dendrites, altering their response to the presynaptic signaling which could potentially explain increased frequency and amplitude of mEPSC observed in neuronal cultures. Such changes have not been observed in the mouse brains consistent with the reduction of both amplitude and frequency of miniature postsynaptic currents.

The disruption of the microtubule architecture in the axons of cultured MPS IIIC neurons (disorganized, sparse and non-parallel filaments) together with reduced levels of proteins involved in vesicle trafficking and endocytosis in the synaptosomes from MPS IIIC mouse brains allowed us to hypothesize that the scarcity of synaptic vesicles in the axonal terminals can be caused by partially impaired trafficking of synaptic vesicle precursors from the soma. Indeed, direct analysis of the axonal movement of Syn1 positive vesicles by live imaging microscopy demonstrated that in MPS IIIC neurons fewer vesicles were moving in a stable direction towards the terminal. Besides, vesicles that were moving in one direction had slower speed that that of vesicles in WT neurons. In addition, the majority of newly synthesized Syn1-GFP in MPS IIIC neurons was localized in coarse spheroid granules distributed along the axon. Formation of focal granular enlargements within axons (axonal spheroids or “torpedoes”; neuroaxonal dystrophy) and dendrites is a phenomenon described in a variety of lysosomal storage diseases including mannosidosis, GM1and GM2-gangliosidosis, prosaposin deficiency, and Niemann-Pick type C (20, 21, 52–55). In the axons spheroids often contained electron-dense concentric lamellar bodies and neurofilaments (55).

The mechanism underlying formation of the axonal spheroids in LSD is not completely understood. They could be formed by merging of lysosomes in the proximal axon since accumulation of storage materials in lysosomes impairs their onward BORC-dependent axonal transport (56). Besides, the block in anterograde transport of lysosomes prevents their merging with early endosomes and autophagosomes in the axon terminal (57). As a result, the dynein-mediated retrograde transport of autophagosomes and their maturation to autolysosomes (58) is impaired causing their accumulation in distal axon and formation of axonal spheroids. Our data, however, show that the lysosomal storage bodies are primarily found in the soma, and axonal Syn1-positive spheroid granules are negative for the markers of endo-lysosomal pathway or for the autophagosomal marker LC3. We speculate, therefore, that they appear in the process of accumulation of the synaptic vesicle precursors at the places in the axons where the transport is blocked due to the microtubule defects.

## Conclusions

Together, our experiments demonstrate that lysosomal storage in MPS IIIC neurons results in appearance of early and drastic defects in the synapse of pyramidal CA1 neurons. Further studies are necessary to define the exact mechanism, however based on our current results we speculate that the defects are caused by the appearance of lysosomal storage bodies that disrupt microtubule structure and normal transport of vesicles carrying the synaptic proteins along the axon (59). This, in concert with impaired autophagy and endocytosis at the axonal terminals, could result in the deficit of synaptic vesicles affecting synaptic transmission and plasticity. Reduction of PSD-95-positive puncta was detected in post-mortem cortices of human MPS I, II, IIIA, C and D patients indicating that they manifest with similar excitatory synaptic defects. Moreover, the density of synaptic spines is also reduced in mouse models of sialidosis and Tay-Sachs disease suggesting that similar pathogenic pathways can also exist in other lysosomal diseases. In the light of these findings it is tempting to speculate that drugs known to enhance excitatory synaptic transmission should be tested for their ability to improve synaptic function as well as behavioral and cognitive defects in MPS IIIC mice and in the animal models of other neurological LSD.

## Supporting information

Supplementary video 2 MPSIIIC

## List of abbreviations

ACSF: artificial cerebrospinal fluid
AMPA: α-amino-3-hydroxy-5-methyl-4-isoxazolepropionic acid
AP-2: adaptor protein complex
BMI: bicuculline methiodide
BSA: bovine serum albumin
CaMKII: calcium-calmodulin kinase II
CCAC: Canadian Council on Animal Care
Cltc/CHC: clathrin heavy chain 1
CNS: central nervous system
CSPα: cysteine string protein α
CMV: cytomegalovirus
DAPI: 4′,6-diamidino-2-phenylindole
Dil: 1,1’-Dioctadecyl-3,3,3’,3’-Tetramethylindocarbocyanine Perchlorate
DMEM: Dulbecco’s Modified Eagle’s Medium
Dnm1: dynamin-1
DIV: day *in vitro*
DNQX: 6,7-dinitroquinoxaline-2,3-dione
EGFP: enhanced green fluorescent protein
ERT: received enzyme replacement therapy
FA: formic acid
FBS: fetal bovine serum
FDR: false-discovery rate
GABA: gamma aminobutyric acid
GFP: green fluorescent protein
GO: gene ontology
HBSS: Hank’s Balanced Salt Solution
HS: heparan sulfate
HSCT: hematopoietic stem cell transplantation
LAMP1: lysosomal-associated membrane protein 1
LC-MS/MS: liquid chromatography tandem mass spectrometry
LSD: lysosomal storage disease
LV: lentivirus
MAP2: microtubule-associated protein 2
mEPSC: miniature excitatory postsynaptic current
mIPSC: miniature inhibitory postsynaptic current
MOI: multiplicity of infection
MPS: mucopolysaccharidosis
Napb: NSF attachment protein beta
NF-M: neurofilament medium chain
Nlgn1: neuroligin-1
Nlgn2: neuroligin-2
NMDAR: N-methyl-D-aspartate receptor
OCT: optimum cutting temperature
PEI: Polyethylenimine
PFA: paraformaldehyde
PSD: postsynaptic density
PSD-95: postsynaptic density protein 95
SNARE: soluble NSF attachment receptor
Stxbp1/Munc18-1: syntaxin-binding protein 1
Syn1: synapsin 1
Syn2: synapsin 2
TCEP: tris(2-carboxyethyl) phosphine hydrochloride
TEM: transmission electron microscopy
TFA: trifluoroacetic acid
TT: true type
TTX: tetrodotoxin

## Declarations

### Ethics approval and consent to participate

All the experiments performed on mice have been approved by the Animal Care and User Committee of the Ste-Justine Hospital Research Center. Ethical approval for the research involving human tissues was granted by Research Ethics Board (Comité d’éthique de la recherche) of CHU Ste-Justine. Frozen or fixed with PFA, cerebral cortices from clinically confirmed MPS patients (1 case of MPS I, 1 case of MPS II, 2 cases of MPS IIIA, 1 case of MPS IIIC and 2 cases of MPS IIID) and age-matched controls with no pathological changes in the central nervous system were provided by NIH NeuroBioBank (project 1071, MPS Synapse) together with clinical descriptions and results of the neuropathological examination. Families provided the informed consent prior to donation of tissues.

### Consent for publication

Not applicable.

### Availability of data and materials

All data generated or analyzed during this study are included in this published article and its supplementary information files.

### Authors’ contributions

Conducted experiments and acquired data: C.B.P.A., L.B., P.B., X.P., C.H., E.F., E.B., C.O.; analyzed data: C.B.P.A., L.B., P.B., X.P., C.H., E.F., E.B., C.O., P.S.M., J.-C.L., C.R.M., G.D. and A.V.P.; provided reagents: B.B., wrote the manuscript (first draft) C.B.P.A., P.B., E.F., A.V.P.; wrote the manuscript (editing) A.V.P., J.-C.L., G.D., P.S.M., P.T., B.B., C.R.M. All authors read and approved the final manuscript.

## Acknowledgements

The authors thank Dr. Christian Beausejour and Gaël Moquin-Beaudry for the help in production of LV-HGSNAT-GFP. We also thank Jeannie Mui and the Facility for Electron Microscopy Research (FEMR – McGill University) for the help with the transmission electron microscopy, Dr. Elke Küster-Schöck and the Plateforme d’Imagerie Microscopique (PIM – CHU Sainte Justine) for the help with life imaging microscopy and Dr. Mila Ashmarina for critically reading the manuscript and helpful advice.

## SUPPLEMENTARY FIGURES

**Figure S1:**
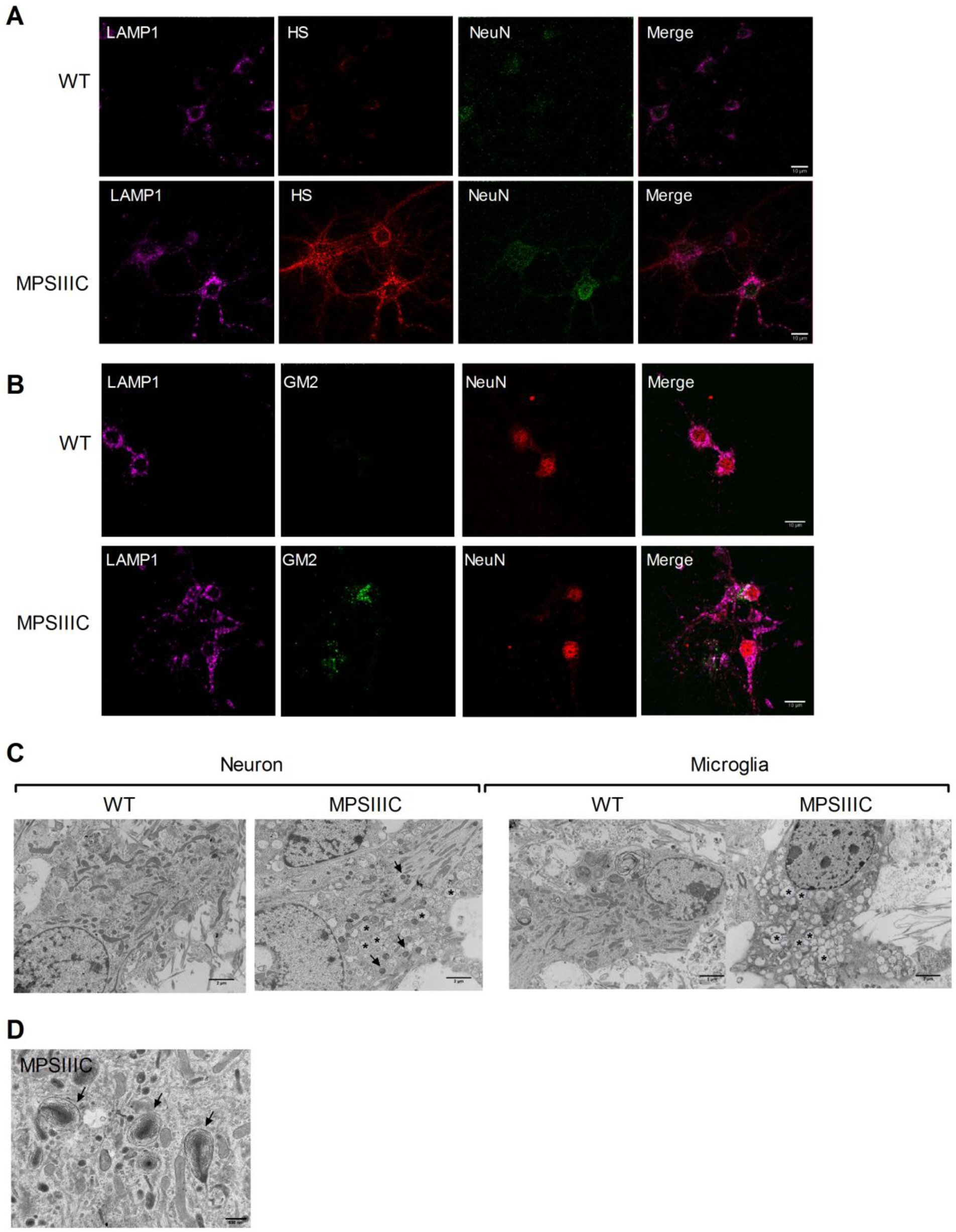
Primary and secondary storage in cultured hippocampal neurons from MPSIIIC mice. **(A)** Cultured primary hippocampal neurons from MPS IIIC mice at DIV 21 contain multiple course HS-positive/LAMP1-positive cytoplasmic puncta, consistent with the lysosomal storage of HS. **(B)** Cultured hippocampal MPSIIIC neurons also show storage of GM2 ganglioside in the granules only partially co-localizing with LAMP1-positive vacuoles. **(C, D)** The dual pattern of storage is confirmed by electron microscopy, where, both, electron-dense storage bodies (arrows) containing lipids and misfolded proteins and electron-lucent organelles (*) with glycan storage can be observed in both cultured neurons and microglia derived from the brains of MPSIIIC embryos. Scale bar is 10 µm in **(A)** and **(B)**, 2 µm in **(C)** and 500 nm in **(D)**. Panels show representative images from 3 independent sets of experiments.

**Figure S2:**
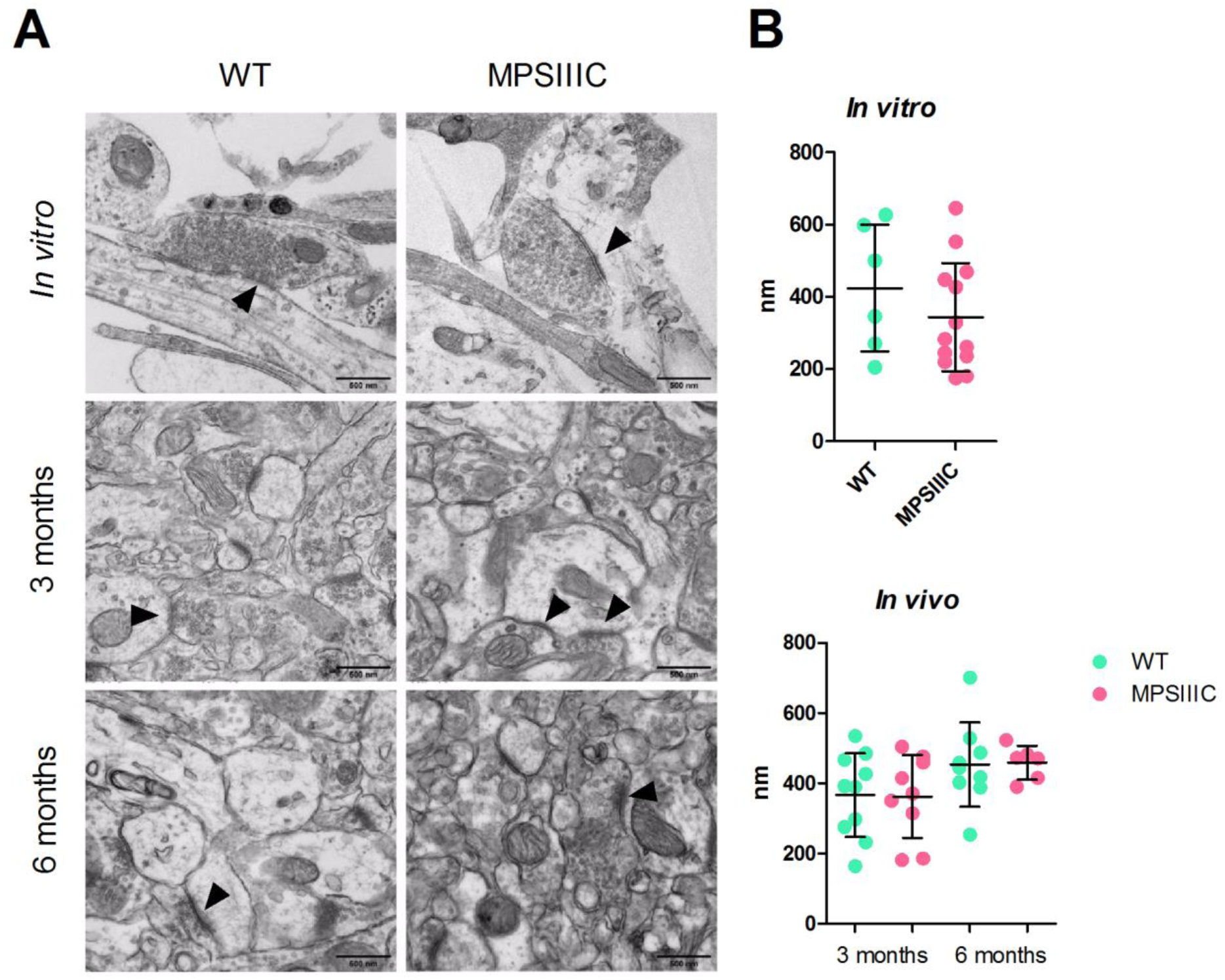
Lengths of PSD in symmetrical inhibitory synapses are similar between MPSIIIC and WT neurons *in vivo* and *in vitro*. **(A)** Electron micrographs of inhibitory symmetric PSD (arrowheads) in cultured neurons and in the CA1 region of the hippocampus from 3- and 6-month-old mice. **(B)** Quantification of length of PSD *in vitro* and *in vivo*. Graphs show mean and SD values from 3 independent sets of experiments. Panels show representative images from 3 independent sets of experiments.

**Figure S3:**
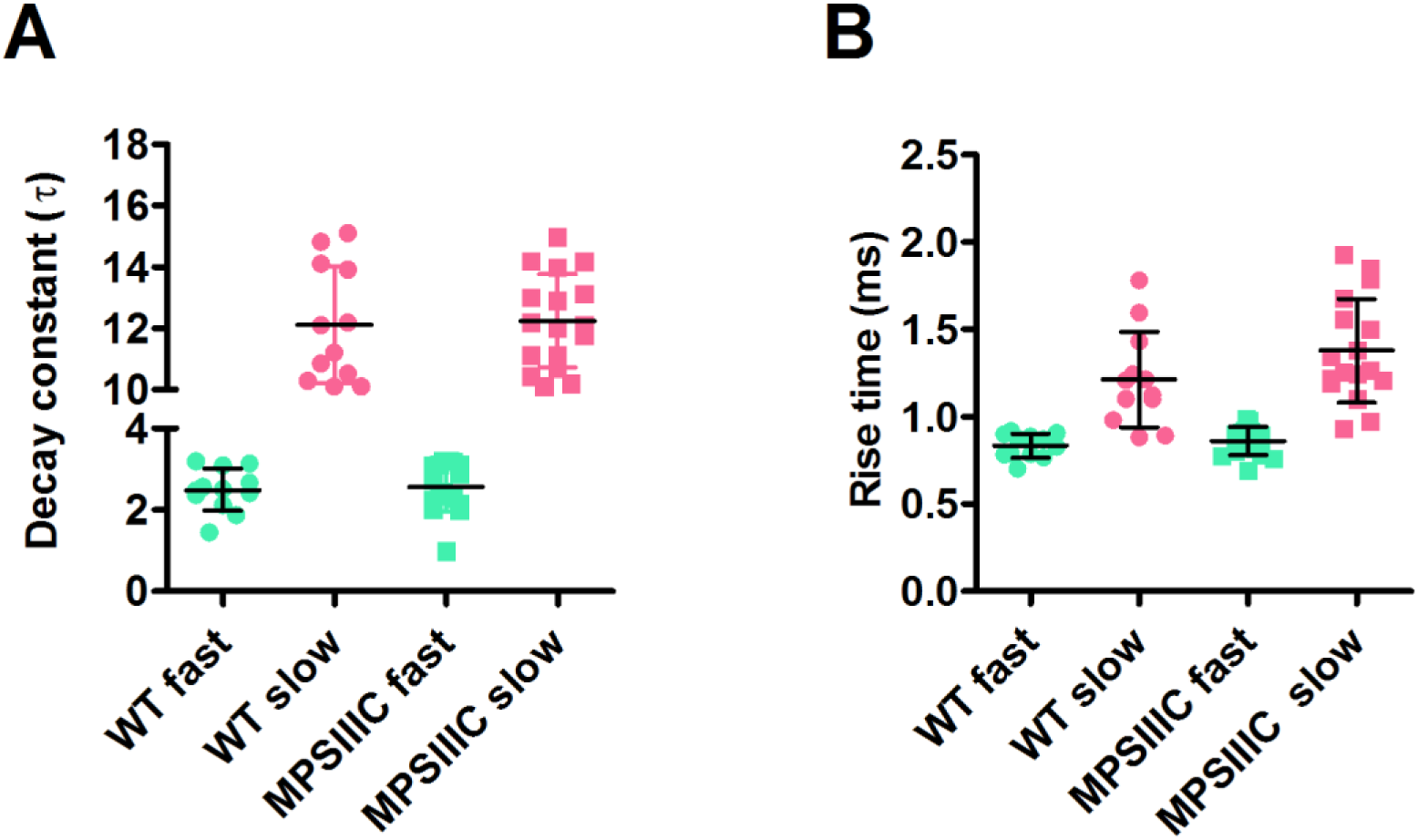
**(A)** No significant differences in the slow or fast decay constant of mIPSCs. Average mIPSC kinetics: τ_slow_: MPS IIIC= 15.2±0.87; WT=14.82±0.56; τ_fast_: MPS IIIC= 2.54±0.29; WT=2.48±0.26). **(B)** No significant differences in the slow or fast rise time of mIPSCs between WT and MPS IIIC animals. Average mIPSC kinetics: rise time _slow_: MPS IIIC= 1.54±0.04; WT=1.422±0.06; rise time _fast_: MPS IIIC= 0.74±0.29; WT=0.88±0.26. n=12 for WT and n=17 for MPS IIIC.

**Figure S4:**
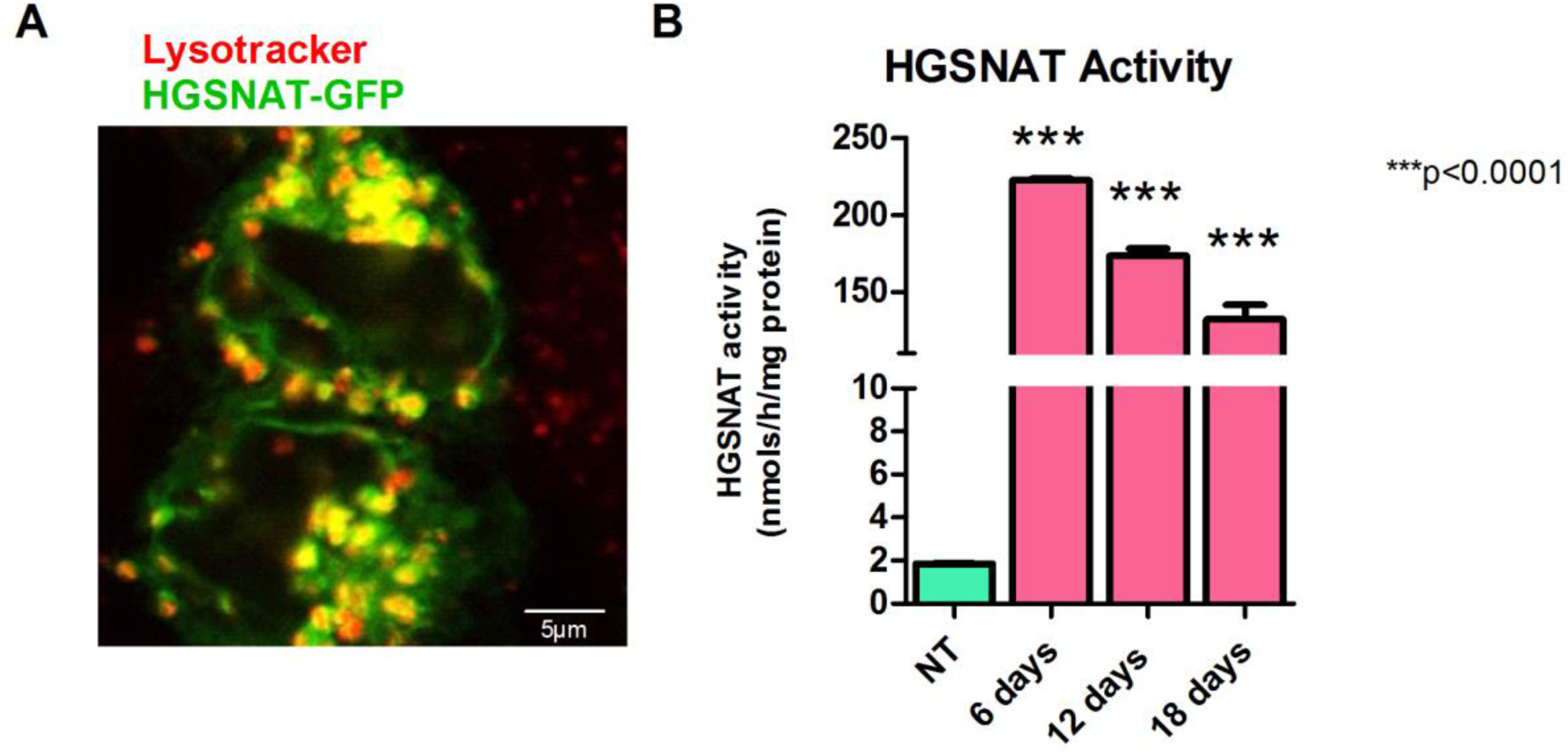
Transduction of HEK 293 cells with LV-HGSNAT-GFP results in expression of enzymatically active and correctly targeted HGSNAT-GFP fusion protein. **(A)** Fluorescent confocal images of HEK 293 cells transduced with LV-HGSNAT-GFP. Colocalization of GFP puncta and Lysotracker red staining demonstrates that HGSNAT-GFP fusion protein is correctly targeted to the lysosomes. **(B)** HGSNAT activity measured in homogenates of HEK293 cells transduced with LV-HGSNAT-GFP is >100-fold increased as compared with the cells transduced with LV-GFP.

**Figure S5:**
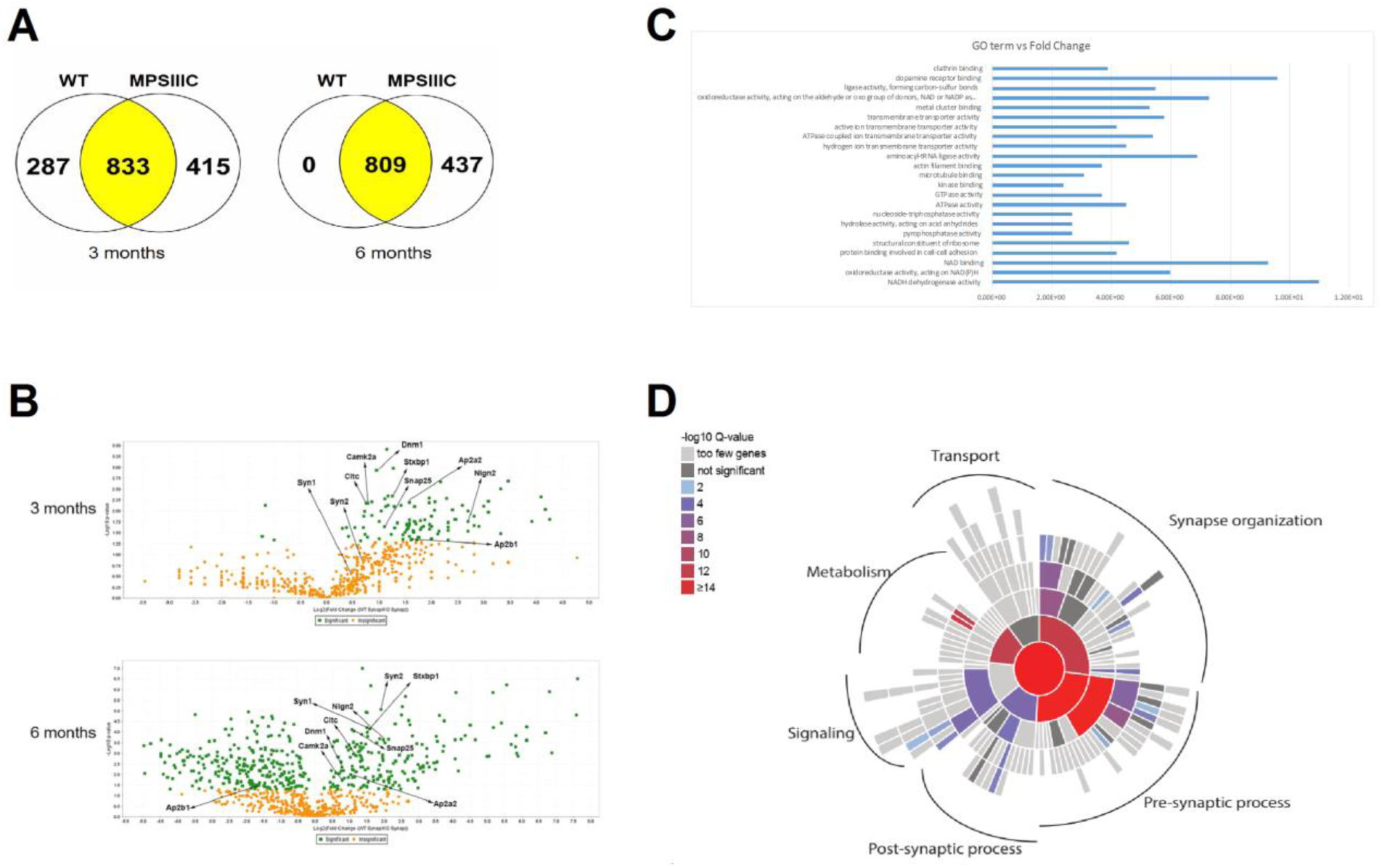
Proteomic statistics. **(A)** Total number of proteins identified by LC/MS in synaptosomes extracted from the brains of WT and MPSIIIC mice at 3 and 6 months of age. **(B)** Volcano plots of the proteins identified in synaptosomes showing the proteins that are statistically different between WT and MPS IIIC. **(C)** Gene ontology terms (GO terms) versus fold-change of the proteins reduced in MPS IIIC synaptosomes. The data represent values where the Benjamini corrected p-value is below the highest Benjamini corrected p-value for the GO terms. **(D)** q values per ontology term visualized for biological process enrichment.

## REFERENCES

1. Boustany RM. Lysosomal storage diseases--the horizon expands. Nat Rev Neurol. 2013;9(10):583–98.

2. Meikle PJ, Hopwood JJ, Clague AE, Carey WF. Prevalence of lysosomal storage disorders. JAMA. 1999;281(3):249–54.

3. Bartsocas C, Grobe H, van de Kamp JJ, von Figura K, Kresse H, Klein U, et al. Sanfilippo type C disease: clinical findings in four patients with a new variant of mucopolysaccharidosis III. Eur J Pediatr. 1979;130(4):251–8.

4. Scriver CR, Beaudet AL, Sly WS, Valle D, Stanbury JB, Wyngaarden JB, et al. The metabolic and molecular bases of inherited disease. New York: McGraw-Hill, Health Professions Division; 2001.

5. Valstar MJ, Ruijter GJ, van Diggelen OP, Poorthuis BJ, Wijburg FA. Sanfilippo syndrome: a mini-review. J Inherit Metab Dis. 2008;31(2):240–52.

6. Ruijter GJ, Valstar MJ, van de Kamp JM, van der Helm RM, Durand S, van Diggelen OP, et al. Clinical and genetic spectrum of Sanfilippo type C (MPS IIIC) disease in The Netherlands. Mol Genet Metab. 2008;93(2):104–11.

7. Berger-Plantinga EG, Vanneste JA, Groener JE, van Schooneveld MJ. Adult-onset dementia and retinitis pigmentosa due to mucopolysaccharidosis III-C in two sisters. J Neurol. 2004;251(4):479–81.

8. Poupetova H, Ledvinova J, Berna L, Dvorakova L, Kozich V, Elleder M. The birth prevalence of lysosomal storage disorders in the Czech Republic: comparison with data in different populations. J Inherit Metab Dis. 2010;33(4):387–96.

9. Kresse H. Mucopolysaccharidosis 3 A (Sanfilippo A disease): deficiency of a heparin sulfamidase in skin fibroblasts and leucocytes. Biochem Biophys Res Commun. 1973;54(3):1111–8.

10. O’Brien JS. Sanfilippo syndrome: profound deficiency of alpha-acetylglucosaminidase activity in organs and skin fibroblasts from type-B patients. Proc Natl Acad Sci U S A. 1972;69(7):1720–2.

11. Klein U, Kresse H, von Figura K. Sanfilippo syndrome type C: deficiency of acetyl-CoA:alpha-glucosaminide N-acetyltransferase in skin fibroblasts. Proc Natl Acad Sci U S A. 1978;75(10):5185–9.

12. Kresse H, Paschke E, von Figura K, Gilberg W, Fuchs W. Sanfilippo disease type D: deficiency of N-acetylglucosamine-6-sulfate sulfatase required for heparan sulfate degradation. Proc Natl Acad Sci U S A. 1980;77(11):6822–6.

13. Martins C, Hulkova H, Dridi L, Dormoy-Raclet V, Grigoryeva L, Choi Y, et al. Neuroinflammation, mitochondrial defects and neurodegeneration in mucopolysaccharidosis III type C mouse model. Brain. 2015;138(Pt 2):336–55.

14. Shapiro EG, Nestrasil I, Delaney KA, Rudser K, Kovac V, Nair N, et al. A Prospective Natural History Study of Mucopolysaccharidosis Type IIIA. J Pediatr. 2016;170:278–87 e1-4.

15. Whitley CB, Cleary M, Eugen Mengel K, Harmatz P, Shapiro E, Nestrasil I, et al. Observational Prospective Natural History of Patients with Sanfilippo Syndrome Type B. J Pediatr. 2018;197:198–206 e2.

16. Shapiro E, Ahmed A, Whitley C, Delaney K. Observing the advanced disease course in mucopolysaccharidosis, type IIIA; a case series. Mol Genet Metab. 2018;123(2):123–6.

17. Barone R, Nigro F, Triulzi F, Musumeci S, Fiumara A, Pavone L. Clinical and neuroradiological follow-up in mucopolysaccharidosis type III (Sanfilippo syndrome). Neuropediatrics. 1999;30(5):270–4.

18. Zafeiriou DI, Savvopoulou-Augoustidou PA, Sewell A, Papadopoulou F, Badouraki M, Vargiami E, et al. Serial magnetic resonance imaging findings in mucopolysaccharidosis IIIB (Sanfilippo’s syndrome B). Brain Dev. 2001;23(6):385–9.

19. Karabelas AB, Walkley SU. Altered patterns of evoked synaptic activity in cortical pyramidal neurons in feline ganglioside storage disease. Brain Res. 1985;339(2):329–36.

20. Purpura DP, Highstein SM, Karabelas AB, Walkley SU. Intracellular recording and HRP-staining of cortical neurons in feline ganglioside storage disease. Brain Res. 1980;181(2):446–9.

21. Pressey SN, Smith DA, Wong AM, Platt FM, Cooper JD. Early glial activation, synaptic changes and axonal pathology in the thalamocortical system of Niemann-Pick type C1 mice. Neurobiol Dis. 2012;45(3):1086–100.

22. Teixeira CA, Miranda CO, Sousa VF, Santos TE, Malheiro AR, Solomon M, et al. Early axonal loss accompanied by impaired endocytosis, abnormal axonal transport, and decreased microtubule stability occur in the model of Krabbe’s disease. Neurobiol Dis. 2014;66:92–103.

23. Dwyer CA, Scudder SL, Lin Y, Dozier LE, Phan D, Allen NJ, et al. Neurodevelopmental Changes in Excitatory Synaptic Structure and Function in the Cerebral Cortex of Sanfilippo Syndrome IIIA Mice. Sci Rep. 2017;7:46576.

24. Sambri I, D’Alessio R, Ezhova Y, Giuliano T, Sorrentino NC, Cacace V, et al. Lysosomal dysfunction disrupts presynaptic maintenance and restoration of presynaptic function prevents neurodegeneration in lysosomal storage diseases. EMBO Mol Med. 2017;9(1):112–32.

25. Crawley AC, Gliddon BL, Auclair D, Brodie SL, Hirte C, King BM, et al. Characterization of a C57BL/6 congenic mouse strain of mucopolysaccharidosis type IIIA. Brain Res. 2006;1104(1):1–17.

26. Hansen GM, Markesich DC, Burnett MB, Zhu Q, Dionne KM, Richter LJ, et al. Large-scale gene trapping in C57BL/6N mouse embryonic stem cells. Genome Res. 2008;18(10):1670–9.

27. Phaneuf D, Wakamatsu N, Huang JQ, Borowski A, Peterson AC, Fortunato SR, et al. Dramatically different phenotypes in mouse models of human Tay-Sachs and Sandhoff diseases. Hum Mol Genet. 1996;5(1):1–14.

28. Pan X, De Aragao CBP, Velasco-Martin JP, Priestman DA, Wu HY, Takahashi K, et al. Neuraminidases 3 and 4 regulate neuronal function by catabolizing brain gangliosides. FASEB J. 2017;31(8):3467–83.

29. Feng G, Mellor RH, Bernstein M, Keller-Peck C, Nguyen QT, Wallace M, et al. Imaging neuronal subsets in transgenic mice expressing multiple spectral variants of GFP. Neuron. 2000;28(1):41–51.

30. Croce A, Pelletier JG, Tartas M, Lacaille JC. Afferent-specific properties of interneuron synapses underlie selective long-term regulation of feedback inhibitory circuits in CA1 hippocampus. J Physiol. 2010;588(Pt 12):2091–107.

31. Tordo J, O’Leary C, Antunes A, Palomar N, Aldrin-Kirk P, Basche M, et al. A novel adeno-associated virus capsid with enhanced neurotropism corrects a lysosomal transmembrane enzyme deficiency. Brain. 2018.

32. Campeau E, Ruhl VE, Rodier F, Smith CL, Rahmberg BL, Fuss JO, et al. A versatile viral system for expression and depletion of proteins in mammalian cells. PLoS One. 2009;4(8):e6529.

33. Toni N, Buchs PA, Nikonenko I, Bron CR, Muller D. LTP promotes formation of multiple spine synapses between a single axon terminal and a dendrite. Nature. 1999;402(6760):421–5.

34. Jeyakumar M, Thomas R, Elliot-Smith E, Smith DA, van der Spoel AC, d’Azzo A, et al. Central nervous system inflammation is a hallmark of pathogenesis in mouse models of GM1 and GM2 gangliosidosis. Brain. 2003;126(Pt 4):974–87.

35. Lowden JA, O’Brien JS. Sialidosis: a review of human neuraminidase deficiency. Am J Hum Genet. 1979;31(1):1–18.

36. Pshezhetsky AV, Richard C, Michaud L, Igdoura S, Wang S, Elsliger MA, et al. Cloning, expression and chromosomal mapping of human lysosomal sialidase and characterization of mutations in sialidosis. Nat Genet. 1997;15(3):316–20.

37. Stavoe AKH, Holzbaur ELF. Axonal autophagy: Mini-review for autophagy in the CNS. Neurosci Lett. 2019;697:17–23.

38. Hunt CA, Schenker LJ, Kennedy MB. PSD-95 is associated with the postsynaptic density and not with the presynaptic membrane at forebrain synapses. J Neurosci. 1996;16(4):1380–8.

39. Song JY, Ichtchenko K, Sudhof TC, Brose N. Neuroligin 1 is a postsynaptic cell-adhesion molecule of excitatory synapses. Proc Natl Acad Sci U S A. 1999;96(3):1100–5.

40. Sudhof TC. Neuroligins and neurexins link synaptic function to cognitive disease. Nature. 2008;455(7215):903–11.

41. Capogna M, Pearce RA. GABA A,slow: causes and consequences. Trends Neurosci. 2011;34(2):101–12.

42. Banks MI, White JA, Pearce RA. Interactions between distinct GABA(A) circuits in hippocampus. Neuron. 2000;25(2):449–57.

43. Whittaker VP, Michaelson IA, Kirkland RJ. The separation of synaptic vesicles from nerve-ending particles (’synaptosomes’). Biochem J. 1964;90(2):293–303.

44. Huang da W, Sherman BT, Lempicki RA. Bioinformatics enrichment tools: paths toward the comprehensive functional analysis of large gene lists. Nucleic Acids Res. 2009;37(1):1–13.

45. Varoqueaux F, Jamain S, Brose N. Neuroligin 2 is exclusively localized to inhibitory synapses. Eur J Cell Biol. 2004;83(9):449–56.

46. Robinson MS. Adaptable adaptors for coated vesicles. Trends Cell Biol. 2004;14(4):167–74.

47. Conroy J, Allen NM, Gorman KM, Shahwan A, Ennis S, Lynch SA, et al. NAPB - a novel SNARE-associated protein for early-onset epileptic encephalopathy. Clin Genet. 2016;89(2):E1–3.

48. Burgalossi A, Jung S, Meyer G, Jockusch WJ, Jahn O, Taschenberger H, et al. SNARE protein recycling by alphaSNAP and betaSNAP supports synaptic vesicle priming. Neuron. 2010;68(3):473–87.

49. D’Arcangelo G, Grossi D, De Chiara G, de Stefano MC, Cortese G, Citro G, et al. Glutamatergic neurotransmission in a mouse model of Niemann-Pick type C disease. Brain Res. 2011;1396:11–9.

50. Canals I, Soriano J, Orlandi JG, Torrent R, Richaud-Patin Y, Jimenez-Delgado S, et al. Activity and High-Order Effective Connectivity Alterations in Sanfilippo C Patient-Specific Neuronal Networks. Stem Cell Reports. 2015;5(4):546–57.

51. Sekino Y, Kojima N, Shirao T. Role of actin cytoskeleton in dendritic spine morphogenesis. Neurochem Int. 2007;51(2-4):92–104.

52. March PA, Thrall MA, Brown DE, Mitchell TW, Lowenthal AC, Walkley SU. GABAergic neuroaxonal dystrophy and other cytopathological alterations in feline Niemann-Pick disease type C. Acta Neuropathol. 1997;94(2):164–72.

53. Walkley SU, Baker HJ, Rattazzi MC, Haskins ME, Wu JY. Neuroaxonal dystrophy in neuronal storage disorders: evidence for major GABAergic neuron involvement. J Neurol Sci. 1991;104(1):1–8.

54. Walkley SU, Blakemore WF, Purpura DP. Alterations in neuron morphology in feline mannosidosis. A Golgi study. Acta Neuropathol. 1981;53(1):75–9.

55. Oya Y, Nakayasu H, Fujita N, Suzuki K, Suzuki K. Pathological study of mice with total deficiency of sphingolipid activator proteins (SAP knockout mice). Acta Neuropathol. 1998;96(1):29–40.

56. Farias GG, Guardia CM, De Pace R, Britt DJ, Bonifacino JS. BORC/kinesin-1 ensemble drives polarized transport of lysosomes into the axon. Proc Natl Acad Sci U S A. 2017;114(14):E2955–E64.

57. Maday S. Mechanisms of neuronal homeostasis: Autophagy in the axon. Brain Res. 2016;1649(Pt B):143–50.

58. Cheng XT, Zhou B, Lin MY, Cai Q, Sheng ZH. Axonal autophagosomes recruit dynein for retrograde transport through fusion with late endosomes. J Cell Biol. 2015;209(3):377–86.

59. Jeyakumar M, Dwek RA, Butters TD, Platt FM. Storage solutions: treating lysosomal disorders of the brain. Nat Rev Neurosci. 2005;6(9):713–25.

