## Supplementary Table S2 for "Early defects in lysosomal storage diseases disrupt excitatory synaptic transmission"

**Supplementary table S2 : List of reduced proteins in MPSIIIC synaptosomes of 3 and 6 month-old mice identified by LC/MS.**

**List of reduced proteins in synaptosomes from brains of 3 month-old mice**

| # | Identified Proteins | Accession Number | Alternate ID | Molecular Weight | T-Test (p-value): (p < 0.05) | Quantitative Profile | KO S1 | KO S2 | KO S3 | WT S1 | WT S2 | WT S3 |
|---|---|---|---|---|---|---|---|---|---|---|---|---|
| 5 | Sodium/potassium-transporting ATPase subunit alpha OS=Mus musculus GN=Atp1a3 PE=1 SV=1 | A0A0G2JGX4_MOUSE (+1) | Atp1a3 | 113 kDa | 0.013 | KO Synap low, WT Synap high | 27 | 30 | 37 | 46 | 50 | 50 |
| 6 | Clathrin heavy chain 1 OS=Mus musculus GN=Cltc PE=1 SV=3 | CLH1_MOUSE (+1) | Cltc | 192 kDa | 0.0066 | KO Synap low, WT Synap high | 19 | 24 | 28 | 33 | 37 | 34 |
| 7 | Excitatory amino acid transporter 2 OS=Mus musculus GN=Slc1a2 PE=1 SV=1 | EAA2_MOUSE | Slc1a2 | 62 kDa | 0.024 | KO Synap low, WT Synap high | 9 | 9 | 13 | 13 | 13 | 14 |
| 8 | Syntaxin-binding protein 1 OS=Mus musculus GN=Stxbp1 PE=1 SV=2 | STXB1_MOUSE | Stxbp1 | 68 kDa | 0.0046 | KO Synap low, WT Synap high | 15 | 22 | 25 | 33 | 37 | 39 |
| 9 | Sodium/potassium-transporting ATPase subunit alpha-2 OS=Mus musculus GN=Atp1a2 PE=1 SV=1 | AT1A2_MOUSE (+1) | Atp1a2 | 112 kDa | 0.049 | KO Synap low, WT Synap high | 12 | 15 | 18 | 23 | 27 | 22 |
| 10 | Fructose-bisphosphate aldolase C OS=Mus musculus GN=Aldoc PE=1 SV=4 | ALDOC_MOUSE | Aldoc | 39 kDa | 0.01 | KO Synap low, WT Synap high | 11 | 10 | 8 | 13 | 15 | 11 |
| 11 | Fructose-bisphosphate aldolase A OS=Mus musculus GN=Aldoa PE=1 SV=2 | ALDOA_MOUSE | Aldoa | 39 kDa | 0.035 | KO Synap low, WT Synap high | 16 | 9 | 16 | 18 | 24 | 18 |
| 12 | Malate dehydrogenase, mitochondrial OS=Mus musculus GN=Mdh2 PE=1 SV=3 | MDHM_MOUSE | Mdh2 | 36 kDa | 0.035 | KO Synap low, WT Synap high | 4 | 19 | 16 | 24 | 31 | 27 |
| 13 | Dynamin-1 OS=Mus musculus GN=Dnm1 PE=1 SV=1 | A0A0J9YUN4_MOUSE | Dnm1 | 97 kDa | 0.0012 | KO Synap low, WT Synap high | 16 | 16 | 16 | 28 | 24 | 27 |
| 14 | Aconitate hydratase, mitochondrial OS=Mus musculus GN=Aco2 PE=1 SV=1 | ACON_MOUSE | Aco2 | 85 kDa | 0.023 | KO Synap low, WT Synap high | 0 | 17 | 17 | 24 | 28 | 27 |
| 15 | Vesicle-fusing ATPase OS=Mus musculus GN=Nsf PE=1 SV=2 | NSF_MOUSE | Nsf | 83 kDa | 0.04 | KO Synap low, WT Synap high | 12 | 12 | 13 | 16 | 22 | 17 |
| 16 | Spectrin alpha chain, non-erythrocytic 1 OS=Mus musculus GN=Sptan1 PE=1 SV=1 | A3KGU7_MOUSE (+1) | Sptan1 | 285 kDa | 0.047 | KO Synap low, WT Synap high | 1 | 10 | 17 | 17 | 31 | 27 |
| 17 | Calcium/calmodulin-dependent protein kinase type II subunit alpha OS=Mus musculus GN=Camk2a PE=1 SV=2 | KCC2A_MOUSE | Camk2a | 54 kDa | 0.0067 | KO Synap low, WT Synap high | 13 | 11 | 14 | 15 | 17 | 17 |
| 18 | Neural cell adhesion molecule 1 OS=Mus musculus GN=Ncam1 PE=1 SV=3 | NCAM1_MOUSE | Ncam1 | 119 kDa | 0.0063 | KO Synap low, WT Synap high | 9 | 9 | 12 | 14 | 13 | 14 |
| 19 | Spectrin beta chain, non-erythrocytic 1 OS=Mus musculus GN=Sptbn1 PE=1 SV=2 | SPTB2_MOUSE | Sptbn1 | 274 kDa | 0.0046 | KO Synap low, WT Synap high | 11 | 8 | 12 | 17 | 21 | 22 |
| 20 | Hexokinase 1, isoform CRA_f OS=Mus musculus GN=Hk1 PE=1 SV=1 | G3UVV4_MOUSE (+1) | Hk1 | 102 kDa | 0.024 | KO Synap low, WT Synap high | 2 | 11 | 12 | 14 | 15 | 18 |
| 21 | 60 kDa heat shock protein, mitochondrial OS=Mus musculus GN=Hspd1 PE=1 SV=1 | CH60_MOUSE | Hspd1 | 61 kDa | 0.0065 | KO Synap low, WT Synap high | 0 | 10 | 13 | 18 | 17 | 20 |
| 22 | Serine/threonine-protein phosphatase 2A 65 kDa regulatory subunit A alpha isoform OS=Mus musculus GN=Ppp2r1a PE=1 SV=3 | 2AAA_MOUSE | Ppp2r1a | 65 kDa | 0.008 | KO Synap low, WT Synap high | 8 | 3 | 8 | 9 | 9 | 9 |
| 23 | Calcium-dependent secretion activator 1 OS=Mus musculus GN=Cadps PE=1 SV=3 | CAPS1_MOUSE | Cadps | 153 kDa | 0.025 | KO Synap low, WT Synap high | 7 | 6 | 6 | 8 | 9 | 8 |
| 24 | Stress-70 protein, mitochondrial OS=Mus musculus GN=Hspa9 PE=1 SV=3 | GRP75_MOUSE | Hspa9 | 73 kDa | 0.021 | KO Synap low, WT Synap high | 0 | 7 | 7 | 13 | 16 | 17 |
| 25 | Ankyrin-2 OS=Mus musculus GN=Ank2 PE=1 SV=2 | ANK2_MOUSE | Ank2 | 426 kDa | 0.0054 | KO Synap low, WT Synap high | 7 | 8 | 9 | 14 | 20 | 16 |
| 26 | Contactin-1 OS=Mus musculus GN=Cntn1 PE=1 SV=1 | CNTN1_MOUSE | Cntn1 | 113 kDa | 0.01 | KO Synap low, WT Synap high | 4 | 4 | 7 | 9 | 11 | 9 |
| 27 | Synaptosomal-associated protein 25 OS=Mus musculus GN=Snap25 PE=1 SV=1 | SNP25_MOUSE | Snap25 | 23 kDa | 0.023 | KO Synap low, WT Synap high | 3 | 6 | 8 | 8 | 10 | 10 |
| 28 | Cytochrome c, somatic OS=Mus musculus GN=Cycs PE=1 SV=2 | CYC_MOUSE | Cycs | 12 kDa | 0.0096 | KO Synap low, WT Synap high | 0 | 4 | 5 | 7 | 8 | 7 |
| 29 | Calcium-binding mitochondrial carrier protein Aralar1 OS=Mus musculus GN=Slc25a12 PE=1 SV=1 | CMC1_MOUSE | Slc25a12 | 75 kDa | 0.025 | KO Synap low, WT Synap high | 0 | 5 | 6 | 11 | 9 | 12 |
| 30 | Cytochrome b-c1 complex subunit 1, mitochondrial OS=Mus musculus GN=Uqcrc1 PE=1 SV=2 | QCR1_MOUSE | Uqcrc1 | 53 kDa | 0.02 | KO Synap low, WT Synap high | 0 | 5 | 7 | 8 | 7 | 8 |
| 31 | Pyruvate dehydrogenase E1 component subunit beta, mitochondrial OS=Mus musculus GN=Pdhb PE=1 SV=1 | ODPB_MOUSE | Pdhb | 39 kDa | 0.045 | KO Synap low, WT Synap high | 0 | 6 | 9 | 9 | 10 | 7 |
| 32 | AP-2 complex subunit alpha-2 OS=Mus musculus GN=Ap2a2 PE=1 SV=2 | AP2A2_MOUSE | Ap2a2 | 104 kDa | 0.0064 | KO Synap low, WT Synap high | 3 | 2 | 3 | 8 | 5 | 9 |
| 33 | Beta-soluble NSF attachment protein OS=Mus musculus GN=Napb PE=1 SV=2 | SNAB_MOUSE | Napb | 34 kDa | 0.0011 | KO Synap low, WT Synap high | 2 | 3 | 4 | 5 | 6 | 5 |
| 34 | Dynamin-like 120 kDa protein, mitochondrial OS=Mus musculus GN=Opa1 PE=1 SV=1 | OPA1_MOUSE | Opa1 | 111 kDa | 0.02 | KO Synap low, WT Synap high | 0 | 7 | 8 | 11 | 13 | 11 |
| 35 | 2',3'-cyclic-nucleotide 3'-phosphodiesterase OS=Mus musculus GN=Cnp PE=1 SV=3 | CN37_MOUSE | Cnp | 47 kDa | 0.017 | KO Synap low, WT Synap high | 1 | 3 | 4 | 6 | 7 | 10 |
| 36 | Isocitrate dehydrogenase [NAD] subunit gamma 1, mitochondrial OS=Mus musculus GN=Idh3g PE=1 SV=1 | IDHG1_MOUSE | Idh3g | 43 kDa | 0.026 | KO Synap low, WT Synap high | 0 | 2 | 1 | 5 | 6 | 6 |
| 37 | Neurofilament 3, medium OS=Mus musculus GN=Nefm PE=1 SV=1 | A0A0R4J036_MOUSE | Nefm | 96 kDa | 0.027 | KO Synap low, WT Synap high | 4 | 1 | 4 | 4 | 8 | 8 |
| 38 | ATP synthase subunit O, mitochondrial OS=Mus musculus GN=Atp5o PE=1 SV=1 | ATPO_MOUSE | Atp5o | 23 kDa | 0.031 | KO Synap low, WT Synap high | 0 | 6 | 5 | 8 | 8 | 9 |
| 39 | ProSAAS OS=Mus musculus GN=Pcsk1n PE=1 SV=2 | PCSK1_MOUSE | Pcsk1n | 27 kDa | 0.018 | KO Synap low, WT Synap high | 2 | 2 | 3 | 5 | 3 | 5 |
| 40 | Dihydrolipoyllysine-residue acetyltransferase component of pyruvate dehydrogenase complex, mitochondrial OS=Mus musculus GN=Dlat PE=1 SV=2 | ODP2_MOUSE | Dlat | 68 kDa | 0.024 | KO Synap low, WT Synap high | 0 | 4 | 5 | 8 | 7 | 8 |
| 41 | NADH dehydrogenase [ubiquinone] 1 alpha subcomplex subunit 9, mitochondrial OS=Mus musculus GN=Ndufa9 PE=1 SV=1 | A0A0R3P9C8_MOUSE (+1) | Ndufa9 | 42 kDa | 0.0052 | KO Synap low, WT Synap high | 0 | 3 | 4 | 7 | 8 | 9 |
| 42 | AP-2 complex subunit beta OS=Mus musculus GN=Ap2b1 PE=1 SV=1 | AP2B1_MOUSE (+1) | Ap2b1 | 105 kDa | 0.045 | KO Synap low, WT Synap high | 3 | 0 | 6 | 10 | 8 | 7 |
| 43 | Fumarate hydratase, mitochondrial OS=Mus musculus GN=Fh PE=1 SV=3 | FUMH_MOUSE | Fh | 54 kDa | 0.027 | KO Synap low, WT Synap high | 1 | 3 | 5 | 7 | 8 | 9 |
| 44 | Septin-7 OS=Mus musculus GN=Sept7 PE=1 SV=2 | E9Q1G8_MOUSE | 7-Sep | 51 kDa | 0.024 | KO Synap low, WT Synap high | 4 | 4 | 6 | 8 | 8 | 7 |
| 45 | Cytochrome b-c1 complex subunit 2, mitochondrial OS=Mus musculus GN=Uqcrc2 PE=1 SV=1 | QCR2_MOUSE | Uqcrc2 | 48 kDa | 0.0052 | KO Synap low, WT Synap high | 0 | 3 | 4 | 7 | 8 | 6 |
| 46 | N(G),N(G)-dimethylarginine dimethylaminohydrolase 1 OS=Mus musculus GN=Ddah1 PE=1 SV=3 | DDAH1_MOUSE | Ddah1 | 31 kDa | 0.024 | KO Synap low, WT Synap high | 2 | 2 | 2 | 3 | 3 | 3 |
| 47 | Peroxiredoxin-5, mitochondrial OS=Mus musculus GN=Prdx5 PE=1 SV=2 | PRDX5_MOUSE (+1) | Prdx5 | 22 kDa | 0.029 | KO Synap low, WT Synap high | 0 | 4 | 1 | 4 | 5 | 6 |
| 48 | Succinate--CoA ligase [ADP-forming] subunit beta, mitochondrial OS=Mus musculus GN=Sucla2 PE=1 SV=2 | SUCB1_MOUSE | Sucla2 | 50 kDa | 0.021 | KO Synap low, WT Synap high | 0 | 2 | 6 | 8 | 9 | 12 |
| 49 | Septin-11 OS=Mus musculus GN=Sept11 PE=1 SV=1 | A0A0J9YTY0_MOUSE (+2) | 11-Sep | 49 kDa | 0.019 | KO Synap low, WT Synap high | 2 | 4 | 4 | 6 | 8 | 7 |
| 50 | Reticulon-4 OS=Mus musculus GN=Rtn4 PE=1 SV=2 | RTN4_MOUSE | Rtn4 | 127 kDa | 0.008 | KO Synap low, WT Synap high | 2 | 2 | 2 | 5 | 5 | 5 |
| 51 | AP-2 complex subunit mu OS=Mus musculus GN=Ap2m1 PE=1 SV=1 | AP2M1_MOUSE (+1) | Ap2m1 | 50 kDa | 0.0062 | KO Synap low, WT Synap high | 3 | 0 | 2 | 4 | 4 | 4 |
| 52 | Inositol 1,4,5-trisphosphate receptor type 1 OS=Mus musculus GN=Itpr1 PE=1 SV=2 | ITPR1_MOUSE | Itpr1 | 313 kDa | 0.019 | KO Synap low, WT Synap high | 2 | 3 | 4 | 7 | 11 | 10 |
| 53 | Isocitrate dehydrogenase [NAD] subunit, mitochondrial OS=Mus musculus GN=Idh3b PE=1 SV=1 | Q91VA7_MOUSE | Idh3b | 42 kDa | 0.0094 | KO Synap low, WT Synap high | 0 | 4 | 3 | 9 | 8 | 11 |
| 54 | Prohibitin OS=Mus musculus GN=Phb PE=1 SV=1 | PHB_MOUSE | Phb | 30 kDa | 0.041 | KO Synap low, WT Synap high | 0 | 2 | 7 | 8 | 9 | 8 |
| 55 | NADH dehydrogenase [ubiquinone] flavoprotein 1, mitochondrial OS=Mus musculus GN=Ndufv1 PE=1 SV=1 | D3YUM1_MOUSE (+1) | Ndufv1 | 50 kDa | 0.024 | KO Synap low, WT Synap high | 0 | 3 | 4 | 6 | 10 | 9 |
| 56 | Glutamate decarboxylase 2 OS=Mus musculus GN=Gad2 PE=1 SV=1 | DCE2_MOUSE | Gad2 | 65 kDa | 0.00039 | KO Synap low, WT Synap high | 3 | 1 | 2 | 5 | 6 | 4 |
| 57 | AP-2 complex subunit alpha-1 OS=Mus musculus GN=Ap2a1 PE=1 SV=1 | AP2A1_MOUSE | Ap2a1 | 108 kDa | 0.039 | KO Synap low, WT Synap high | 5 | 2 | 3 | 2 | 4 | 1 |
| 58 | NADH dehydrogenase [ubiquinone] 1 alpha subcomplex subunit 10, mitochondrial OS=Mus musculus GN=Ndufa10 PE=1 SV=1 | NDUAA_MOUSE | Ndufa10 | 41 kDa | 0.02 | KO Synap low, WT Synap high | 0 | 3 | 5 | 6 | 7 | 9 |
| 59 | Acyl-CoA-binding protein OS=Mus musculus GN=Dbi PE=1 SV=2 | ACBP_MOUSE | Dbi | 10 kDa | 0.011 | KO Synap low, WT Synap high | 0 | 0 | 2 | 2 | 2 | 2 |
| 60 | V-type proton ATPase subunit E 1 OS=Mus musculus GN=Atp6v1e1 PE=1 SV=2 | VATE1_MOUSE | Atp6v1e1 | 26 kDa | 0.041 | KO Synap low, WT Synap high | 3 | 0 | 2 | 2 | 4 | 5 |
| 61 | Tropomyosin 1, alpha, isoform CRA_i OS=Mus musculus GN=Tpm1 PE=1 SV=1 | G5E8R0_MOUSE | Tpm1 | 28 kDa | 0.039 | KO Synap low, WT Synap high | 0 | 1 | 1 | 4 | 3 | 5 |
| 62 | Plasma membrane calcium-transporting ATPase 1 OS=Mus musculus GN=Atp2b1 PE=1 SV=1 | AT2B1_MOUSE | Atp2b1 | 135 kDa | 0.013 | KO Synap low, WT Synap high | 2 | 2 | 2 | 6 | 7 | 4 |
| 63 | Alpha actinin 1a OS=Mus musculus GN=Actn1 PE=1 SV=1 | A1BN54_MOUSE (+1) | Actn1 | 103 kDa | 0.0079 | KO Synap low, WT Synap high | 2 | 1 | 2 | 4 | 5 | 5 |
| 64 | Cytochrome c oxidase subunit 5A, mitochondrial OS=Mus musculus GN=Cox5a PE=1 SV=2 | COX5A_MOUSE | Cox5a | 16 kDa | 0.023 | KO Synap low, WT Synap high | 1 | 2 | 1 | 3 | 4 | 5 |
| 65 | Mitochondrial import receptor subunit TOM70 OS=Mus musculus GN=Tomm70 PE=1 SV=2 | TOM70_MOUSE | Tomm70 | 68 kDa | 0.028 | KO Synap low, WT Synap high | 0 | 2 | 3 | 7 | 7 | 2 |
| 66 | Dihydrolipoyllysine-residue succinyltransferase component of 2-oxoglutarate dehydrogenase complex, mitochondrial OS=Mus musculus GN=Dlst PE=1 SV=1 | ODO2_MOUSE | Dlst | 49 kDa | 0.01 | KO Synap low, WT Synap high | 0 | 2 | 2 | 3 | 5 | 5 |
| 67 | Calretinin OS=Mus musculus GN=Calb2 PE=1 SV=3 | CALB2_MOUSE | Calb2 | 31 kDa | 0.019 | KO Synap low, WT Synap high | 0 | 0 | 0 | 3 | 3 | 2 |
| 68 | Aldehyde dehydrogenase, mitochondrial OS=Mus musculus GN=Aldh2 PE=1 SV=1 | A0A0G2JEU1_MOUSE (+1) | Aldh2 | 51 kDa | 0.028 | KO Synap low, WT Synap high | 0 | 3 | 4 | 7 | 7 | 6 |
| 69 | Catenin alpha-2 OS=Mus musculus GN=Ctnna2 PE=1 SV=3 | CTNA2_MOUSE (+1) | Ctnna2 | 105 kDa | 0.047 | KO Synap low, WT Synap high | 0 | 3 | 4 | 6 | 9 | 4 |
| 70 | Synaptophysin OS=Mus musculus GN=Syp PE=1 SV=2 | SYPH_MOUSE | Syp | 34 kDa | 0.022 | KO Synap low, WT Synap high | 0 | 1 | 1 | 2 | 3 | 3 |
| 71 | LETM1 and EF-hand domain-containing protein 1, mitochondrial OS=Mus musculus GN=Letm1 PE=1 SV=1 | LETM1_MOUSE | Letm1 | 83 kDa | 0.031 | KO Synap low, WT Synap high | 0 | 3 | 1 | 4 | 7 | 7 |
| 72 | Anion exchange protein OS=Mus musculus GN=Slc4a4 PE=1 SV=1 | E9Q8N8_MOUSE | Slc4a4 | 123 kDa | 0.0065 | KO Synap low, WT Synap high | 0 | 0 | 0 | 4 | 6 | 3 |
| 73 | Ras-related protein Rab-7a OS=Mus musculus GN=Rab7a PE=1 SV=2 | RAB7A_MOUSE | Rab7a | 23 kDa | 0.018 | KO Synap low, WT Synap high | 0 | 2 | 1 | 2 | 3 | 3 |
| 74 | Alpha-internexin OS=Mus musculus GN=Ina PE=1 SV=3 | AINX_MOUSE | Ina | 55 kDa | 0.0092 | KO Synap low, WT Synap high | 0 | 0 | 1 | 4 | 6 | 6 |
| 75 | Septin-5 OS=Mus musculus GN=Sept5 PE=1 SV=2 | SEPT5_MOUSE | 5-Sep | 43 kDa | 0.044 | KO Synap low, WT Synap high | 0 | 0 | 3 | 3 | 2 | 3 |
| 76 | AP-1 complex subunit beta-1 OS=Mus musculus GN=Ap1b1 PE=1 SV=2 | AP1B1_MOUSE (+2) | Ap1b1 | 104 kDa | 0.044 | KO Synap low, WT Synap high | 2 | 2 | 2 | 2 | 3 | 2 |
| 77 | Disks large homolog 4 OS=Mus musculus GN=Dlg4 PE=1 SV=1 | DLG4_MOUSE | Dlg4 | 80 kDa | 0.021 | KO Synap low, WT Synap high | 0 | 1 | 2 | 4 | 5 | 7 |
| 78 | Elongation factor 1-beta OS=Mus musculus GN=Eef1b PE=1 SV=5 | EF1B_MOUSE | Eef1b | 25 kDa | 0.016 | KO Synap low, WT Synap high | 1 | 0 | 1 | 1 | 1 | 1 |
| 79 | NADH dehydrogenase [ubiquinone] iron-sulfur protein 7, mitochondrial OS=Mus musculus GN=Ndufs7 PE=1 SV=1 | NDUS7_MOUSE | Ndufs7 | 25 kDa | 0.024 | KO Synap low, WT Synap high | 0 | 2 | 1 | 3 | 2 | 3 |
| 80 | Cytochrome b-c1 complex subunit 6, mitochondrial OS=Mus musculus GN=Uqcrh PE=1 SV=2 | QCR6_MOUSE | Uqcrh | 10 kDa | 0.018 | KO Synap low, WT Synap high | 0 | 1 | 1 | 2 | 2 | 2 |
| 81 | Serine/threonine-protein phosphatase 2A catalytic subunit alpha isoform OS=Mus musculus GN=Ppp2ca PE=1 SV=1 | PP2AA_MOUSE | Ppp2ca | 36 kDa | 0.016 | KO Synap low, WT Synap high | 1 | 0 | 1 | 2 | 2 | 2 |
| 82 | NADH dehydrogenase [ubiquinone] 1 alpha subcomplex subunit 8 OS=Mus musculus GN=Ndufa8 PE=1 SV=3 | NDUA8_MOUSE | Ndufa8 | 20 kDa | 0.016 | KO Synap low, WT Synap high | 0 | 0 | 1 | 3 | 4 | 2 |
| 83 | Protein disulfide-isomerase A6 OS=Mus musculus GN=Pdia6 PE=1 SV=1 | Q3TML0_MOUSE | Pdia6 | 49 kDa | 0.0075 | KO Synap low, WT Synap high | 1 | 1 | 1 | 1 | 1 | 1 |
| 84 | Actin-related protein 3 OS=Mus musculus GN=Actr3 PE=1 SV=3 | ARP3_MOUSE | Actr3 | 47 kDa | 0.0021 | KO Synap low, WT Synap high | 0 | 1 | 0 | 4 | 3 | 2 |
| 85 | Neuroligin-2 OS=Mus musculus GN=Nlgn2 PE=1 SV=2 | NLGN2_MOUSE | Nlgn2 | 91 kDa | 0.018 | KO Synap low, WT Synap high | 0 | 1 | 1 | 3 | 4 | 5 |
| 86 | Dynamin-3 OS=Mus musculus GN=Dnm3 PE=1 SV=1 | DYN3_MOUSE (+1) | Dnm3 | 97 kDa | 0.018 | KO Synap low, WT Synap high | 0 | 1 | 0 | 4 | 6 | 4 |
| 87 | Rho-related GTP-binding protein RhoB OS=Mus musculus GN=Rhob PE=1 SV=1 | RHOB_MOUSE | Rhob | 22 kDa | 0.035 | KO Synap low, WT Synap high | 0 | 1 | 2 | 3 | 3 | 2 |
| 88 | Importin subunit beta-1 OS=Mus musculus GN=Kpnb1 PE=1 SV=2 | IMB1_MOUSE | Kpnb1 | 97 kDa | 0.00056 | KO Synap low, WT Synap high | 0 | 0 | 0 | 2 | 4 | 3 |

| | | | | | | | | | | | | |
|---|---|---|---|---|---|---|---|---|---|---|---|---|
| 89 | Methylmalonate-semialdehyde dehydrogenase [acylating], mitochondrial OS=Mus musculus GN=Aldh6a1 PE=1 SV=1 | MMSA_MOUSE | Aldh6a1 | 58 kDa | 0.023 | KO Synap low, WT Synap high | 0 | 1 | 1 | 5 | 2 | 5 |
| 90 | Voltage-dependent anion-selective channel protein 1 OS=Mus musculus GN=Vdac1 PE=1 SV=3 | VDAC1_MOUSE | Vdac1 | 32 kDa | 0.04 | KO Synap low, WT Synap high | 0 | 0 | 0 | 1 | 3 | 4 |
| 91 | NADH dehydrogenase [ubiquinone] iron-sulfur protein 5 OS=Mus musculus GN=Ndufs5 PE=1 SV=3 | NDUS5_MOUSE | Ndufs5 | 13 kDa | 0.0048 | KO Synap low, WT Synap high | 0 | 0 | 1 | 2 | 3 | 3 |
| 92 | Serine/threonine-protein phosphatase PP1-gamma catalytic subunit OS=Mus musculus GN=Ppp1cc PE=1 SV=1 | PP1G_MOUSE | Ppp1cc | 37 kDa | 0.047 | KO Synap low, WT Synap high | 1 | 1 | 0 | 2 | 3 | 4 |
| 93 | Anion exchange protein OS=Mus musculus GN=Slc4a10 PE=1 SV=1 | B1AWV9_MOUSE (+1) | Slc4a10 | 125 kDa | 0.018 | KO Synap low, WT Synap high | 0 | 1 | 0 | 6 | 3 | 4 |
| 94 | Cytochrome b-c1 complex subunit 10 OS=Mus musculus GN=Uqcr11 PE=3 SV=1 | QCR10_MOUSE | Uqcr11 | 7 kDa | 0.0061 | KO Synap low, WT Synap high | 0 | 2 | 0 | 2 | 2 | 2 |
| 95 | Cytosolic non-specific dipeptidase OS=Mus musculus GN=Cndp2 PE=1 SV=1 | CNDP2_MOUSE | Cndp2 | 53 kDa | 0.013 | KO Synap low, WT Synap high | 1 | 1 | 0 | 1 | 2 | 3 |
| 96 | V-type proton ATPase subunit G 2 OS=Mus musculus GN=Atp6v1g2 PE=1 SV=1 | VATG2_MOUSE | Atp6v1g2 | 14 kDa | 0.013 | KO Synap low, WT Synap high | 0 | 1 | 1 | 2 | 1 | 2 |
| 97 | F-actin-capping protein subunit alpha-2 OS=Mus musculus GN=Capza2 PE=1 SV=3 | CAZA2 MOUSE | Capza2 | 33 kDa | 0.0078 | KO Synap low, WT Synap high | 0 | 0 | 1 | 1 | 2 | 1 |
| 98 | Superoxide dismutase [Mn], mitochondrial OS=Mus musculus GN=Sod2 PE=1 SV=3 | SODM MOUSE | Sod2 | 25 kDa | 0.029 | KO Synap low, WT Synap high | 0 | 0 | 2 | 3 | 2 | 4 |
| 99 | Electron transfer flavoprotein subunit beta OS=Mus musculus GN=Etfb PE=1 SV=3 | ETFB MOUSE | Etfb | 28 kDa | 0.016 | KO Synap low, WT Synap high | 0 | 1 | 2 | 3 | 2 | 3 |
| 100 | Casein kinase II subunit alpha OS=Mus musculus GN=Csnk2a1 PE=1 SV=2 | CSK21 MOUSE | Csnk2a1 | 45 kDa | 0.016 | KO Synap low, WT Synap high | 0 | 0 | 0 | 2 | 3 | 1 |
| 101 | MAGUK p55 subfamily member 2 OS=Mus musculus GN=Mpp2 PE=1 SV=1 | MPP2_MOUSE | Mpp2 | 62 kDa | 0.016 | KO Synap low, WT Synap high | 0 | 0 | 2 | 3 | 2 | 3 |
| 102 | Glycerol-3-phosphate dehydrogenase, mitochondrial OS=Mus musculus GN=Gpd2 PE=1 SV=2 | GPDM_MOUSE | Gpd2 | 81 kDa | 0.033 | KO Synap low, WT Synap high | 0 | 0 | 1 | 3 | 4 | 2 |
| 103 | Type I inositol 3,4-bisphosphate 4-phosphatase OS=Mus musculus GN=Inpp4a PE=1 SV=2 | E9Q9A0_MOUSE (+5) | Inpp4a | 105 kDa | 0.0075 | KO Synap low, WT Synap high | 1 | 1 | 1 | 3 | 2 | 2 |
| 104 | Glia maturation factor beta OS=Mus musculus GN=Gmfb PE=1 SV=3 | GMFB_MOUSE | Gmfb | 17 kDa | 0.025 | KO Synap low, WT Synap high | 0 | 0 | 0 | 2 | 1 | 2 |
| 105 | Homer protein homolog 1 OS=Mus musculus GN=Homer1 PE=1 SV=1 | E9Q0I7_MOUSE (+1) | Homer1 | 22 kDa | 0.0065 | KO Synap low, WT Synap high | 0 | 0 | 0 | 1 | 2 | 2 |
| 106 | NADH dehydrogenase [ubiquinone] 1 beta subcomplex subunit 10 (Fragment) OS=Mus musculus GN=Ndufb10 PE=1 SV=1 | D3YUK4_MOUSE (+1) | Ndufb10 | 18 kDa | 0.0022 | KO Synap low, WT Synap high | 0 | 1 | 1 | 1 | 2 | 2 |
| 107 | NADH dehydrogenase [ubiquinone] iron-sulfur protein 2, mitochondrial OS=Mus musculus GN=Ndufs2 PE=1 SV=1 | NDUS2_MOUSE | Ndufs2 | 53 kDa | 0.0078 | KO Synap low, WT Synap high | 0 | 0 | 1 | 2 | 2 | 2 |
| 108 | Glutamate receptor 2 OS=Mus musculus GN=Gria2 PE=1 SV=1 | A0A0A6YW90_MOUSE (+2) | Gria2 | 87 kDa | 0.024 | KO Synap low, WT Synap high | 0 | 1 | 1 | 2 | 3 | 2 |
| 109 | Metaxin-2 OS=Mus musculus GN=Mtx2 PE=1 SV=1 | MTX2_MOUSE | Mtx2 | 30 kDa | 0.0031 | KO Synap low, WT Synap high | 0 | 1 | 0 | 3 | 3 | 3 |
| 110 | Complement component 1 Q subcomponent-binding protein, mitochondrial OS=Mus musculus GN=C1qbp PE=1 SV=1 | Q8R5L1 MOUSE | C1qbp | 31 kDa | 0.0065 | KO Synap low, WT Synap high | 0 | 0 | 0 | 2 | 2 | 2 |
| 111 | Mitochondrial import receptor subunit TOM22 homolog OS=Mus musculus GN=Tomm22 PE=1 SV=3 | TOM22 MOUSE | Tomm22 | 16 kDa | 0.0023 | KO Synap low, WT Synap high | 0 | 0 | 0 | 1 | 2 | 1 |
| 112 | Basigin OS=Mus musculus GN=Bsg PE=1 SV=2 | BASI MOUSE (+2) | Bsg | 42 kDa | 0.0075 | KO Synap low, WT Synap high | 0 | 1 | 0 | 2 | 2 | 2 |
| 113 | Arf-GAP with GTPase, ANK repeat and PH domain-containing protein 2 OS=Mus musculus GN=Agap2 PE=1 SV=1 | AGAP2 MOUSE | Agap2 | 125 kDa | 0.026 | KO Synap low, WT Synap high | 0 | 0 | 0 | 1 | 2 | 2 |
| 114 | Dynamin-1 OS=Mus musculus GN=Dnm1 PE=1 SV=2 | DYN1_MOUSE | Dnm1 | 98 kDa | 0.016 | KO Synap low, WT Synap high | 0 | 0 | 1 | 1 | 1 | 1 |
| 115 | MCG10748, isoform CRA_b OS=Mus musculus GN=Rap1a PE=1 SV=1 | A0A0G2JDL9_MOUSE (+2) | Rap1a | 13 kDa | 0.024 | KO Synap low, WT Synap high | 0 | 1 | 1 | 2 | 3 | 2 |
| 116 | Microtubule-associated protein RP/EB family member 3 OS=Mus musculus GN=Mapre3 PE=1 SV=1 | D3Z6G3_MOUSE (+1) | Mapre3 | 30 kDa | 0.0022 | KO Synap low, WT Synap high | 0 | 0 | 0 | 2 | 1 | 2 |
| 117 | Serine/threonine-protein phosphatase 2A 56 kDa regulatory subunit epsilon isoform OS=Mus musculus GN=Ppp2r5e PE=1 SV=3 | 2A5E_MOUSE | Ppp2r5e | 55 kDa | 0.0075 | KO Synap low, WT Synap high | 0 | 0 | 1 | 1 | 1 | 1 |
| 118 | V-type proton ATPase subunit D OS=Mus musculus GN=Atp6v1d PE=1 SV=1 | VATD_MOUSE | Atp6v1d | 28 kDa | 0.0022 | KO Synap low, WT Synap high | 0 | 0 | 0 | 2 | 1 | 1 |
| 119 | Ras-related protein Rab-3C OS=Mus musculus GN=Rab3c PE=1 SV=1 | RAB3C_MOUSE | Rab3c | 26 kDa | 0.034 | KO Synap low, WT Synap high | 0 | 1 | 0 | 1 | 2 | 2 |
| 120 | Protein tweety homolog OS=Mus musculus GN=Ttyh1 PE=1 SV=1 | A0A0U1RPU8_MOUSE (+1) | Ttyh1 | 39 kDa | 0.016 | KO Synap low, WT Synap high | 0 | 1 | 1 | 1 | 1 | 1 |
| 121 | ADP-ribosylation factor-like protein 3 OS=Mus musculus GN=Arl3 PE=1 SV=1 | ARL3_MOUSE | Arl3 | 20 kDa | 0.0075 | KO Synap low, WT Synap high | 0 | 0 | 0 | 1 | 1 | 1 |
| 122 | Disks large homolog 2 OS=Mus musculus GN=Dlg2 PE=1 SV=2 | DLG2 MOUSE | Dlg2 | 95 kDa | 0.0065 | KO Synap low, WT Synap high | 0 | 0 | 0 | 3 | 1 | 2 |
| 123 | Voltage-dependent calcium channel subunit alpha-2/delta-1 OS=Mus musculus GN=Cacna2d1 PE=1 SV=1 | CA2D1 MOUSE (+1) | Cacna2d1 | 125 kDa | 0.013 | KO Synap low, WT Synap high | 0 | 0 | 1 | 2 | 3 | 2 |
| 124 | Histone H1.0 OS=Mus musculus GN=H1f0 PE=2 SV=4 | H10 MOUSE | H1f0 | 21 kDa | 0.0013 | KO Synap low, WT Synap high | 0 | 0 | 0 | 2 | 1 | 2 |
| 125 | ERC protein 2 OS=Mus musculus GN=Erc2 PE=1 SV=2 | ERC2 MOUSE (+1) | Erc2 | 111 kDa | 0.016 | KO Synap low, WT Synap high | 0 | 0 | 0 | 1 | 2 | 1 |
| 126 | CB1 cannabinoid receptor-interacting protein 1 OS=Mus musculus GN=Cnrip1 PE=1 SV=1 | CNRP1_MOUSE | Cnrip1 | 19 kDa | 0.0022 | KO Synap low, WT Synap high | 0 | 0 | 0 | 1 | 2 | 2 |
| 127 | Disks large-associated protein 2 OS=Mus musculus GN=Dlgap2 PE=1 SV=2 | DLGP2_MOUSE (+1) | Dlgap2 | 119 kDa | 0.016 | KO Synap low, WT Synap high | 0 | 0 | 0 | 1 | 2 | 1 |
| 128 | Zinc transporter 9 OS=Mus musculus GN=Slc30a9 PE=1 SV=2 | ZNT9_MOUSE (+1) | Slc30a9 | 63 kDa | 0.016 | KO Synap low, WT Synap high | 0 | 0 | 0 | 1 | 2 | 1 |
| 129 | Neuronal pentraxin-1 OS=Mus musculus GN=Nptx1 PE=1 SV=1 | NPTX1_MOUSE | Nptx1 | 47 kDa | 0.016 | KO Synap low, WT Synap high | 0 | 0 | 0 | 1 | 1 | 1 |
| 130 | Syntaxin-binding protein 5 OS=Mus musculus GN=Stxbp5 PE=1 SV=1 | D3Z079_MOUSE (+2) | Stxbp5 | 124 kDa | 0.0075 | KO Synap low, WT Synap high | 0 | 0 | 0 | 1 | 2 | 2 |
| 131 | 3-ketoacyl-CoA thiolase, mitochondrial OS=Mus musculus GN=Acaa2 PE=1 SV=3 | THIM_MOUSE | Acaa2 | 42 kDa | 0.0075 | KO Synap low, WT Synap high | 0 | 0 | 0 | 2 | 2 | 1 |
| 132 | Atlastin-2 OS=Mus musculus GN=Atl2 PE=1 SV=1 | ATLA2_MOUSE (+1) | Atl2 | 66 kDa | 0.0075 | KO Synap low, WT Synap high | 0 | 0 | 0 | 1 | 1 | 1 |
| 133 | WD repeat-containing protein 7 OS=Mus musculus GN=Wdr7 PE=1 SV=3 | WDR7_MOUSE | Wdr7 | 163 kDa | 0.016 | KO Synap low, WT Synap high | 0 | 0 | 0 | 1 | 2 | 1 |
| 134 | Tetratricopeptide repeat protein 7B OS=Mus musculus GN=Ttc7b PE=1 SV=1 | TTC7B_MOUSE | Ttc7b | 94 kDa | 0.016 | KO Synap low, WT Synap high | 0 | 0 | 0 | 2 | 1 | 1 |
| 135 | Glutamate receptor 4 OS=Mus musculus GN=Gria4 PE=1 SV=1 | G5E863 MOUSE (+2) | Gria4 | 101 kDa | 0.016 | KO Synap low, WT Synap high | 0 | 0 | 0 | 1 | 1 | 1 |
| 136 | NADP-dependent malic enzyme, mitochondrial OS=Mus musculus GN=Me3 PE=1 SV=2 | MAON MOUSE | Me3 | 67 kDa | 0.016 | KO Synap low, WT Synap high | 0 | 0 | 0 | 1 | 1 | 1 |
| 137 | Microsomal glutathione S-transferase 3 OS=Mus musculus GN=Mgst3 PE=1 SV=1 | MGST3 MOUSE | Mgst3 | 17 kDa | 0.016 | KO Synap low, WT Synap high | 0 | 0 | 0 | 1 | 1 | 1 |

**List of reduced proteins in synaptosomes from brains of 6 month-old mice**

| # | Identified Proteins | Accession Number | Alternate ID | Molecular Weight | T-Test (p-value): (p < 0.05) | Quantitative Profile | KO S4 | KO S5 | KO S6 | WT S4 | WT S5 | WT S6 |
|---|---|---|---|---|---|---|---|---|---|---|---|---|
| 435 | Sodium/potassium-transporting ATPase subunit alpha OS=Mus musculus GN=Atp1a3 PE=1 SV=1 | A0A0G2JGX4_MOUSE (+1) | Atp1a3 | 113 kDa | 0.0015 | KO Synap low, WT Synap high | 46 | 52 | 57 | 54 | 59 | 62 |
| 436 | Synapsin-1 OS=Mus musculus GN=Syn1 PE=1 SV=2 | SYN1_MOUSE | Syn1 | 74 kDa | < 0.00010 | KO Synap low, WT Synap high | 24 | 24 | 21 | 39 | 39 | 40 |
| 437 | Aconitate hydratase, mitochondrial OS=Mus musculus GN=Aco2 PE=1 SV=1 | ACON_MOUSE | Aco2 | 85 kDa | < 0.00010 | KO Synap low, WT Synap high | 13 | 9 | 13 | 64 | 59 | 61 |
| 438 | Syntaxin-binding protein 1 OS=Mus musculus GN=Stxbp1 PE=1 SV=2 | STXB1_MOUSE | Stxbp1 | 68 kDa | 0.0001 | KO Synap low, WT Synap high | 23 | 20 | 24 | 41 | 47 | 46 |
| 439 | Dynamin-1 OS=Mus musculus GN=Dnm1 PE=1 SV=1 | A0A0J9YUN4_MOUSE | Dnm1 | 97 kDa | 0.0046 | KO Synap low, WT Synap high | 43 | 41 | 33 | 53 | 53 | 54 |
| 440 | Clathrin heavy chain 1 OS=Mus musculus GN=Cltc PE=1 SV=3 | CLH1_MOUSE (+1) | Cltc | 192 kDa | 0.0013 | KO Synap low, WT Synap high | 27 | 31 | 21 | 41 | 50 | 50 |
| 441 | Fructose-bisphosphate aldolase A OS=Mus musculus GN=Aldoa PE=1 SV=2 | ALDOA_MOUSE (+1) | Aldoa | 39 kDa | 0.00075 | KO Synap low, WT Synap high | 23 | 20 | 24 | 25 | 31 | 31 |
| 442 | Brain acid soluble protein 1 OS=Mus musculus GN=Basp1 PE=1 SV=3 | BASP1_MOUSE | Basp1 | 22 kDa | < 0.00010 | KO Synap low, WT Synap high | 13 | 12 | 14 | 28 | 28 | 35 |
| 443 | ATP synthase subunit beta, mitochondrial OS=Mus musculus GN=Atp5b PE=1 SV=2 | ATPB_MOUSE | Atp5b | 56 kDa | < 0.00010 | KO Synap low, WT Synap high | 3 | 3 | 4 | 20 | 22 | 25 |
| 444 | Microtubule-associated protein 6 OS=Mus musculus GN=Map6 PE=1 SV=2 | MAP6_MOUSE | Map6 | 96 kDa | 0.00026 | KO Synap low, WT Synap high | 24 | 19 | 19 | 27 | 24 | 34 |
| 445 | Vesicle-fusing ATPase OS=Mus musculus GN=Nsf PE=1 SV=2 | NSF_MOUSE | Nsf | 83 kDa | 0.00048 | KO Synap low, WT Synap high | 28 | 26 | 27 | 29 | 30 | 31 |
| 446 | Myelin basic protein (Fragment) OS=Mus musculus GN=Mbp PE=1 SV=1 | F6VME3_MOUSE | Mbp | 21 kDa | 0.05 | KO Synap low, WT Synap high | 20 | 0 | 0 | 18 | 21 | 23 |
| 447 | Glutamate dehydrogenase 1, mitochondrial OS=Mus musculus GN=Glud1 PE=1 SV=1 | DHE3_MOUSE | Glud1 | 61 kDa | < 0.00010 | KO Synap low, WT Synap high | 3 | 3 | 2 | 29 | 33 | 35 |
| 448 | Excitatory amino acid transporter 2 OS=Mus musculus GN=Slc1a2 PE=1 SV=1 | EAA2_MOUSE | Slc1a2 | 62 kDa | 0.0024 | KO Synap low, WT Synap high | 12 | 10 | 14 | 17 | 19 | 20 |
| 449 | Hexokinase 1, isoform CRA_f OS=Mus musculus GN=Hk1 PE=1 SV=1 | G3UVV4_MOUSE (+1) | Hk1 | 102 kDa | < 0.00010 | KO Synap low, WT Synap high | 9 | 9 | 12 | 33 | 33 | 34 |
| 450 | Spectrin alpha chain, non-erythrocytic 1 OS=Mus musculus GN=Sptan1 PE=1 SV=1 | A3KGU7_MOUSE (+1) | Sptan1 | 285 kDa | 0.0059 | KO Synap low, WT Synap high | 1 | 28 | 24 | 45 | 42 | 47 |
| 451 | Guanine nucleotide-binding protein G(o) subunit alpha OS=Mus musculus GN=Gnao1 PE=1 SV=3 | GNAO_MOUSE | Gnao1 | 40 kDa | 0.0005 | KO Synap low, WT Synap high | 16 | 20 | 20 | 27 | 24 | 23 |
| 452 | Sodium/potassium-transporting ATPase subunit alpha-2 OS=Mus musculus GN=Atp1a2 PE=1 SV=1 | AT1A2_MOUSE | Atp1a2 | 112 kDa | 0.00038 | KO Synap low, WT Synap high | 19 | 20 | 23 | 34 | 42 | 35 |
| 453 | ATP synthase subunit alpha, mitochondrial OS=Mus musculus GN=Atp5a1 PE=1 SV=1 | ATPA_MOUSE | Atp5a1 | 60 kDa | < 0.00010 | KO Synap low, WT Synap high | 1 | 1 | 2 | 28 | 31 | 36 |
| 454 | Synapsin-2 OS=Mus musculus GN=Syn2 PE=1 SV=2 | SYN2_MOUSE | Syn2 | 63 kDa | < 0.00010 | KO Synap low, WT Synap high | 11 | 10 | 9 | 20 | 20 | 22 |
| 455 | Neuromodulin OS=Mus musculus GN=Gap43 PE=1 SV=1 | NEUM_MOUSE | Gap43 | 24 kDa | 0.00071 | KO Synap low, WT Synap high | 17 | 12 | 14 | 21 | 17 | 20 |
| 456 | Calcium/calmodulin-dependent protein kinase type II subunit alpha OS=Mus musculus GN=Camk2a PE=1 SV=2 | KCC2A_MOUSE | Camk2a | 54 kDa | 0.018 | KO Synap low, WT Synap high | 20 | 15 | 13 | 19 | 16 | 17 |
| 457 | 14-3-3 protein zeta/delta OS=Mus musculus GN=Ywhaz PE=1 SV=1 | 1433Z_MOUSE | Ywhaz | 28 kDa | 0.00029 | KO Synap low, WT Synap high | 14 | 14 | 13 | 16 | 15 | 16 |
| 458 | Malate dehydrogenase, mitochondrial OS=Mus musculus GN=Mdh2 PE=1 SV=3 | MDHM_MOUSE | Mdh2 | 36 kDa | < 0.00010 | KO Synap low, WT Synap high | 0 | 0 | 0 | 27 | 29 | 31 |
| 459 | 60 kDa heat shock protein, mitochondrial OS=Mus musculus GN=Hspd1 PE=1 SV=1 | CH60_MOUSE | Hspd1 | 61 kDa | < 0.00010 | KO Synap low, WT Synap high | 4 | 3 | 3 | 28 | 23 | 32 |
| 460 | Citrate synthase, mitochondrial OS=Mus musculus GN=Cs PE=1 SV=1 | CISY_MOUSE | Cs | 52 kDa | < 0.00010 | KO Synap low, WT Synap high | 3 | 3 | 3 | 31 | 28 | 30 |
| 461 | Ras-related protein Rab-3A OS=Mus musculus GN=Rab3a PE=1 SV=1 | RAB3A_MOUSE | Rab3a | 25 kDa | 0.0087 | KO Synap low, WT Synap high | 14 | 16 | 17 | 15 | 14 | 15 |
| 462 | Syntaxin-1B OS=Mus musculus GN=Stx1b PE=1 SV=1 | STX1B_MOUSE | Stx1b | 33 kDa | < 0.00010 | KO Synap low, WT Synap high | 10 | 8 | 11 | 20 | 22 | 21 |
| 463 | Microtubule-associated protein OS=Mus musculus GN=Mapt PE=1 SV=1 | A0A0A0MQC7_MOUSE (+2) | Mapt | 76 kDa | 0.034 | KO Synap low, WT Synap high | 20 | 20 | 15 | 19 | 17 | 20 |
| 464 | Synaptosomal-associated protein 25 OS=Mus musculus GN=Snap25 PE=1 SV=1 | SNP25_MOUSE | Snap25 | 23 kDa | < 0.00010 | KO Synap low, WT Synap high | 14 | 14 | 12 | 17 | 17 | 16 |
| 465 | Protein kinase C and casein kinase substrate in neurons protein 1 OS=Mus musculus GN=Pacsin1 PE=1 SV=1 | PACN1_MOUSE | Pacsin1 | 51 kDa | 0.023 | KO Synap low, WT Synap high | 25 | 25 | 21 | 23 | 18 | 23 |
| 466 | Cell cycle exit and neuronal differentiation protein 1 OS=Mus musculus GN=Cend1 PE=1 SV=1 | CEND_MOUSE | Cend1 | 15 kDa | 0.00022 | KO Synap low, WT Synap high | 10 | 8 | 7 | 19 | 20 | 21 |
| 467 | V-type proton ATPase catalytic subunit A OS=Mus musculus GN=Atp6v1a PE=1 SV=2 | VATA_MOUSE | Atp6v1a | 68 kDa | 0.039 | KO Synap low, WT Synap high | 13 | 14 | 13 | 15 | 14 | 18 |
| 468 | Creatine kinase U-type, mitochondrial OS=Mus musculus GN=Ckmt1 PE=1 SV=1 | KCRU_MOUSE | Ckmt1 | 47 kDa | 0.0014 | KO Synap low, WT Synap high | 3 | 3 | 3 | 27 | 19 | 23 |
| 469 | NADH-ubiquinone oxidoreductase 75 kDa subunit, mitochondrial OS=Mus musculus GN=Ndufs1 PE=1 SV=2 | NDUS1_MOUSE | Ndufs1 | 80 kDa | < 0.00010 | KO Synap low, WT Synap high | 0 | 0 | 1 | 28 | 28 | 31 |
| 470 | Protein bassoon OS=Mus musculus GN=Bsn PE=1 SV=4 | BSN_MOUSE | Bsn | 419 kDa | < 0.00010 | KO Synap low, WT Synap high | 1 | 0 | 1 | 46 | 45 | 48 |
| 471 | Pyruvate dehydrogenase E1 component subunit beta, mitochondrial OS=Mus musculus GN=Pdhb PE=1 SV=1 | ODPB_MOUSE | Pdhb | 39 kDa | < 0.00010 | KO Synap low, WT Synap high | 1 | 0 | 0 | 17 | 18 | 17 |
| 472 | Endophilin-A1 OS=Mus musculus GN=Sh3gl2 PE=1 SV=1 | A2ALV3_MOUSE (+1) | Sh3gl2 | 48 kDa | 0.007 | KO Synap low, WT Synap high | 11 | 16 | 11 | 12 | 12 | 14 |
| 473 | Acetyl-CoA acetyltransferase, mitochondrial OS=Mus musculus GN=Acat1 PE=1 SV=1 | THIL_MOUSE | Acat1 | 45 kDa | < 0.00010 | KO Synap low, WT Synap high | 0 | 0 | 0 | 19 | 19 | 22 |
| 474 | Vesicle-associated membrane protein 2 OS=Mus musculus GN=Vamp2 PE=1 SV=1 | B0QZN5_MOUSE (+1) | Vamp2 | 18 kDa | 0.0026 | KO Synap low, WT Synap high | 11 | 9 | 11 | 10 | 11 | 11 |
| 475 | Calcium-binding mitochondrial carrier protein Aralar1 OS=Mus musculus GN=Slc25a12 PE=1 SV=1 | CMC1_MOUSE | Slc25a12 | 75 kDa | 0.00045 | KO Synap low, WT Synap high | 0 | 0 | 0 | 23 | 22 | 27 |
| 476 | 2-oxoglutarate dehydrogenase, mitochondrial OS=Mus musculus GN=Ogdh PE=1 SV=3 | ODO1_MOUSE (+1) | Ogdh | 116 kDa | 0.00015 | KO Synap low, WT Synap high | 1 | 1 | 1 | 23 | 26 | 30 |
| 477 | Sodium/potassium-transporting ATPase subunit alpha-1 OS=Mus musculus GN=Atp1a1 PE=1 SV=1 | AT1A1_MOUSE | Atp1a1 | 113 kDa | 0.0005 | KO Synap low, WT Synap high | 12 | 12 | 12 | 14 | 18 | 16 |
| 478 | MICOS complex subunit Mic60 OS=Mus musculus GN=Immt PE=1 SV=1 | MIC60_MOUSE | Immt | 84 kDa | < 0.00010 | KO Synap low, WT Synap high | 0 | 0 | 0 | 28 | 31 | 34 |
| 479 | Septin-7 OS=Mus musculus GN=Sept7 PE=1 SV=2 | E9Q1G8_MOUSE | 7-Sep | 51 kDa | < 0.00010 | KO Synap low, WT Synap high | 9 | 8 | 7 | 20 | 23 | 18 |
| 480 | Calcium/calmodulin-dependent protein kinase type II subunit beta OS=Mus musculus GN=Camk2b PE=1 SV=2 | KCC2B_MOUSE | Camk2b | 60 kDa | 0.029 | KO Synap low, WT Synap high | 13 | 14 | 9 | 16 | 14 | 16 |
| 481 | Stress-70 protein, mitochondrial OS=Mus musculus GN=Hspa9 PE=1 SV=3 | GRP75_MOUSE | Hspa9 | 73 kDa | < 0.00010 | KO Synap low, WT Synap high | 0 | 0 | 0 | 24 | 24 | 27 |
| 482 | 4-aminobutyrate aminotransferase, mitochondrial OS=Mus musculus GN=Abat PE=1 SV=1 | GABT_MOUSE | Abat | 56 kDa | < 0.00010 | KO Synap low, WT Synap high | 0 | 0 | 0 | 17 | 22 | 21 |
| 483 | Pyruvate dehydrogenase E1 component subunit alpha, somatic form, mitochondrial OS=Mus musculus GN=Pdha1 PE=1 SV=1 | ODPA_MOUSE | Pdha1 | 43 kDa | < 0.00010 | KO Synap low, WT Synap high | 0 | 0 | 0 | 20 | 15 | 19 |
| 484 | Neural cell adhesion molecule 1 OS=Mus musculus GN=Ncam1 PE=1 SV=3 | NCAM1_MOUSE | Ncam1 | 119 kDa | < 0.00010 | KO Synap low, WT Synap high | 0 | 0 | 0 | 20 | 22 | 21 |
| 485 | ADP/ATP translocase 1 OS=Mus musculus GN=Slc25a4 PE=1 SV=4 | ADT1_MOUSE | Slc25a4 | 33 kDa | < 0.00010 | KO Synap low, WT Synap high | 0 | 0 | 0 | 22 | 20 | 22 |
| 486 | Peroxiredoxin-5, mitochondrial OS=Mus musculus GN=Prdx5 PE=1 SV=2 | PRDX5_MOUSE | Prdx5 | 22 kDa | 0.00032 | KO Synap low, WT Synap high | 8 | 5 | 5 | 12 | 13 | 15 |
| 487 | Beta-soluble NSF attachment protein OS=Mus musculus GN=Napb PE=1 SV=2 | SNAB_MOUSE | Napb | 34 kDa | 0.00049 | KO Synap low, WT Synap high | 11 | 7 | 9 | 14 | 15 | 15 |
| 488 | Ankyrin-2 OS=Mus musculus GN=Ank2 PE=1 SV=2 | ANK2_MOUSE | Ank2 | 426 kDa | 0.00016 | KO Synap low, WT Synap high | 6 | 3 | 1 | 31 | 37 | 31 |
| 489 | Isocitrate dehydrogenase [NAD] subunit gamma 1, mitochondrial OS=Mus musculus GN=Idh3g PE=1 SV=1 | IDHG1_MOUSE | Idh3g | 43 kDa | 0.00097 | KO Synap low, WT Synap high | 0 | 0 | 0 | 10 | 14 | 13 |
| 490 | Isocitrate dehydrogenase [NAD] subunit, mitochondrial OS=Mus musculus GN=Idh3b PE=1 SV=1 | Q91VA7_MOUSE | Idh3b | 42 kDa | 0.00012 | KO Synap low, WT Synap high | 0 | 0 | 0 | 13 | 17 | 22 |
| 491 | Myc box-dependent-interacting protein 1 OS=Mus musculus GN=Bin1 PE=1 SV=1 | BIN1_MOUSE | Bin1 | 64 kDa | 0.0013 | KO Synap low, WT Synap high | 9 | 8 | 5 | 12 | 13 | 15 |
| 492 | ATP-dependent 6-phosphofructokinase, muscle type OS=Mus musculus GN=Pfkm PE=1 SV=3 | PFKAM_MOUSE | Pfkm | 85 kDa | 0.014 | KO Synap low, WT Synap high | 9 | 5 | 9 | 15 | 14 | 21 |
| 493 | Glutaminase kidney isoform, mitochondrial OS=Mus musculus GN=Gls PE=1 SV=1 | GLSK_MOUSE | Gls | 74 kDa | < 0.00010 | KO Synap low, WT Synap high | 0 | 0 | 0 | 13 | 14 | 17 |
| 494 | Isocitrate dehydrogenase [NAD] subunit alpha, mitochondrial OS=Mus musculus GN=Idh3a PE=1 SV=1 | IDH3A_MOUSE | Idh3a | 40 kDa | < 0.00010 | KO Synap low, WT Synap high | 0 | 0 | 0 | 17 | 15 | 20 |
| 495 | Pyruvate carboxylase OS=Mus musculus GN=Pcx PE=1 SV=1 | E9QPD7_MOUSE (+1) | Pcx | 130 kDa | < 0.00010 | KO Synap low, WT Synap high | 0 | 0 | 0 | 18 | 19 | 20 |
| 496 | Excitatory amino acid transporter 1 OS=Mus musculus GN=Slc1a3 PE=1 SV=2 | EAA1_MOUSE | Slc1a3 | 60 kDa | 0.0019 | KO Synap low, WT Synap high | 3 | 0 | 5 | 9 | 9 | 8 |
| 497 | Elongation factor Tu, mitochondrial OS=Mus musculus GN=Tufm PE=1 SV=1 | EFTU_MOUSE | Tufm | 50 kDa | < 0.00010 | KO Synap low, WT Synap high | 0 | 0 | 0 | 16 | 19 | 19 |
| 498 | ATPase inhibitor, mitochondrial OS=Mus musculus GN=Atpif1 PE=1 SV=2 | ATIF1_MOUSE | Atpif1 | 12 kDa | 0.0019 | KO Synap low, WT Synap high | 2 | 2 | 0 | 8 | 10 | 10 |
| 499 | Succinate--CoA ligase [ADP-forming] subunit beta, mitochondrial OS=Mus musculus GN=Sucla2 PE=1 SV=2 | SUCB1_MOUSE | Sucla2 | 50 kDa | < 0.00010 | KO Synap low, WT Synap high | 0 | 0 | 0 | 14 | 13 | 14 |
| 500 | Succinyl-CoA:3-ketoacid coenzyme A transferase 1, mitochondrial OS=Mus musculus GN=Oxct1 PE=1 SV=1 | SCOT1_MOUSE | Oxct1 | 56 kDa | 0.00099 | KO Synap low, WT Synap high | 1 | 0 | 0 | 17 | 18 | 18 |
| 501 | V-type proton ATPase subunit H OS=Mus musculus GN=Atp6v1h PE=1 SV=1 | A0A0A6YX18_MOUSE (+1) | Atp6v1h | 54 kDa | 0.042 | KO Synap low, WT Synap high | 10 | 9 | 9 | 9 | 8 | 9 |
| 502 | Glutathione S-transferase Mu 1 OS=Mus musculus GN=Gstm1 PE=1 SV=2 | A2AE89_MOUSE (+1) | Gstm1 | 29 kDa | 0.03 | KO Synap low, WT Synap high | 6 | 5 | 9 | 11 | 8 | 11 |
| 503 | Sarcoplasmic/endoplasmic reticulum calcium ATPase 2 OS=Mus musculus GN=Atp2a2 PE=1 SV=2 | AT2A2_MOUSE | Atp2a2 | 115 kDa | 0.0016 | KO Synap low, WT Synap high | 11 | 11 | 12 | 16 | 16 | 16 |
| 504 | NADH dehydrogenase [ubiquinone] flavoprotein 1, mitochondrial OS=Mus musculus GN=Ndufv1 PE=1 SV=1 | D3YUM1_MOUSE (+1) | Ndufv1 | 50 kDa | < 0.00010 | KO Synap low, WT Synap high | 0 | 0 | 0 | 12 | 13 | 14 |
| 505 | Alpha-synuclein OS=Mus musculus GN=Snca PE=1 SV=2 | SYUA_MOUSE | Snca | 14 kDa | 0.0064 | KO Synap low, WT Synap high | 4 | 6 | 9 | 7 | 6 | 5 |
| 506 | Mitochondrial import receptor subunit TOM70 OS=Mus musculus GN=Tomm70 PE=1 SV=2 | TOM70_MOUSE | Tomm70 | 68 kDa | < 0.00010 | KO Synap low, WT Synap high | 0 | 0 | 0 | 15 | 14 | 16 |
| 507 | Protein piccolo OS=Mus musculus GN=Pclo PE=1 SV=4 | PCLO_MOUSE | Pclo | 551 kDa | 0.0014 | KO Synap low, WT Synap high | 1 | 1 | 0 | 23 | 30 | 22 |
| 508 | 14-3-3 protein eta OS=Mus musculus GN=Ywhah PE=1 SV=2 | 1433F_MOUSE | Ywhah | 28 kDa | 0.0065 | KO Synap low, WT Synap high | 9 | 8 | 10 | 9 | 9 | 11 |
| 509 | Rabphilin-3A OS=Mus musculus GN=Rph3a PE=1 SV=2 | RP3A_MOUSE | Rph3a | 75 kDa | < 0.00010 | KO Synap low, WT Synap high | 6 | 6 | 6 | 14 | 10 | 12 |
| 510 | Dynamin-like 120 kDa protein, mitochondrial OS=Mus musculus GN=Opa1 PE=1 SV=1 | OPA1_MOUSE | Opa1 | 111 kDa | 0.00014 | KO Synap low, WT Synap high | 0 | 0 | 0 | 16 | 15 | 19 |
| 511 | Cytochrome c oxidase subunit 5B, mitochondrial OS=Mus musculus GN=Cox5b PE=1 SV=1 | COX5B_MOUSE (+1) | Cox5b | 14 kDa | 0.00015 | KO Synap low, WT Synap high | 2 | 0 | 2 | 8 | 9 | 9 |
| 512 | DmX-like protein 2 OS=Mus musculus GN=Dmxl2 PE=1 SV=3 | DMXL2_MOUSE | Dmxl2 | 338 kDa | 0.0015 | KO Synap low, WT Synap high | 8 | 8 | 7 | 15 | 18 | 16 |
| 513 | AP-2 complex subunit mu OS=Mus musculus GN=Ap2m1 PE=1 SV=1 | AP2M1_MOUSE (+1) | Ap2m1 | 50 kDa | 0.0004 | KO Synap low, WT Synap high | 5 | 5 | 5 | 8 | 7 | 9 |
| 514 | Amphiphysin OS=Mus musculus GN=Amph PE=1 SV=1 | A0A0G2JEG8_MOUSE (+1) | Amph | 75 kDa | 0.0051 | KO Synap low, WT Synap high | 7 | 8 | 3 | 12 | 11 | 13 |
| 515 | Guanine nucleotide-binding protein G(I)/G(S)/G(T) subunit beta-1 OS=Mus musculus GN=Gnb1 PE=1 SV=3 | GBB1_MOUSE | Gnb1 | 37 kDa | < 0.00010 | KO Synap low, WT Synap high | 8 | 6 | 8 | 11 | 10 | 11 |
| 516 | Succinate dehydrogenase [ubiquinone] flavoprotein subunit, mitochondrial OS=Mus musculus GN=Sdha PE=1 SV=1 | SDHA_MOUSE | Sdha | 73 kDa | 0.00011 | KO Synap low, WT Synap high | 0 | 0 | 1 | 17 | 16 | 17 |
| 517 | ProSAAS OS=Mus musculus GN=Pcsk1n PE=1 SV=2 | PCSK1_MOUSE | Pcsk1n | 27 kDa | 0.0057 | KO Synap low, WT Synap high | 5 | 5 | 5 | 8 | 6 | 10 |
| 518 | Cathepsin D OS=Mus musculus GN=Ctsd PE=1 SV=1 | CATD_MOUSE (+1) | Ctsd | 45 kDa | 0.0008 | KO Synap low, WT Synap high | 1 | 2 | 2 | 9 | 8 | 7 |
| 519 | Plasma membrane calcium-transporting ATPase 1 OS=Mus musculus GN=Atp2b1 PE=1 SV=1 | AT2B1_MOUSE | Atp2b1 | 135 kDa | 0.00013 | KO Synap low, WT Synap high | 5 | 5 | 6 | 14 | 13 | 10 |
| 520 | ATP synthase subunit O, mitochondrial OS=Mus musculus GN=Atp5o PE=1 SV=1 | ATPO_MOUSE | Atp5o | 23 kDa | 0.00013 | KO Synap low, WT Synap high | 0 | 0 | 0 | 10 | 11 | 12 |
| 521 | Protein Ogdhl OS=Mus musculus GN=Ogdhl PE=1 SV=1 | E9Q7L0_MOUSE | Ogdhl | 117 kDa | 0.00088 | KO Synap low, WT Synap high | 0 | 0 | 0 | 11 | 14 | 15 |
| 522 | NADH dehydrogenase [ubiquinone] 1 alpha subcomplex subunit 10, mitochondrial OS=Mus musculus GN=Ndufa10 PE=1 SV=1 | NDUAA_MOUSE | Ndufa10 | 41 kDa | 0.00091 | KO Synap low, WT Synap high | 0 | 0 | 0 | 9 | 11 | 11 |

| | | | | | | | | | | | | |
|---|---|---|---|---|---|---|---|---|---|---|---|---|
| 523 | Aldehyde dehydrogenase, mitochondrial OS=Mus musculus GN=Aldh2 PE=1 SV=1 | ALDH2_MOUSE | Aldh2 | 57 kDa | 0.00064 | KO Synap low, WT Synap high | 0 | 0 | 0 | 16 | 15 | 12 |
| 524 | Proline-rich transmembrane protein 2 OS=Mus musculus GN=Prrt2 PE=1 SV=1 | PRRT2_MOUSE | Prrt2 | 36 kDa | 0.016 | KO Synap low, WT Synap high | 2 | 5 | 6 | 7 | 7 | 9 |
| 525 | Trifunctional enzyme subunit alpha, mitochondrial OS=Mus musculus GN=Hadha PE=1 SV=1 | ECHA_MOUSE | Hadha | 83 kDa | < 0.00010 | KO Synap low, WT Synap high | 0 | 0 | 0 | 11 | 13 | 9 |
| 526 | Sodium/potassium-transporting ATPase subunit beta-1 OS=Mus musculus GN=Atp1b1 PE=1 SV=1 | AT1B1_MOUSE | Atp1b1 | 35 kDa | 0.009 | KO Synap low, WT Synap high | 6 | 7 | 7 | 8 | 6 | 7 |
| 527 | Immunoglobulin superfamily member 8 OS=Mus musculus GN=Igsf8 PE=1 SV=1 | A0A0R4J117_MOUSE (+2) | Igsf8 | 65 kDa | 0.00061 | KO Synap low, WT Synap high | 5 | 6 | 7 | 9 | 8 | 10 |
| 528 | Septin-11 OS=Mus musculus GN=Sept11 PE=1 SV=4 | SEP11_MOUSE (+2) | 11-Sep | 50 kDa | 0.0024 | KO Synap low, WT Synap high | 4 | 3 | 4 | 11 | 12 | 7 |
| 529 | Tubulin polymerization-promoting protein OS=Mus musculus GN=Tppp PE=1 SV=1 | TPPP_MOUSE | Tppp | 24 kDa | 0.003 | KO Synap low, WT Synap high | 5 | 6 | 1 | 12 | 9 | 9 |
| 530 | Sodium- and chloride-dependent GABA transporter 3 OS=Mus musculus GN=Slc6a11 PE=1 SV=2 | S6A11_MOUSE | Slc6a11 | 70 kDa | 0.004 | KO Synap low, WT Synap high | 5 | 4 | 5 | 8 | 7 | 6 |
| 531 | Cytochrome b-c1 complex subunit 2, mitochondrial OS=Mus musculus GN=Uqcrc2 PE=1 SV=1 | QCR2_MOUSE | Uqcrc2 | 48 kDa | 0.001 | KO Synap low, WT Synap high | 0 | 1 | 0 | 9 | 12 | 11 |
| 532 | Neurofascin OS=Mus musculus GN=Nfasc PE=1 SV=1 | NFASC_MOUSE | Nfasc | 138 kDa | < 0.00010 | KO Synap low, WT Synap high | 3 | 2 | 2 | 13 | 14 | 15 |
| 533 | SRC kinase-signaling inhibitor 1 OS=Mus musculus GN=Srcin1 PE=1 SV=1 | B1AQX9_MOUSE | Srcin1 | 131 kDa | 0.00024 | KO Synap low, WT Synap high | 1 | 2 | 0 | 19 | 15 | 18 |
| 534 | Fumarate hydratase, mitochondrial OS=Mus musculus GN=Fh PE=1 SV=3 | FUMH_MOUSE | Fh | 54 kDa | 0.0005 | KO Synap low, WT Synap high | 0 | 0 | 1 | 8 | 9 | 8 |
| 535 | Complexin-2 OS=Mus musculus GN=Cplx2 PE=1 SV=1 | CPLX2_MOUSE | Cplx2 | 15 kDa | 0.014 | KO Synap low, WT Synap high | 6 | 6 | 3 | 4 | 5 | 5 |
| 536 | G protein-regulated inducer of neurite outgrowth 1 OS=Mus musculus GN=Gprin1 PE=1 SV=2 | GRIN1_MOUSE | Gprin1 | 95 kDa | 0.0011 | KO Synap low, WT Synap high | 7 | 7 | 10 | 12 | 14 | 14 |
| 537 | Protein Rab1a OS=Mus musculus GN=Rab1a PE=1 SV=1 | Q5SW88_MOUSE (+1) | Rab1a | 22 kDa | 0.045 | KO Synap low, WT Synap high | 7 | 5 | 5 | 5 | 5 | 4 |
| 538 | 10 kDa heat shock protein, mitochondrial OS=Mus musculus GN=Hspe1 PE=1 SV=2 | CH10_MOUSE | Hspe1 | 11 kDa | 0.00024 | KO Synap low, WT Synap high | 1 | 0 | 0 | 8 | 7 | 9 |
| 539 | NADH dehydrogenase [ubiquinone] 1 alpha subcomplex subunit 9, mitochondrial OS=Mus musculus GN=Ndufa9 PE=1 SV=1 | A0A0R3P9C8_MOUSE (+1) | Ndufa9 | 42 kDa | 0.00012 | KO Synap low, WT Synap high | 0 | 0 | 0 | 11 | 11 | 11 |
| 540 | Cytochrome c, somatic OS=Mus musculus GN=Cycs PE=1 SV=2 | CYC_MOUSE | Cycs | 12 kDa | < 0.00010 | KO Synap low, WT Synap high | 0 | 0 | 0 | 7 | 7 | 6 |
| 541 | NADH dehydrogenase [ubiquinone] iron-sulfur protein 2, mitochondrial OS=Mus musculus GN=Ndufs2 PE=1 SV=1 | NDUS2_MOUSE | Ndufs2 | 53 kDa | 0.0051 | KO Synap low, WT Synap high | 0 | 0 | 0 | 4 | 6 | 9 |
| 542 | Putative adenosylhomocysteinase 2 OS=Mus musculus GN=Ahcyl1 PE=1 SV=1 | SAHH2_MOUSE | Ahcyl1 | 59 kDa | 0.0061 | KO Synap low, WT Synap high | 8 | 2 | 8 | 9 | 11 | 10 |
| 543 | Ras/Rap GTPase-activating protein SynGAP (Fragment) OS=Mus musculus GN=Syngap1 PE=1 SV=2 | A0A0A6YVS6_MOUSE (+3) | Syngap1 | 142 kDa | 0.00035 | KO Synap low, WT Synap high | 3 | 4 | 0 | 9 | 12 | 10 |
| 544 | Septin-5 OS=Mus musculus GN=Sept5 PE=1 SV=2 | SEPT5_MOUSE | 5-Sep | 43 kDa | 0.0022 | KO Synap low, WT Synap high | 2 | 1 | 2 | 8 | 7 | 7 |
| 545 | Guanine nucleotide-binding protein G(I)/G(S)/G(T) subunit beta-2 OS=Mus musculus GN=Gnb2 PE=1 SV=1 | E9QKR0_MOUSE (+1) | Gnb2 | 41 kDa | 0.00033 | KO Synap low, WT Synap high | 5 | 5 | 5 | 7 | 7 | 7 |
| 546 | Cytochrome b-c1 complex subunit 1, mitochondrial OS=Mus musculus GN=Uqcrc1 PE=1 SV=2 | QCR1_MOUSE | Uqcrc1 | 53 kDa | 0.00038 | KO Synap low, WT Synap high | 0 | 2 | 2 | 5 | 8 | 9 |
| 547 | Septin-6 OS=Mus musculus GN=Sept6 PE=1 SV=4 | SEPT6_MOUSE | 6-Sep | 50 kDa | 0.00021 | KO Synap low, WT Synap high | 3 | 3 | 3 | 5 | 7 | 6 |
| 548 | NADH dehydrogenase [ubiquinone] 1 alpha subcomplex subunit 12 OS=Mus musculus GN=Ndufa12 PE=1 SV=1 | A0A0R4J275_MOUSE (+1) | Ndufa12 | 18 kDa | < 0.00010 | KO Synap low, WT Synap high | 0 | 0 | 0 | 8 | 8 | 10 |
| 549 | Actin-related protein 2 OS=Mus musculus GN=Actr2 PE=1 SV=1 | ARP2_MOUSE | Actr2 | 45 kDa | 0.048 | KO Synap low, WT Synap high | 6 | 7 | 6 | 7 | 5 | 8 |
| 550 | Clathrin coat assembly protein AP180 OS=Mus musculus GN=Snap91 PE=1 SV=1 | AP180_MOUSE | Snap91 | 92 kDa | 0.03 | KO Synap low, WT Synap high | 5 | 3 | 5 | 6 | 6 | 9 |
| 551 | NADH dehydrogenase [ubiquinone] flavoprotein 2, mitochondrial OS=Mus musculus GN=Ndufv2 PE=1 SV=2 | NDUV2_MOUSE | Ndufv2 | 27 kDa | < 0.00010 | KO Synap low, WT Synap high | 0 | 0 | 0 | 8 | 7 | 5 |
| 552 | NADH dehydrogenase [ubiquinone] iron-sulfur protein 3, mitochondrial OS=Mus musculus GN=Ndufs3 PE=1 SV=2 | NDUS3_MOUSE | Ndufs3 | 30 kDa | < 0.00010 | KO Synap low, WT Synap high | 0 | 0 | 0 | 9 | 11 | 11 |
| 553 | Dihydrolipoyllysine-residue acetyltransferase component of pyruvate dehydrogenase complex, mitochondrial OS=Mus musculus GN=Dlat PE=1 SV=2 | ODP2_MOUSE | Dlat | 68 kDa | 0.00045 | KO Synap low, WT Synap high | 0 | 0 | 0 | 8 | 8 | 12 |
| 554 | Dihydrolipoyl dehydrogenase, mitochondrial OS=Mus musculus GN=Dld PE=1 SV=2 | DLDH_MOUSE | Dld | 54 kDa | < 0.00010 | KO Synap low, WT Synap high | 0 | 0 | 0 | 10 | 12 | 11 |
| 555 | Cathepsin B OS=Mus musculus GN=Ctsb PE=1 SV=2 | CATB_MOUSE | Ctsb | 37 kDa | 0.014 | KO Synap low, WT Synap high | 2 | 3 | 3 | 9 | 6 | 9 |
| 556 | Succinate--CoA ligase [ADP/GDP-forming] subunit alpha, mitochondrial OS=Mus musculus GN=Suclg1 PE=1 SV=4 | SUCA_MOUSE | Suclg1 | 36 kDa | 0.00028 | KO Synap low, WT Synap high | 2 | 2 | 0 | 6 | 5 | 6 |
| 557 | Methylmalonate-semialdehyde dehydrogenase [acylating], mitochondrial OS=Mus musculus GN=Aldh6a1 PE=1 SV=1 | MMSA_MOUSE | Aldh6a1 | 58 kDa | < 0.00010 | KO Synap low, WT Synap high | 0 | 0 | 0 | 8 | 7 | 9 |
| 558 | Calcium-transporting ATPase OS=Mus musculus GN=Atp2b3 PE=1 SV=1 | A2ALL9_MOUSE (+2) | Atp2b3 | 129 kDa | < 0.00010 | KO Synap low, WT Synap high | 3 | 5 | 6 | 12 | 12 | 14 |
| 559 | Isocitrate dehydrogenase [NADP], mitochondrial OS=Mus musculus GN=Idh2 PE=1 SV=3 | IDHP_MOUSE | Idh2 | 51 kDa | < 0.00010 | KO Synap low, WT Synap high | 0 | 0 | 0 | 8 | 7 | 9 |
| 560 | Capping protein (Actin filament) muscle Z-line, beta, isoform CRA_a OS=Mus musculus GN=Capzb PE=1 SV=1 | A2AMW0_MOUSE | Capzb | 29 kDa | 0.014 | KO Synap low, WT Synap high | 6 | 5 | 6 | 4 | 6 | 5 |
| 561 | MICOS complex subunit Mic19 OS=Mus musculus GN=Chchd3 PE=1 SV=1 | MIC19_MOUSE | Chchd3 | 26 kDa | 0.00085 | KO Synap low, WT Synap high | 0 | 0 | 0 | 13 | 10 | 12 |
| 562 | Electron transfer flavoprotein subunit alpha, mitochondrial OS=Mus musculus GN=Etfa PE=1 SV=2 | ETFA_MOUSE | Etfa | 35 kDa | 0.00047 | KO Synap low, WT Synap high | 0 | 0 | 0 | 7 | 6 | 11 |
| 563 | Transforming protein RhoA OS=Mus musculus GN=Rhoa PE=1 SV=1 | RHOA_MOUSE | Rhoa | 22 kDa | 0.00084 | KO Synap low, WT Synap high | 4 | 3 | 3 | 6 | 6 | 9 |
| 564 | Solute carrier family 12 member 5 OS=Mus musculus GN=Slc12a5 PE=1 SV=2 | S12A5_MOUSE | Slc12a5 | 126 kDa | < 0.00010 | KO Synap low, WT Synap high | 2 | 3 | 3 | 7 | 9 | 9 |
| 565 | Inositol 1,4,5-trisphosphate receptor type 1 OS=Mus musculus GN=Itpr1 PE=1 SV=2 | ITPR1_MOUSE | Itpr1 | 313 kDa | 0.00031 | KO Synap low, WT Synap high | 6 | 3 | 4 | 11 | 13 | 12 |
| 566 | AP-2 complex subunit alpha-1 OS=Mus musculus GN=Ap2a1 PE=1 SV=1 | AP2A1_MOUSE | Ap2a1 | 108 kDa | 0.031 | KO Synap low, WT Synap high | 4 | 7 | 3 | 6 | 11 | 9 |
| 567 | Synaptic vesicle glycoprotein 2A OS=Mus musculus GN=Sv2a PE=1 SV=1 | SV2A_MOUSE | Sv2a | 83 kDa | 0.0065 | KO Synap low, WT Synap high | 7 | 5 | 6 | 9 | 9 | 9 |
| 568 | LETM1 and EF-hand domain-containing protein 1, mitochondrial OS=Mus musculus GN=Letm1 PE=1 SV=1 | LETM1_MOUSE | Letm1 | 83 kDa | 0.0002 | KO Synap low, WT Synap high | 0 | 0 | 0 | 9 | 11 | 9 |
| 569 | Sodium- and chloride-dependent GABA transporter 1 OS=Mus musculus GN=Slc6a1 PE=1 SV=2 | SC6A1_MOUSE | Slc6a1 | 67 kDa | 0.00017 | KO Synap low, WT Synap high | 3 | 2 | 1 | 3 | 3 | 3 |
| 570 | Electron transfer flavoprotein subunit beta OS=Mus musculus GN=Etfb PE=1 SV=3 | ETFB_MOUSE | Etfb | 28 kDa | 0.00047 | KO Synap low, WT Synap high | 0 | 0 | 0 | 7 | 8 | 8 |
| 571 | NADH dehydrogenase [ubiquinone] 1 alpha subcomplex subunit 8 OS=Mus musculus GN=Ndufa8 PE=1 SV=3 | NDUA8_MOUSE | Ndufa8 | 20 kDa | 0.00041 | KO Synap low, WT Synap high | 0 | 0 | 0 | 5 | 5 | 5 |
| 572 | Phosphate carrier protein, mitochondrial OS=Mus musculus GN=Slc25a3 PE=1 SV=1 | MPCP_MOUSE | Slc25a3 | 40 kDa | < 0.00010 | KO Synap low, WT Synap high | 0 | 0 | 0 | 7 | 6 | 7 |
| 573 | Limbic system-associated membrane protein OS=Mus musculus GN=Lsamp PE=1 SV=1 | A0A087WP80_MOUSE (+2) | Lsamp | 39 kDa | 0.0006 | KO Synap low, WT Synap high | 0 | 2 | 2 | 5 | 5 | 8 |
| 574 | ATP synthase subunit d, mitochondrial OS=Mus musculus GN=Atp5h PE=1 SV=3 | ATP5H_MOUSE (+1) | Atp5h | 19 kDa | 0.002 | KO Synap low, WT Synap high | 0 | 0 | 0 | 6 | 10 | 10 |
| 575 | Transgelin-3 OS=Mus musculus GN=Tagln3 PE=1 SV=1 | TAGL3_MOUSE | Tagln3 | 22 kDa | 0.0006 | KO Synap low, WT Synap high | 4 | 5 | 4 | 6 | 8 | 7 |
| 576 | Cytochrome c oxidase subunit NDUFA4 OS=Mus musculus GN=Ndufa4 PE=1 SV=2 | NDUA4_MOUSE (+1) | Ndufa4 | 9 kDa | 0.0026 | KO Synap low, WT Synap high | 0 | 0 | 0 | 10 | 9 | 9 |
| 577 | AP-2 complex subunit alpha-2 OS=Mus musculus GN=Ap2a2 PE=1 SV=2 | AP2A2_MOUSE | Ap2a2 | 104 kDa | 0.01 | KO Synap low, WT Synap high | 5 | 4 | 4 | 6 | 7 | 9 |
| 578 | Complement component 1 Q subcomponent-binding protein, mitochondrial OS=Mus musculus GN=C1qbp PE=1 SV=1 | Q8R5L1_MOUSE | C1qbp | 31 kDa | 0.0007 | KO Synap low, WT Synap high | 0 | 0 | 0 | 5 | 4 | 5 |
| 579 | NADH dehydrogenase [ubiquinone] iron-sulfur protein 6, mitochondrial OS=Mus musculus GN=Ndufs6 PE=1 SV=2 | NDUS6_MOUSE | Ndufs6 | 13 kDa | < 0.00010 | KO Synap low, WT Synap high | 0 | 0 | 0 | 8 | 8 | 9 |
| 580 | RAS-related C3 botulinum substrate 1, isoform CRA_a OS=Mus musculus GN=Rac1 PE=1 SV=1 | Q3TLP8_MOUSE (+1) | Rac1 | 23 kDa | < 0.00010 | KO Synap low, WT Synap high | 3 | 3 | 3 | 5 | 6 | 6 |
| 581 | Cytochrome b-c1 complex subunit 6, mitochondrial OS=Mus musculus GN=Uqcrh PE=1 SV=2 | QCR6_MOUSE | Uqcrh | 10 kDa | 0.0001 | KO Synap low, WT Synap high | 0 | 0 | 0 | 4 | 3 | 3 |
| 582 | Myelin proteolipid protein OS=Mus musculus GN=Plp1 PE=1 SV=2 | MYPR_MOUSE | Plp1 | 30 kDa | 0.00052 | KO Synap low, WT Synap high | 2 | 3 | 2 | 2 | 3 | 3 |
| 583 | Contactin-1 OS=Mus musculus GN=Cntn1 PE=1 SV=1 | CNTN1_MOUSE | Cntn1 | 113 kDa | 0.00016 | KO Synap low, WT Synap high | 2 | 1 | 1 | 7 | 8 | 11 |
| 584 | Neuronal-specific septin-3 OS=Mus musculus GN=Sept3 PE=1 SV=2 | SEPT3_MOUSE | 3-Sep | 40 kDa | 0.00096 | KO Synap low, WT Synap high | 2 | 3 | 2 | 6 | 9 | 8 |
| 585 | ATP synthase F(0) complex subunit B1, mitochondrial OS=Mus musculus GN=Atp5f1 PE=1 SV=1 | AT5F1_MOUSE | Atp5f1 | 29 kDa | 0.00022 | KO Synap low, WT Synap high | 0 | 1 | 1 | 7 | 7 | 6 |
| 586 | Caskin-1 OS=Mus musculus GN=Caskin1 PE=1 SV=2 | CSKI1_MOUSE | Caskin1 | 150 kDa | 0.0025 | KO Synap low, WT Synap high | 2 | 3 | 2 | 12 | 11 | 10 |
| 587 | 3-hydroxyacyl-CoA dehydrogenase type-2 OS=Mus musculus GN=Hsd17b10 PE=1 SV=1 | A2AFQ2_MOUSE (+1) | Hsd17b10 | 28 kDa | 0.00025 | KO Synap low, WT Synap high | 0 | 0 | 0 | 5 | 7 | 6 |
| 588 | Mitochondrial glutamate carrier 1 OS=Mus musculus GN=Slc25a22 PE=1 SV=1 | GHC1_MOUSE | Slc25a22 | 35 kDa | < 0.00010 | KO Synap low, WT Synap high | 0 | 0 | 0 | 7 | 7 | 6 |
| 589 | Syntaxin-1A OS=Mus musculus GN=Stx1a PE=1 SV=1 | D6RFB9_MOUSE (+1) | Stx1a | 29 kDa | < 0.00010 | KO Synap low, WT Synap high | 2 | 2 | 3 | 6 | 7 | 7 |
| 590 | Septin-8 OS=Mus musculus GN=Sept8 PE=1 SV=1 | B1AQZ0_MOUSE | 8-Sep | 56 kDa | 0.0014 | KO Synap low, WT Synap high | 0 | 0 | 0 | 7 | 9 | 8 |
| 591 | 14-3-3 protein gamma OS=Mus musculus GN=Ywhag PE=1 SV=2 | 1433G_MOUSE | Ywhag | 28 kDa | 0.037 | KO Synap low, WT Synap high | 3 | 3 | 4 | 4 | 7 | 6 |
| 592 | NADH dehydrogenase [ubiquinone] 1 alpha subcomplex subunit 7 OS=Mus musculus GN=Ndufa7 PE=1 SV=3 | NDUA7_MOUSE | Ndufa7 | 13 kDa | 0.0043 | KO Synap low, WT Synap high | 0 | 0 | 0 | 9 | 9 | 10 |
| 593 | ATP synthase subunit gamma OS=Mus musculus GN=Atp5c1 PE=1 SV=1 | A2AKU9_MOUSE (+2) | Atp5c1 | 33 kDa | 0.0031 | KO Synap low, WT Synap high | 0 | 0 | 0 | 7 | 6 | 8 |
| 594 | D-beta-hydroxybutyrate dehydrogenase, mitochondrial OS=Mus musculus GN=Bdh1 PE=1 SV=2 | BDH_MOUSE | Bdh1 | 38 kDa | 0.00051 | KO Synap low, WT Synap high | 0 | 0 | 0 | 7 | 7 | 6 |
| 595 | Plectin OS=Mus musculus GN=Plec PE=1 SV=3 | PLEC_MOUSE | Plec | 534 kDa | 0.012 | KO Synap low, WT Synap high | 6 | 1 | 1 | 12 | 9 | 10 |
| 596 | Disks large homolog 4 OS=Mus musculus GN=Dlg4 PE=1 SV=1 | DLG4_MOUSE | Dlg4 | 80 kDa | < 0.00010 | KO Synap low, WT Synap high | 0 | 1 | 0 | 8 | 8 | 9 |
| 597 | Sideroflexin-3 OS=Mus musculus GN=Sfxn3 PE=1 SV=1 | SFXN3_MOUSE | Sfxn3 | 35 kDa | < 0.00010 | KO Synap low, WT Synap high | 0 | 1 | 0 | 5 | 7 | 5 |
| 598 | NADH dehydrogenase [ubiquinone] 1 beta subcomplex subunit 4 OS=Mus musculus GN=Ndufb4 PE=1 SV=3 | NDUB4_MOUSE | Ndufb4 | 15 kDa | 0.00049 | KO Synap low, WT Synap high | 0 | 0 | 0 | 7 | 6 | 5 |
| 599 | Protein NipSnap homolog 2 OS=Mus musculus GN=Gbas PE=1 SV=1 | NIPS2_MOUSE | Gbas | 33 kDa | < 0.00010 | KO Synap low, WT Synap high | 0 | 0 | 0 | 5 | 5 | 6 |
| 600 | Succinate-semialdehyde dehydrogenase, mitochondrial OS=Mus musculus GN=Aldh5a1 PE=1 SV=1 | SSDH_MOUSE | Aldh5a1 | 56 kDa | < 0.00010 | KO Synap low, WT Synap high | 0 | 0 | 0 | 5 | 6 | 6 |
| 601 | Nucleoside diphosphate kinase A OS=Mus musculus GN=Nme1 PE=1 SV=1 | NDKA_MOUSE | Nme1 | 17 kDa | 0.011 | KO Synap low, WT Synap high | 0 | 2 | 3 | 3 | 5 | 3 |
| 602 | NADH dehydrogenase [ubiquinone] 1 alpha subcomplex subunit 5 OS=Mus musculus GN=Ndufa5 PE=1 SV=3 | NDUA5_MOUSE | Ndufa5 | 13 kDa | 0.00046 | KO Synap low, WT Synap high | 0 | 0 | 0 | 4 | 4 | 6 |
| 603 | Cytochrome b-c1 complex subunit Rieske, mitochondrial OS=Mus musculus GN=Uqcrfs1 PE=1 SV=1 | UCRI_MOUSE | Uqcrfs1 | 29 kDa | < 0.00010 | KO Synap low, WT Synap high | 0 | 0 | 0 | 4 | 4 | 4 |
| 604 | NADH dehydrogenase [ubiquinone] iron-sulfur protein 8, mitochondrial OS=Mus musculus GN=Ndufs8 PE=1 SV=1 | NDUS8_MOUSE | Ndufs8 | 24 kDa | 0.0033 | KO Synap low, WT Synap high | 0 | 0 | 0 | 9 | 7 | 9 |
| 605 | NADH dehydrogenase [ubiquinone] 1 alpha subcomplex subunit 13 OS=Mus musculus GN=Ndufa13 PE=1 SV=3 | NDUAD_MOUSE | Ndufa13 | 17 kDa | 0.0032 | KO Synap low, WT Synap high | 0 | 0 | 0 | 5 | 4 | 3 |
| 606 | Thiosulfate sulfurtransferase OS=Mus musculus GN=Tst PE=1 SV=3 | THTR_MOUSE | Tst | 33 kDa | < 0.00010 | KO Synap low, WT Synap high | 0 | 0 | 0 | 7 | 5 | 7 |
| 607 | Vesicle-associated membrane protein-associated protein A OS=Mus musculus GN=Vapa PE=1 SV=2 | VAPA_MOUSE | Vapa | 28 kDa | 0.00091 | KO Synap low, WT Synap high | 2 | 2 | 2 | 4 | 3 | 4 |
| 608 | 3-ketoacyl-CoA thiolase, mitochondrial OS=Mus musculus GN=Acaa2 PE=1 SV=3 | THIM_MOUSE | Acaa2 | 42 kDa | < 0.00010 | KO Synap low, WT Synap high | 0 | 0 | 0 | 5 | 9 | 9 |
| 609 | ADP/ATP translocase 2 OS=Mus musculus GN=Slc25a5 PE=1 SV=3 | ADT2_MOUSE | Slc25a5 | 33 kDa | < 0.00010 | KO Synap low, WT Synap high | 0 | 0 | 0 | 5 | 7 | 8 |
| 610 | Neurofilament light polypeptide OS=Mus musculus GN=Nefl PE=1 SV=5 | NFL_MOUSE | Nefl | 62 kDa | 0.046 | KO Synap low, WT Synap high | 1 | 2 | 1 | 6 | 3 | 4 |
| 611 | NADH dehydrogenase [ubiquinone] 1 alpha subcomplex subunit 6 OS=Mus musculus GN=Ndufa6 PE=1 SV=1 | NDUA6_MOUSE | Ndufa6 | 15 kDa | 0.0014 | KO Synap low, WT Synap high | 1 | 0 | 0 | 3 | 3 | 4 |
| 612 | Cytochrome c oxidase subunit 6B1 OS=Mus musculus GN=Cox6b1 PE=1 SV=2 | CX6B1_MOUSE | Cox6b1 | 10 kDa | 0.0018 | KO Synap low, WT Synap high | 0 | 0 | 0 | 3 | 5 | 5 |
| 613 | NADH dehydrogenase [ubiquinone] iron-sulfur protein 7, mitochondrial OS=Mus musculus GN=Ndufs7 PE=1 SV=1 | NDUS7_MOUSE | Ndufs7 | 25 kDa | < 0.00010 | KO Synap low, WT Synap high | 0 | 0 | 0 | 6 | 5 | 5 |

| | | | | | | | | | | | | |
|---|---|---|---|---|---|---|---|---|---|---|---|---|
| 614 | Trifunctional enzyme subunit beta, mitochondrial OS=Mus musculus GN=Hadhb PE=1 SV=1 | ECHB_MOUSE | Hadhb | 51 kDa | < 0.00010 | KO Synap low, WT Synap high | 0 | 0 | 0 | 4 | 4 | 5 |
| 615 | Serine/threonine-protein phosphatase OS=Mus musculus GN=Ppp3cb PE=1 SV=1 | E0CZ78_MOUSE (+1) | Ppp3cb | 59 kDa | 0.036 | KO Synap low, WT Synap high | 5 | 4 | 3 | 3 | 5 | 4 |
| 616 | Thioredoxin-dependent peroxide reductase, mitochondrial OS=Mus musculus GN=Prdx3 PE=1 SV=1 | PRDX3_MOUSE | Prdx3 | 28 kDa | < 0.00010 | KO Synap low, WT Synap high | 0 | 0 | 0 | 6 | 6 | 6 |
| 617 | Ornithine aminotransferase, mitochondrial OS=Mus musculus GN=Oat PE=1 SV=1 | OAT_MOUSE | Oat | 48 kDa | 0.0087 | KO Synap low, WT Synap high | 0 | 0 | 0 | 8 | 3 | 5 |
| 618 | Lon protease homolog, mitochondrial OS=Mus musculus GN=Lonp1 PE=1 SV=2 | LONM_MOUSE | Lonp1 | 106 kDa | 0.0007 | KO Synap low, WT Synap high | 0 | 0 | 0 | 5 | 7 | 5 |
| 619 | LanC-like protein 2 (Fragment) OS=Mus musculus GN=Lancl2 PE=1 SV=1 | F6RJV6_MOUSE (+1) | Lancl2 | 50 kDa | 0.034 | KO Synap low, WT Synap high | 4 | 2 | 2 | 5 | 6 | 5 |
| 620 | Alpha-internexin OS=Mus musculus GN=Ina PE=1 SV=3 | AINX_MOUSE | Ina | 55 kDa | 0.0013 | KO Synap low, WT Synap high | 1 | 0 | 1 | 6 | 8 | 6 |
| 621 | Guanine nucleotide-binding protein G(i) subunit alpha-2 OS=Mus musculus GN=Gnai2 PE=1 SV=5 | GNAI2_MOUSE | Gnai2 | 40 kDa | 0.0012 | KO Synap low, WT Synap high | 2 | 0 | 2 | 5 | 3 | 6 |
| 622 | ATP synthase subunit e, mitochondrial OS=Mus musculus GN=Atp5i PE=1 SV=2 | ATP5I_MOUSE | Atp5i | 8 kDa | 0.00047 | KO Synap low, WT Synap high | 0 | 0 | 0 | 4 | 3 | 3 |
| 623 | GTPase HRas OS=Mus musculus GN=Hras PE=1 SV=2 | RASH_MOUSE | Hras | 21 kDa | 0.0019 | KO Synap low, WT Synap high | 0 | 2 | 2 | 5 | 5 | 5 |
| 624 | Neurotrimin OS=Mus musculus GN=Ntm PE=1 SV=1 | D3Z396_MOUSE (+2) | Ntm | 35 kDa | 0.0076 | KO Synap low, WT Synap high | 3 | 2 | 1 | 7 | 4 | 3 |
| 625 | Leucine-rich PPR motif-containing protein, mitochondrial OS=Mus musculus GN=Lrpprc PE=1 SV=2 | LPPRC_MOUSE | Lrpprc | 157 kDa | 0.0017 | KO Synap low, WT Synap high | 0 | 0 | 0 | 8 | 6 | 11 |
| 626 | Basigin (Fragment) OS=Mus musculus GN=Bsg PE=1 SV=1 | J3QP71_MOUSE | Bsg | 22 kDa | 0.014 | KO Synap low, WT Synap high | 2 | 3 | 3 | 4 | 3 | 4 |
| 627 | Cytoplasmic FMR1-interacting protein 2 OS=Mus musculus GN=Cyfip2 PE=1 SV=2 | CYFP2_MOUSE | Cyfip2 | 146 kDa | 0.0076 | KO Synap low, WT Synap high | 3 | 3 | 3 | 5 | 3 | 5 |
| 628 | Brain-specific angiogenesis inhibitor 1-associated protein 2 OS=Mus musculus GN=Baiap2 PE=1 SV=1 | B1AZ46_MOUSE (+1) | Baiap2 | 58 kDa | < 0.00010 | KO Synap low, WT Synap high | 1 | 0 | 1 | 8 | 8 | 7 |
| 629 | Propionyl-CoA carboxylase beta chain, mitochondrial OS=Mus musculus GN=Pccb PE=1 SV=2 | PCCB_MOUSE (+1) | Pccb | 58 kDa | 0.002 | KO Synap low, WT Synap high | 0 | 0 | 0 | 6 | 6 | 5 |
| 630 | Oligodendrocyte myelin glycoprotein OS=Mus musculus GN=Omg PE=1 SV=1 | G3XA53_MOUSE (+1) | Omg | 50 kDa | 0.0044 | KO Synap low, WT Synap high | 2 | 2 | 1 | 3 | 4 | 3 |
| 631 | Long chain acyl-CoA synthetase 6 isoform 3 OS=Mus musculus GN=Acsl6 PE=1 SV=1 | Q5ICG5_MOUSE (+1) | Acsl6 | 78 kDa | 0.0017 | KO Synap low, WT Synap high | 2 | 1 | 2 | 5 | 4 | 6 |
| 632 | Visinin-like protein 1 OS=Mus musculus GN=Vsnl1 PE=1 SV=2 | VISL1_MOUSE | Vsnl1 | 22 kDa | 0.0015 | KO Synap low, WT Synap high | 0 | 1 | 0 | 4 | 6 | 5 |
| 633 | AFG3-like protein 2 OS=Mus musculus GN=Afg3l2 PE=1 SV=1 | AFG32_MOUSE | Afg3l2 | 90 kDa | 0.00019 | KO Synap low, WT Synap high | 0 | 0 | 0 | 6 | 7 | 5 |
| 634 | Neuronal membrane glycoprotein M6-a OS=Mus musculus GN=Gpm6a PE=1 SV=1 | GPM6A_MOUSE | Gpm6a | 31 kDa | 0.0021 | KO Synap low, WT Synap high | 2 | 2 | 2 | 4 | 5 | 3 |
| 635 | Actin-related protein 2/3 complex subunit 2 OS=Mus musculus GN=Arpc2 PE=1 SV=3 | ARPC2_MOUSE | Arpc2 | 34 kDa | 0.039 | KO Synap low, WT Synap high | 2 | 0 | 3 | 2 | 3 | 3 |
| 636 | ATP synthase-coupling factor 6, mitochondrial OS=Mus musculus GN=Atp5j PE=1 SV=1 | ATP5J_MOUSE | Atp5j | 12 kDa | 0.0048 | KO Synap low, WT Synap high | 0 | 0 | 0 | 4 | 3 | 5 |
| 637 | Enoyl-CoA hydratase, mitochondrial OS=Mus musculus GN=Echs1 PE=1 SV=1 | ECHM_MOUSE | Echs1 | 31 kDa | 0.0027 | KO Synap low, WT Synap high | 0 | 0 | 0 | 4 | 5 | 7 |
| 638 | Cytochrome c1, heme protein, mitochondrial OS=Mus musculus GN=Cyc1 PE=1 SV=1 | CY1_MOUSE | Cyc1 | 35 kDa | 0.016 | KO Synap low, WT Synap high | 0 | 0 | 0 | 2 | 4 | 3 |
| 639 | Guanine nucleotide-binding protein G(I)/G(S)/G(O) subunit gamma-3 OS=Mus musculus GN=Gng3 PE=1 SV=1 | GBG3_MOUSE | Gng3 | 8 kDa | 0.021 | KO Synap low, WT Synap high | 1 | 1 | 1 | 2 | 2 | 2 |
| 640 | Pyruvate dehydrogenase protein X component, mitochondrial OS=Mus musculus GN=Pdhx PE=1 SV=1 | ODPX_MOUSE | Pdhx | 54 kDa | < 0.00010 | KO Synap low, WT Synap high | 0 | 0 | 0 | 4 | 5 | 5 |
| 641 | ES1 protein homolog, mitochondrial OS=Mus musculus GN=D10Jhu81e PE=1 SV=1 | ES1_MOUSE | D10Jhu81e | 28 kDa | 0.0011 | KO Synap low, WT Synap high | 0 | 0 | 0 | 4 | 5 | 4 |
| 642 | Sideroflexin-1 OS=Mus musculus GN=Sfxn1 PE=1 SV=3 | SFXN1_MOUSE | Sfxn1 | 36 kDa | 0.0016 | KO Synap low, WT Synap high | 0 | 0 | 0 | 5 | 6 | 6 |
| 643 | Mitochondrial 2-oxoglutarate/malate carrier protein OS=Mus musculus GN=Slc25a11 PE=1 SV=3 | M2OM_MOUSE | Slc25a11 | 34 kDa | 0.0044 | KO Synap low, WT Synap high | 0 | 0 | 0 | 4 | 5 | 4 |
| 644 | Fumarylacetoacetate hydrolase domain-containing protein 2A OS=Mus musculus GN=Fahd2a PE=1 SV=1 | A0A0R4J094_MOUSE | Fahd2a | 35 kDa | < 0.00010 | KO Synap low, WT Synap high | 0 | 0 | 0 | 3 | 5 | 2 |
| 645 | Ganglioside-induced differentiation-associated protein 1-like 1 OS=Mus musculus GN=Gdap1l1 PE=1 SV=1 | GD1L1_MOUSE (+1) | Gdap1l1 | 42 kDa | 0.043 | KO Synap low, WT Synap high | 0 | 1 | 1 | 2 | 3 | 4 |
| 646 | Methylglutaconyl-CoA hydratase, mitochondrial OS=Mus musculus GN=Auh PE=1 SV=1 | A0A0R4J023_MOUSE (+1) | Auh | 33 kDa | < 0.00010 | KO Synap low, WT Synap high | 0 | 0 | 0 | 5 | 7 | 6 |
| 647 | ATP synthase subunit delta, mitochondrial OS=Mus musculus GN=Atp5d PE=1 SV=1 | ATPD_MOUSE | Atp5d | 18 kDa | 0.0076 | KO Synap low, WT Synap high | 0 | 0 | 0 | 3 | 2 | 3 |
| 648 | Secretogranin-2 OS=Mus musculus GN=Scg2 PE=1 SV=1 | SCG2_MOUSE | Scg2 | 71 kDa | 0.0035 | KO Synap low, WT Synap high | 3 | 2 | 1 | 6 | 6 | 5 |
| 649 | WD repeat-containing protein 7 OS=Mus musculus GN=Wdr7 PE=1 SV=3 | WDR7_MOUSE | Wdr7 | 163 kDa | 0.00044 | KO Synap low, WT Synap high | 0 | 0 | 0 | 5 | 8 | 3 |
| 650 | Very long-chain specific acyl-CoA dehydrogenase, mitochondrial OS=Mus musculus GN=Acadvl PE=1 SV=3 | ACADV_MOUSE | Acadvl | 71 kDa | 0.0013 | KO Synap low, WT Synap high | 0 | 0 | 0 | 6 | 6 | 8 |
| 651 | Propionyl-CoA carboxylase alpha chain, mitochondrial OS=Mus musculus GN=Pcca PE=1 SV=2 | PCCA_MOUSE | Pcca | 80 kDa | < 0.00010 | KO Synap low, WT Synap high | 0 | 0 | 0 | 4 | 4 | 4 |
| 652 | Cytochrome c oxidase subunit 7A2, mitochondrial OS=Mus musculus GN=Cox7a2 PE=1 SV=2 | CX7A2_MOUSE | Cox7a2 | 9 kDa | 0.00046 | KO Synap low, WT Synap high | 0 | 0 | 0 | 3 | 3 | 3 |
| 653 | NADH dehydrogenase [ubiquinone] 1 subunit C2 OS=Mus musculus GN=Ndufc2 PE=1 SV=1 | NDUC2_MOUSE | Ndufc2 | 14 kDa | 0.0044 | KO Synap low, WT Synap high | 0 | 0 | 0 | 4 | 4 | 2 |
| 654 | Calcium-transporting ATPase OS=Mus musculus GN=Atp2b4 PE=1 SV=1 | E9Q828_MOUSE (+1) | Atp2b4 | 129 kDa | 0.00036 | KO Synap low, WT Synap high | 2 | 2 | 2 | 4 | 4 | 5 |
| 655 | CDGSH iron-sulfur domain-containing protein 1 OS=Mus musculus GN=Cisd1 PE=1 SV=1 | CISD1_MOUSE | Cisd1 | 12 kDa | 0.0028 | KO Synap low, WT Synap high | 0 | 1 | 1 | 3 | 3 | 3 |
| 656 | Phospholipase D3 OS=Mus musculus GN=Pld3 PE=1 SV=1 | PLD3_MOUSE | Pld3 | 54 kDa | 0.00014 | KO Synap low, WT Synap high | 0 | 0 | 0 | 3 | 2 | 3 |
| 657 | Leucine-rich glioma-inactivated protein 1 OS=Mus musculus GN=Lgi1 PE=1 SV=1 | LGI1_MOUSE | Lgi1 | 64 kDa | 0.0031 | KO Synap low, WT Synap high | 0 | 0 | 0 | 6 | 6 | 4 |
| 658 | Paralemmin-1 OS=Mus musculus GN=Palm PE=1 SV=1 | PALM_MOUSE | Palm | 42 kDa | 0.024 | KO Synap low, WT Synap high | 1 | 2 | 1 | 4 | 3 | 2 |
| 659 | Catenin alpha-2 OS=Mus musculus GN=Ctnna2 PE=1 SV=3 | CTNA2_MOUSE (+1) | Ctnna2 | 105 kDa | 0.019 | KO Synap low, WT Synap high | 1 | 1 | 0 | 2 | 3 | 5 |
| 660 | Glycerol-3-phosphate dehydrogenase, mitochondrial OS=Mus musculus GN=Gpd2 PE=1 SV=2 | GPDM_MOUSE | Gpd2 | 81 kDa | 0.00049 | KO Synap low, WT Synap high | 0 | 0 | 0 | 4 | 4 | 3 |
| 661 | Cysteine and glycine-rich protein 1 OS=Mus musculus GN=Csrp1 PE=1 SV=3 | CSRP1_MOUSE | Csrp1 | 21 kDa | 0.023 | KO Synap low, WT Synap high | 0 | 0 | 2 | 4 | 3 | 3 |
| 662 | NADH dehydrogenase [ubiquinone] 1 alpha subcomplex subunit 2 OS=Mus musculus GN=Ndufa2 PE=1 SV=3 | NDUA2_MOUSE | Ndufa2 | 11 kDa | 0.0092 | KO Synap low, WT Synap high | 1 | 1 | 1 | 3 | 2 | 3 |
| 663 | Disintegrin and metalloproteinase domain-containing protein 22 OS=Mus musculus GN=Adam22 PE=1 SV=2 | ADA22_MOUSE | Adam22 | 100 kDa | < 0.00010 | KO Synap low, WT Synap high | 0 | 0 | 0 | 5 | 6 | 4 |
| 664 | Uncharacterized protein KIAA0513 OS=Mus musculus GN=Kiaa0513 PE=1 SV=1 | K0513_MOUSE | Kiaa0513 | 46 kDa | 0.0044 | KO Synap low, WT Synap high | 0 | 0 | 2 | 7 | 4 | 5 |
| 665 | ATP synthase subunit g, mitochondrial OS=Mus musculus GN=Atp5l PE=1 SV=1 | ATP5L_MOUSE | Atp5l | 11 kDa | 0.0027 | KO Synap low, WT Synap high | 0 | 0 | 0 | 2 | 2 | 2 |
| 666 | Protein NipSnap homolog 1 OS=Mus musculus GN=Nipsnap1 PE=1 SV=1 | NIPS1_MOUSE | Nipsnap1 | 33 kDa | 0.0058 | KO Synap low, WT Synap high | 0 | 0 | 0 | 4 | 2 | 4 |
| 667 | Catenin delta-2 OS=Mus musculus GN=Ctnnd2 PE=1 SV=1 | CTND2_MOUSE (+1) | Ctnnd2 | 135 kDa | 0.0013 | KO Synap low, WT Synap high | 1 | 1 | 1 | 5 | 4 | 6 |
| 668 | Hepatocyte cell adhesion molecule OS=Mus musculus GN=Hepacam PE=1 SV=2 | HECAM_MOUSE | Hepacam | 46 kDa | 0.043 | KO Synap low, WT Synap high | 2 | 0 | 2 | 2 | 2 | 2 |
| 669 | Neural cell adhesion molecule L1 OS=Mus musculus GN=L1cam PE=1 SV=1 | A2AFG8_MOUSE (+1) | L1cam | 140 kDa | 0.0013 | KO Synap low, WT Synap high | 0 | 0 | 0 | 3 | 4 | 4 |
| 670 | Mitochondrial import receptor subunit TOM22 homolog OS=Mus musculus GN=Tomm22 PE=1 SV=3 | TOM22_MOUSE | Tomm22 | 16 kDa | 0.00044 | KO Synap low, WT Synap high | 0 | 0 | 0 | 3 | 3 | 4 |
| 671 | Metaxin-2 OS=Mus musculus GN=Mtx2 PE=1 SV=1 | MTX2_MOUSE | Mtx2 | 30 kDa | 0.0032 | KO Synap low, WT Synap high | 0 | 0 | 0 | 2 | 4 | 3 |
| 672 | Ankyrin-2 (Fragment) OS=Mus musculus GN=Ank2 PE=1 SV=2 | S4R2T7_MOUSE | Ank2 | 131 kDa | 0.0034 | KO Synap low, WT Synap high | 1 | 0 | 0 | 3 | 3 | 2 |
| 673 | Homer protein homolog 1 OS=Mus musculus GN=Homer1 PE=1 SV=2 | HOME1_MOUSE | Homer1 | 41 kDa | 0.007 | KO Synap low, WT Synap high | 1 | 0 | 0 | 3 | 6 | 3 |
| 674 | Syntaxin-12 OS=Mus musculus GN=Stx12 PE=1 SV=1 | STX12_MOUSE | Stx12 | 31 kDa | < 0.00010 | KO Synap low, WT Synap high | 0 | 0 | 0 | 3 | 4 | 4 |
| 675 | Mitochondrial import inner membrane translocase subunit Tim9 OS=Mus musculus GN=Timm9 PE=1 SV=1 | TIM9_MOUSE | Timm9 | 10 kDa | 0.0086 | KO Synap low, WT Synap high | 0 | 0 | 0 | 4 | 3 | 4 |
| 676 | EH domain-containing protein 3 OS=Mus musculus GN=Ehd3 PE=1 SV=2 | EHD3_MOUSE | Ehd3 | 61 kDa | 0.031 | KO Synap low, WT Synap high | 2 | 1 | 0 | 3 | 5 | 2 |
| 677 | Rho-related GTP-binding protein RhoB OS=Mus musculus GN=Rhob PE=1 SV=1 | RHOB_MOUSE | Rhob | 22 kDa | 0.017 | KO Synap low, WT Synap high | 1 | 1 | 1 | 4 | 3 | 4 |
| 678 | Alpha-actinin-1 OS=Mus musculus GN=Actn1 PE=1 SV=1 | ACTN1_MOUSE | Actn1 | 103 kDa | 0.013 | KO Synap low, WT Synap high | 1 | 1 | 0 | 3 | 3 | 3 |
| 679 | Voltage-dependent anion-selective channel protein 1 OS=Mus musculus GN=Vdac1 PE=1 SV=3 | VDAC1_MOUSE | Vdac1 | 32 kDa | 0.0029 | KO Synap low, WT Synap high | 0 | 1 | 0 | 2 | 3 | 3 |
| 680 | Succinate dehydrogenase [ubiquinone] iron-sulfur subunit, mitochondrial OS=Mus musculus GN=Sdhb PE=1 SV=1 | SDHB_MOUSE | Sdhb | 32 kDa | 0.0028 | KO Synap low, WT Synap high | 0 | 0 | 0 | 3 | 6 | 4 |
| 681 | Talin-2 OS=Mus musculus GN=Tln2 PE=1 SV=1 | E9PUM4_MOUSE (+1) | Tln2 | 272 kDa | 0.036 | KO Synap low, WT Synap high | 3 | 1 | 1 | 3 | 5 | 4 |
| 682 | Breast carcinoma-amplified sequence 1 homolog OS=Mus musculus GN=Bcas1 PE=1 SV=3 | BCAS1_MOUSE (+1) | Bcas1 | 67 kDa | 0.0039 | KO Synap low, WT Synap high | 1 | 1 | 0 | 5 | 3 | 4 |
| 683 | MICOS complex subunit Mic25 OS=Mus musculus GN=Chchd6 PE=1 SV=2 | MIC25_MOUSE | Chchd6 | 30 kDa | 0.011 | KO Synap low, WT Synap high | 0 | 0 | 0 | 8 | 6 | 3 |
| 684 | C2 domain-containing protein 2-like OS=Mus musculus GN=C2cd2l PE=1 SV=3 | C2C2L_MOUSE | C2cd2l | 76 kDa | 0.0047 | KO Synap low, WT Synap high | 0 | 2 | 0 | 3 | 5 | 4 |
| 685 | Mitochondrial amidoxime reducing component 2 OS=Mus musculus GN=Marc2 PE=1 SV=1 | MARC2_MOUSE | 2-Mar | 38 kDa | 0.0024 | KO Synap low, WT Synap high | 0 | 0 | 0 | 4 | 5 | 4 |
| 686 | NADH dehydrogenase [ubiquinone] flavoprotein 3, mitochondrial OS=Mus musculus GN=Ndufv3 PE=1 SV=1 | NDUV3_MOUSE | Ndufv3 | 12 kDa | 0.00028 | KO Synap low, WT Synap high | 0 | 0 | 0 | 1 | 1 | 1 |
| 687 | Kinesin-like protein OS=Mus musculus GN=Kif2a PE=1 SV=1 | E0CZ72_MOUSE (+1) | Kif2a | 84 kDa | 0.0052 | KO Synap low, WT Synap high | 2 | 0 | 0 | 3 | 4 | 4 |
| 688 | Acyl-CoA dehydrogenase family member 9, mitochondrial OS=Mus musculus GN=Acad9 PE=1 SV=2 | ACAD9_MOUSE | Acad9 | 69 kDa | 0.005 | KO Synap low, WT Synap high | 0 | 0 | 0 | 5 | 6 | 3 |
| 689 | Synaptic vesicle glycoprotein 2B OS=Mus musculus GN=Sv2b PE=1 SV=1 | SV2B_MOUSE | Sv2b | 77 kDa | 0.031 | KO Synap low, WT Synap high | 0 | 0 | 0 | 2 | 3 | 7 |
| 690 | Inorganic pyrophosphatase 2, mitochondrial OS=Mus musculus GN=Ppa2 PE=1 SV=1 | D3Z636_MOUSE (+1) | Ppa2 | 38 kDa | 0.0026 | KO Synap low, WT Synap high | 0 | 0 | 1 | 3 | 3 | 4 |
| 691 | Prohibitin-2 OS=Mus musculus GN=Phb2 PE=1 SV=1 | PHB2_MOUSE | Phb2 | 33 kDa | 0.0013 | KO Synap low, WT Synap high | 0 | 0 | 0 | 3 | 5 | 5 |
| 692 | Protein Pcdh1 OS=Mus musculus GN=Pcdh1 PE=1 SV=1 | Q8CFX3_MOUSE | Pcdh1 | 112 kDa | 0.0059 | KO Synap low, WT Synap high | 0 | 0 | 0 | 5 | 4 | 3 |
| 693 | Succinate--CoA ligase [GDP-forming] subunit beta, mitochondrial OS=Mus musculus GN=Suclg2 PE=1 SV=3 | SUCB2_MOUSE | Suclg2 | 47 kDa | 0.0059 | KO Synap low, WT Synap high | 0 | 0 | 0 | 5 | 3 | 3 |
| 694 | Disintegrin and metalloproteinase domain-containing protein 23 OS=Mus musculus GN=Adam23 PE=1 SV=1 | ADA23_MOUSE (+1) | Adam23 | 92 kDa | 0.00051 | KO Synap low, WT Synap high | 0 | 0 | 0 | 4 | 4 | 4 |
| 695 | Vesicular inhibitory amino acid transporter OS=Mus musculus GN=Slc32a1 PE=1 SV=3 | VIAAT_MOUSE | Slc32a1 | 57 kDa | 0.015 | KO Synap low, WT Synap high | 0 | 0 | 0 | 3 | 2 | 2 |
| 696 | Neurofilament heavy polypeptide OS=Mus musculus GN=Nefh PE=1 SV=3 | NFH_MOUSE | Nefh | 117 kDa | 0.0059 | KO Synap low, WT Synap high | 1 | 0 | 0 | 4 | 3 | 2 |
| 697 | Ganglioside-induced differentiation-associated protein 1 OS=Mus musculus GN=Gdap1 PE=1 SV=1 | GDAP1_MOUSE | Gdap1 | 41 kDa | 0.00044 | KO Synap low, WT Synap high | 0 | 0 | 0 | 4 | 6 | 6 |
| 698 | Mitochondrial glutamate carrier 2 OS=Mus musculus GN=Slc25a18 PE=1 SV=4 | GHC2_MOUSE | Slc25a18 | 34 kDa | 0.00085 | KO Synap low, WT Synap high | 0 | 0 | 0 | 3 | 3 | 4 |
| 699 | Casein kinase II subunit alpha OS=Mus musculus GN=Csnk2a1 PE=1 SV=2 | CSK21_MOUSE | Csnk2a1 | 45 kDa | 0.03 | KO Synap low, WT Synap high | 0 | 0 | 0 | 1 | 2 | 4 |
| 700 | 3-hydroxyisobutyryl-CoA hydrolase, mitochondrial OS=Mus musculus GN=Hibch PE=1 SV=1 | HIBCH_MOUSE | Hibch | 43 kDa | 0.019 | KO Synap low, WT Synap high | 0 | 0 | 0 | 2 | 1 | 3 |
| 701 | Acyl-coenzyme A thioesterase 13 OS=Mus musculus GN=Acot13 PE=1 SV=1 | ACO13_MOUSE | Acot13 | 15 kDa | 0.0085 | KO Synap low, WT Synap high | 0 | 0 | 0 | 1 | 1 | 3 |
| 702 | Homer protein homolog 3 OS=Mus musculus GN=Homer3 PE=1 SV=2 | HOME3_MOUSE | Homer3 | 40 kDa | 0.0073 | KO Synap low, WT Synap high | 0 | 1 | 1 | 3 | 2 | 4 |
| 703 | Phospholemman OS=Mus musculus GN=Fxyd1 PE=1 SV=1 | A0A0U1RPI1_MOUSE (+1) | Fxyd1 | 8 kDa | 0.014 | KO Synap low, WT Synap high | 1 | 1 | 1 | 1 | 1 | 1 |
| 704 | Adenylate kinase 4, mitochondrial OS=Mus musculus GN=Ak4 PE=1 SV=1 | KAD4_MOUSE | Ak4 | 25 kDa | 0.0043 | KO Synap low, WT Synap high | 0 | 0 | 0 | 3 | 2 | 3 |

| | | | | | | | | | | | | |
|---|---|---|---|---|---|---|---|---|---|---|---|---|
| 705 | NADH dehydrogenase [ubiquinone] flavoprotein 3, mitochondrial OS=Mus musculus GN=Ndufv3 PE=1 SV=1 | Q3U422_MOUSE | Ndufv3 | 50 kDa | < 0.00010 | KO Synap low, WT Synap high | 0 | 0 | 0 | 3 | 2 | 3 |
| 706 | Alpha-soluble NSF attachment protein OS=Mus musculus GN=Napa PE=1 SV=1 | SNAA_MOUSE | Napa | 33 kDa | 0.023 | KO Synap low, WT Synap high | 2 | 0 | 1 | 2 | 3 | 4 |
| 707 | Guanine nucleotide-binding protein G(i) subunit alpha-1 OS=Mus musculus GN=Gnai1 PE=1 SV=1 | GNAI1_MOUSE | Gnai1 | 40 kDa | 0.0012 | KO Synap low, WT Synap high | 1 | 1 | 1 | 1 | 2 | 2 |
| 708 | Intercellular adhesion molecule 5 OS=Mus musculus GN=Icam5 PE=1 SV=2 | ICAM5_MOUSE | Icam5 | 97 kDa | 0.00024 | KO Synap low, WT Synap high | 1 | 0 | 0 | 4 | 4 | 4 |
| 709 | Heat shock protein 75 kDa, mitochondrial OS=Mus musculus GN=Trap1 PE=1 SV=1 | TRAP1_MOUSE | Trap1 | 80 kDa | < 0.00010 | KO Synap low, WT Synap high | 0 | 0 | 0 | 3 | 3 | 4 |
| 710 | Delta-1-pyrroline-5-carboxylate dehydrogenase, mitochondrial OS=Mus musculus GN=Aldh4a1 PE=1 SV=3 | AL4A1_MOUSE | Aldh4a1 | 62 kDa | 0.012 | KO Synap low, WT Synap high | 0 | 0 | 0 | 4 | 2 | 2 |
| 711 | Thioredoxin, mitochondrial (Fragment) OS=Mus musculus GN=Txn2 PE=1 SV=1 | G3UZY2_MOUSE (+1) | Txn2 | 14 kDa | 0.006 | KO Synap low, WT Synap high | 0 | 0 | 0 | 2 | 2 | 2 |
| 712 | Neurocalcin-delta OS=Mus musculus GN=Ncald PE=1 SV=4 | NCALD_MOUSE | Ncald | 22 kDa | 0.011 | KO Synap low, WT Synap high | 0 | 0 | 0 | 1 | 2 | 3 |
| 713 | Synapsin-3 OS=Mus musculus GN=Syn3 PE=1 SV=2 | SYN3_MOUSE | Syn3 | 63 kDa | 0.016 | KO Synap low, WT Synap high | 0 | 2 | 0 | 4 | 2 | 2 |
| 714 | Peptidyl-prolyl cis-trans isomerase D OS=Mus musculus GN=Ppid PE=1 SV=3 | PPID_MOUSE | Ppid | 41 kDa | 0.0015 | KO Synap low, WT Synap high | 0 | 0 | 1 | 1 | 1 | 2 |
| 715 | Voltage-dependent calcium channel subunit alpha-2/delta-1 OS=Mus musculus GN=Cacna2d1 PE=1 SV=1 | CA2D1_MOUSE (+1) | Cacna2d1 | 125 kDa | 0.0043 | KO Synap low, WT Synap high | 0 | 0 | 0 | 3 | 6 | 5 |
| 716 | Ankyrin repeat and sterile alpha motif domain-containing protein 1B OS=Mus musculus GN=Anks1b PE=1 SV=1 | A0A0R4J2A6_MOUSE (+9) | Anks1b | 48 kDa | 0.00068 | KO Synap low, WT Synap high | 0 | 0 | 0 | 3 | 2 | 3 |
| 717 | Glutamate decarboxylase 2 OS=Mus musculus GN=Gad2 PE=1 SV=1 | DCE2_MOUSE | Gad2 | 65 kDa | 0.0012 | KO Synap low, WT Synap high | 0 | 0 | 0 | 3 | 2 | 3 |
| 718 | Neuroligin-2 OS=Mus musculus GN=Nlgn2 PE=1 SV=2 | NLGN2_MOUSE | Nlgn2 | 91 kDa | 0.00023 | KO Synap low, WT Synap high | 1 | 1 | 1 | 1 | 3 | 3 |
| 719 | SH3 and multiple ankyrin repeat domains protein 1 OS=Mus musculus GN=Shank1 PE=1 SV=1 | D3YZU4_MOUSE (+2) | Shank1 | 225 kDa | 0.016 | KO Synap low, WT Synap high | 1 | 1 | 0 | 1 | 2 | 1 |
| 720 | Histidine triad nucleotide-binding protein 2, mitochondrial OS=Mus musculus GN=Hint2 PE=1 SV=1 | HINT2_MOUSE | Hint2 | 17 kDa | 0.024 | KO Synap low, WT Synap high | 0 | 0 | 0 | 1 | 3 | 1 |
| 721 | Ras-related protein Rab-35 OS=Mus musculus GN=Rab35 PE=1 SV=1 | RAB35_MOUSE | Rab35 | 23 kDa | 0.036 | KO Synap low, WT Synap high | 0 | 1 | 0 | 3 | 3 | 3 |
| 722 | Ras-related protein Ral-A OS=Mus musculus GN=Rala PE=1 SV=1 | RALA_MOUSE | Rala | 24 kDa | 0.0076 | KO Synap low, WT Synap high | 0 | 1 | 0 | 3 | 2 | 3 |
| 723 | Sodium channel protein OS=Mus musculus GN=Scn2a PE=1 SV=1 | A0A0J9YTW6_MOUSE (+1) | Scn2a | 228 kDa | 0.00046 | KO Synap low, WT Synap high | 1 | 0 | 0 | 3 | 3 | 3 |
| 724 | Glutaredoxin-related protein 5, mitochondrial OS=Mus musculus GN=Glrx5 PE=1 SV=2 | GLRX5_MOUSE | Glrx5 | 16 kDa | 0.00044 | KO Synap low, WT Synap high | 0 | 0 | 0 | 2 | 3 | 3 |
| 725 | OCIA domain-containing protein 1 (Fragment) OS=Mus musculus GN=Ociad1 PE=1 SV=1 | A0A0J9YUB6_MOUSE (+1) | Ociad1 | 20 kDa | 0.03 | KO Synap low, WT Synap high | 0 | 0 | 0 | 3 | 1 | 2 |
| 726 | Lymphocyte antigen 6H OS=Mus musculus GN=Ly6h PE=1 SV=2 | LY6H_MOUSE (+1) | Ly6h | 15 kDa | < 0.00010 | KO Synap low, WT Synap high | 0 | 0 | 0 | 2 | 1 | 2 |
| 727 | Contactin-associated protein-like 2 OS=Mus musculus GN=Cntnap2 PE=1 SV=2 | CNTP2_MOUSE (+1) | Cntnap2 | 148 kDa | < 0.00010 | KO Synap low, WT Synap high | 0 | 0 | 0 | 2 | 3 | 3 |
| 728 | Guanine nucleotide-binding protein G(z) subunit alpha OS=Mus musculus GN=Gnaz PE=1 SV=4 | GNAZ_MOUSE | Gnaz | 41 kDa | 0.019 | KO Synap low, WT Synap high | 0 | 2 | 1 | 2 | 2 | 2 |
| 729 | Trans-2-enoyl-CoA reductase, mitochondrial OS=Mus musculus GN=Mecr PE=1 SV=2 | MECR_MOUSE | Mecr | 40 kDa | 0.00063 | KO Synap low, WT Synap high | 0 | 0 | 0 | 2 | 2 | 3 |
| 730 | MCG9827 OS=Mus musculus GN=Opcml PE=1 SV=1 | G5E8G3_MOUSE (+1) | Opcml | 38 kDa | 0.022 | KO Synap low, WT Synap high | 0 | 0 | 0 | 2 | 2 | 2 |
| 731 | Voltage-dependent anion-selective channel protein 2 (Fragment) OS=Mus musculus GN=Vdac2 PE=1 SV=1 | G3UX26_MOUSE (+1) | Vdac2 | 30 kDa | 0.0045 | KO Synap low, WT Synap high | 0 | 0 | 1 | 2 | 3 | 3 |
| 732 | Neuroligin-3 OS=Mus musculus GN=Nlgn3 PE=1 SV=1 | A2AGI2_MOUSE (+1) | Nlgn3 | 93 kDa | 0.0059 | KO Synap low, WT Synap high | 0 | 0 | 0 | 3 | 2 | 2 |
| 733 | Anion exchange protein OS=Mus musculus GN=Slc4a10 PE=1 SV=1 | B1AWV9_MOUSE (+1) | Slc4a10 | 125 kDa | 0.0024 | KO Synap low, WT Synap high | 0 | 0 | 0 | 2 | 3 | 2 |
| 734 | ATP-dependent Clp protease proteolytic subunit, mitochondrial OS=Mus musculus GN=Clpp PE=1 SV=1 | CLPP_MOUSE | Clpp | 30 kDa | 0.0007 | KO Synap low, WT Synap high | 0 | 0 | 0 | 2 | 3 | 2 |
| 735 | Probable G-protein coupled receptor 158 OS=Mus musculus GN=Gpr158 PE=1 SV=2 | GP158_MOUSE | Gpr158 | 134 kDa | < 0.00010 | KO Synap low, WT Synap high | 0 | 0 | 0 | 3 | 4 | 3 |
| 736 | Spectrin beta 1 OS=Mus musculus GN=Sptb PE=1 SV=1 | Q3UGX2_MOUSE | Sptb | 268 kDa | 0.042 | KO Synap low, WT Synap high | 1 | 1 | 1 | 3 | 1 | 4 |
| 737 | Synaptotagmin-1 OS=Mus musculus GN=Syt1 PE=1 SV=1 | SYT1_MOUSE | Syt1 | 47 kDa | 0.0015 | KO Synap low, WT Synap high | 0 | 0 | 1 | 3 | 3 | 2 |
| 738 | 2,4-dienoyl-CoA reductase, mitochondrial OS=Mus musculus GN=Decr1 PE=1 SV=1 | DECR_MOUSE | Decr1 | 36 kDa | < 0.00010 | KO Synap low, WT Synap high | 0 | 0 | 0 | 2 | 2 | 2 |
| 739 | Neutral amino acid transporter A OS=Mus musculus GN=Slc1a4 PE=1 SV=1 | SATT_MOUSE | Slc1a4 | 56 kDa | 0.0003 | KO Synap low, WT Synap high | 0 | 0 | 0 | 1 | 1 | 1 |
| 740 | Toll-interacting protein OS=Mus musculus GN=Tollip PE=1 SV=1 | Q8C5G6_MOUSE (+1) | Tollip | 25 kDa | 0.036 | KO Synap low, WT Synap high | 1 | 1 | 1 | 2 | 2 | 2 |
| 741 | Programmed cell death protein 5 OS=Mus musculus GN=Pdcd5 PE=1 SV=3 | PDCD5_MOUSE | Pdcd5 | 14 kDa | 0.013 | KO Synap low, WT Synap high | 1 | 0 | 1 | 2 | 2 | 2 |
| 742 | Guanine nucleotide-binding protein G(s) subunit alpha isoforms XLas OS=Mus musculus GN=Gnas PE=1 SV=1 | GNAS1_MOUSE (+2) | Gnas | 122 kDa | 0.0071 | KO Synap low, WT Synap high | 1 | 0 | 1 | 1 | 1 | 1 |
| 743 | Contactin-2 OS=Mus musculus GN=Cntn2 PE=1 SV=2 | CNTN2_MOUSE | Cntn2 | 113 kDa | 0.017 | KO Synap low, WT Synap high | 0 | 1 | 1 | 2 | 2 | 2 |
| 744 | Adenosylhomocysteinase OS=Mus musculus GN=Ahcyl2 PE=1 SV=1 | F8WGT1_MOUSE (+1) | Ahcyl2 | 67 kDa | 0.0009 | KO Synap low, WT Synap high | 1 | 1 | 2 | 2 | 2 | 3 |
| 745 | Gamma-aminobutyric acid type B receptor subunit 2 OS=Mus musculus GN=Gabbr2 PE=1 SV=2 | GABR2_MOUSE | Gabbr2 | 106 kDa | 0.0085 | KO Synap low, WT Synap high | 0 | 0 | 0 | 2 | 2 | 4 |
| 746 | OX-2 membrane glycoprotein OS=Mus musculus GN=Cd200 PE=1 SV=1 | E9QMY1_MOUSE (+2) | Cd200 | 30 kDa | 0.00063 | KO Synap low, WT Synap high | 0 | 0 | 0 | 2 | 2 | 3 |
| 747 | ATP synthase subunit f, mitochondrial OS=Mus musculus GN=Atp5j2 PE=1 SV=3 | ATPK_MOUSE | Atp5j2 | 10 kDa | 0.019 | KO Synap low, WT Synap high | 0 | 0 | 0 | 1 | 2 | 2 |
| 748 | Cytochrome c oxidase subunit 5A, mitochondrial OS=Mus musculus GN=Cox5a PE=1 SV=2 | COX5A_MOUSE | Cox5a | 16 kDa | < 0.00010 | KO Synap low, WT Synap high | 0 | 0 | 0 | 1 | 1 | 1 |
| 749 | Mitochondrial import inner membrane translocase subunit TIM50 OS=Mus musculus GN=Timm50 PE=1 SV=1 | TIM50_MOUSE | Timm50 | 40 kDa | 0.00058 | KO Synap low, WT Synap high | 0 | 0 | 0 | 3 | 3 | 3 |
| 750 | Microtubule-actin cross-linking factor 1 OS=Mus musculus GN=Macf1 PE=1 SV=1 | A0A0A0MQA6_MOUSE (+6) | Macf1 | 608 kDa | 0.0043 | KO Synap low, WT Synap high | 1 | 1 | 1 | 2 | 2 | 3 |
| 751 | Contactin-associated protein 1 OS=Mus musculus GN=Cntnap1 PE=1 SV=2 | CNTP1_MOUSE | Cntnap1 | 156 kDa | 0.0027 | KO Synap low, WT Synap high | 0 | 0 | 0 | 2 | 3 | 4 |
| 752 | Mitochondrial fission 1 protein OS=Mus musculus GN=Fis1 PE=1 SV=1 | FIS1_MOUSE | Fis1 | 17 kDa | 0.048 | KO Synap low, WT Synap high | 0 | 1 | 0 | 2 | 2 | 1 |
| 753 | Electron transfer flavoprotein-ubiquinone oxidoreductase, mitochondrial OS=Mus musculus GN=Etfdh PE=1 SV=1 | ETFD_MOUSE | Etfdh | 68 kDa | 0.011 | KO Synap low, WT Synap high | 0 | 0 | 0 | 4 | 2 | 3 |
| 754 | Regulating synaptic membrane exocytosis protein 1 OS=Mus musculus GN=Rims1 PE=1 SV=2 | F6TZK4_MOUSE (+5) | Rims1 | 140 kDa | 0.027 | KO Synap low, WT Synap high | 0 | 0 | 0 | 3 | 2 | 1 |
| 755 | Protein Tmed7 OS=Mus musculus GN=Tmed7 PE=1 SV=1 | D3YZZ5_MOUSE (+1) | Tmed7 | 25 kDa | 0.0044 | KO Synap low, WT Synap high | 0 | 0 | 0 | 2 | 3 | 3 |
| 756 | ATPase family AAA domain-containing protein 3 OS=Mus musculus GN=Atad3 PE=1 SV=1 | ATAD3_MOUSE | Atad3 | 67 kDa | 0.027 | KO Synap low, WT Synap high | 0 | 0 | 0 | 2 | 1 | 1 |
| 757 | Hydroxyacyl-coenzyme A dehydrogenase, mitochondrial OS=Mus musculus GN=Hadh PE=1 SV=2 | HCDH_MOUSE | Hadh | 34 kDa | 0.0024 | KO Synap low, WT Synap high | 0 | 0 | 0 | 1 | 2 | 1 |
| 758 | Metabotropic glutamate receptor 3 OS=Mus musculus GN=Grm3 PE=1 SV=1 | GRM3_MOUSE | Grm3 | 99 kDa | 0.0021 | KO Synap low, WT Synap high | 0 | 0 | 0 | 3 | 2 | 4 |
| 759 | Palmitoyl-protein thioesterase 1 OS=Mus musculus GN=Ppt1 PE=1 SV=2 | PPT1_MOUSE | Ppt1 | 34 kDa | 0.00044 | KO Synap low, WT Synap high | 0 | 0 | 0 | 2 | 3 | 3 |
| 760 | Citrate lyase subunit beta-like protein, mitochondrial OS=Mus musculus GN=Clybl PE=1 SV=2 | CLYBL_MOUSE | Clybl | 38 kDa | 0.00063 | KO Synap low, WT Synap high | 0 | 0 | 0 | 2 | 2 | 2 |
| 761 | Ribosome-recycling factor, mitochondrial OS=Mus musculus GN=Mrrf PE=1 SV=1 | RRFM_MOUSE | Mrrf | 29 kDa | 0.00063 | KO Synap low, WT Synap high | 0 | 0 | 0 | 2 | 2 | 3 |
| 762 | Cytochrome b-c1 complex subunit 9 OS=Mus musculus GN=Uqcr10 PE=1 SV=1 | QCR9_MOUSE | Uqcr10 | 7 kDa | 0.024 | KO Synap low, WT Synap high | 0 | 0 | 0 | 1 | 1 | 1 |
| 763 | Dolichyl-diphosphooligosaccharide--protein glycosyltransferase subunit 1 OS=Mus musculus GN=Rpn1 PE=1 SV=1 | RPN1_MOUSE | Rpn1 | 69 kDa | 0.02 | KO Synap low, WT Synap high | 0 | 0 | 1 | 2 | 2 | 2 |
| 764 | Glutamate receptor 3 OS=Mus musculus GN=Gria3 PE=1 SV=1 | B0QZW1_MOUSE (+1) | Gria3 | 100 kDa | 0.0024 | KO Synap low, WT Synap high | 0 | 0 | 0 | 2 | 2 | 2 |
| 765 | Mitochondrial import inner membrane translocase subunit Tim13 OS=Mus musculus GN=Timm13 PE=1 SV=1 | TIM13_MOUSE | Timm13 | 10 kDa | 0.0044 | KO Synap low, WT Synap high | 0 | 0 | 0 | 1 | 1 | 2 |
| 766 | L-2-hydroxyglutarate dehydrogenase, mitochondrial OS=Mus musculus GN=L2hgdh PE=1 SV=1 | L2HDH_MOUSE | L2hgdh | 51 kDa | 0.0019 | KO Synap low, WT Synap high | 0 | 0 | 0 | 2 | 2 | 3 |
| 767 | Prohibitin OS=Mus musculus GN=Phb PE=1 SV=1 | PHB_MOUSE | Phb | 30 kDa | < 0.00010 | KO Synap low, WT Synap high | 0 | 0 | 0 | 1 | 1 | 1 |
| 768 | NADH dehydrogenase [ubiquinone] iron-sulfur protein 4, mitochondrial OS=Mus musculus GN=Ndufs4 PE=1 SV=1 | E9QPX3_MOUSE (+1) | Ndufs4 | 20 kDa | 0.0085 | KO Synap low, WT Synap high | 0 | 0 | 0 | 2 | 1 | 2 |
| 769 | Large neutral amino acids transporter small subunit 1 OS=Mus musculus GN=Slc7a5 PE=1 SV=2 | LAT1_MOUSE | Slc7a5 | 56 kDa | 0.00063 | KO Synap low, WT Synap high | 0 | 0 | 0 | 1 | 1 | 1 |
| 770 | Neuroligin 4-like OS=Mus musculus GN=Nlgn4l PE=1 SV=1 | NLGN4_MOUSE | Nlgn4l | 97 kDa | 0.0003 | KO Synap low, WT Synap high | 1 | 1 | 1 | 1 | 1 | 1 |
| 771 | Serine/threonine-protein kinase BRSK1 OS=Mus musculus GN=Brsk1 PE=1 SV=1 | BRSK1_MOUSE (+1) | Brsk1 | 85 kDa | 0.016 | KO Synap low, WT Synap high | 0 | 0 | 0 | 1 | 2 | 3 |
| 772 | Actin-related protein 2/3 complex subunit 3 OS=Mus musculus GN=Arpc3 PE=1 SV=3 | ARPC3_MOUSE | Arpc3 | 21 kDa | 0.0085 | KO Synap low, WT Synap high | 0 | 0 | 0 | 1 | 1 | 2 |
| 773 | Mitofusin-2 OS=Mus musculus GN=Mfn2 PE=1 SV=3 | MFN2_MOUSE | Mfn2 | 86 kDa | 0.016 | KO Synap low, WT Synap high | 0 | 0 | 0 | 1 | 2 | 3 |
| 774 | Cysteine desulfurase, mitochondrial OS=Mus musculus GN=Nfs1 PE=1 SV=3 | NFS1_MOUSE | Nfs1 | 51 kDa | 0.049 | KO Synap low, WT Synap high | 0 | 0 | 0 | 1 | 1 | 2 |
| 775 | ARF6 guanine nucleotide exchange factor IQArfGEF OS=Mus musculus GN=Iqsec2 PE=1 SV=1 | A4GZ26_MOUSE (+3) | Iqsec2 | 162 kDa | 0.027 | KO Synap low, WT Synap high | 0 | 0 | 0 | 3 | 2 | 2 |
| 776 | MICOS complex subunit MIC60 OS=Mus musculus GN=Immt PE=1 SV=1 | A0A0U1RP81_MOUSE | Immt | 54 kDa | 0.00019 | KO Synap low, WT Synap high | 0 | 0 | 0 | 1 | 1 | 1 |
| 777 | Glutamate receptor 2 OS=Mus musculus GN=Gria2 PE=1 SV=1 | E9QKC0_MOUSE (+2) | Gria2 | 99 kDa | 0.0023 | KO Synap low, WT Synap high | 0 | 0 | 0 | 1 | 1 | 1 |
| 778 | NADH dehydrogenase [ubiquinone] iron-sulfur protein 5 OS=Mus musculus GN=Ndufs5 PE=1 SV=3 | NDUS5_MOUSE | Ndufs5 | 13 kDa | 0.0027 | KO Synap low, WT Synap high | 0 | 0 | 0 | 2 | 3 | 3 |
| 779 | Endophilin-A3 OS=Mus musculus GN=Sh3gl3 PE=1 SV=1 | A0A0R4J0B8_MOUSE (+1) | Sh3gl3 | 39 kDa | 0.027 | KO Synap low, WT Synap high | 0 | 0 | 0 | 2 | 1 | 1 |
| 780 | Potassium voltage-gated channel subfamily A member 1 OS=Mus musculus GN=Kcna1 PE=1 SV=1 | KCNA1_MOUSE | Kcna1 | 56 kDa | 0.012 | KO Synap low, WT Synap high | 1 | 0 | 1 | 1 | 2 | 2 |
| 781 | Pyridoxal phosphate phosphatase OS=Mus musculus GN=Pdxp PE=1 SV=1 | PLPP_MOUSE | Pdxp | 32 kDa | 0.0044 | KO Synap low, WT Synap high | 0 | 0 | 0 | 1 | 2 | 2 |
| 782 | Up-regulated during skeletal muscle growth protein 5 OS=Mus musculus GN=Usmg5 PE=1 SV=1 | USMG5_MOUSE | Usmg5 | 6 kDa | 0.0024 | KO Synap low, WT Synap high | 0 | 0 | 0 | 2 | 2 | 2 |
| 783 | Monoacylglycerol lipase ABHD12 OS=Mus musculus GN=Abhd12 PE=1 SV=2 | ABD12_MOUSE (+1) | Abhd12 | 45 kDa | 0.0074 | KO Synap low, WT Synap high | 0 | 0 | 1 | 1 | 2 | 2 |
| 784 | Catechol O-methyltransferase domain-containing protein 1 OS=Mus musculus GN=Comtd1 PE=1 SV=1 | CMTD1_MOUSE | Comtd1 | 29 kDa | 0.049 | KO Synap low, WT Synap high | 0 | 0 | 0 | 1 | 1 | 3 |
| 785 | Arfaptin-2 OS=Mus musculus GN=Arfip2 PE=1 SV=2 | ARFP2_MOUSE (+2) | Arfip2 | 38 kDa | 0.032 | KO Synap low, WT Synap high | 0 | 1 | 0 | 1 | 2 | 1 |
| 786 | Serine/threonine-protein kinase MARK2 (Fragment) OS=Mus musculus GN=Mark2 PE=1 SV=1 | F6ZS70_MOUSE (+7) | Mark2 | 87 kDa | 0.019 | KO Synap low, WT Synap high | 0 | 0 | 0 | 2 | 1 | 3 |
| 787 | MICOS complex subunit (Fragment) OS=Mus musculus GN=Chchd3 PE=1 SV=1 | D3Z0L4_MOUSE | Chchd3 | 22 kDa | 0.00038 | KO Synap low, WT Synap high | 0 | 1 | 0 | 1 | 1 | 1 |
| 788 | Leucine-rich repeat-containing protein 7 OS=Mus musculus GN=Lrrc7 PE=1 SV=1 | A0A0G2JDT9_MOUSE (+4) | Lrrc7 | 168 kDa | 0.0087 | KO Synap low, WT Synap high | 0 | 0 | 0 | 1 | 1 | 2 |
| 789 | Long-chain-fatty-acid--CoA ligase 1 OS=Mus musculus GN=Acsl1 PE=1 SV=2 | ACSL1_MOUSE (+1) | Acsl1 | 78 kDa | 0.0085 | KO Synap low, WT Synap high | 0 | 0 | 0 | 1 | 1 | 2 |
| 790 | GTPase NRas (Fragment) OS=Mus musculus GN=Nras PE=1 SV=1 | A0A0G2JGP4_MOUSE (+2) | Nras | 21 kDa | 0.00022 | KO Synap low, WT Synap high | 0 | 0 | 0 | 1 | 1 | 1 |
| 791 | Thioredoxin-related transmembrane protein 2 OS=Mus musculus GN=Tmx2 PE=1 SV=1 | D3Z2J6_MOUSE (+1) | Tmx2 | 30 kDa | 0.024 | KO Synap low, WT Synap high | 0 | 0 | 0 | 1 | 2 | 2 |
| 792 | ATPase family AAA domain-containing protein 1 OS=Mus musculus GN=Atad1 PE=1 SV=1 | ATAD1_MOUSE | Atad1 | 41 kDa | 0.027 | KO Synap low, WT Synap high | 0 | 0 | 0 | 2 | 1 | 1 |
| 793 | Calcium-transporting ATPase OS=Mus musculus GN=Atp2b2 PE=1 SV=1 | S4R1C4_MOUSE | Atp2b2 | 127 kDa | < 0.00010 | KO Synap low, WT Synap high | 1 | 1 | 1 | 1 | 1 | 1 |
| 794 | WD repeat-containing protein 37 OS=Mus musculus GN=Wdr37 PE=1 SV=1 | WDR37_MOUSE | Wdr37 | 55 kDa | 0.016 | KO Synap low, WT Synap high | 0 | 0 | 0 | 1 | 1 | 1 |
| 795 | Cytochrome b5 OS=Mus musculus GN=Cyb5a PE=1 SV=2 | CYB5_MOUSE (+1) | Cyb5a | 15 kDa | 0.0085 | KO Synap low, WT Synap high | 0 | 0 | 0 | 1 | 1 | 2 |

| | | | | | | | | | | | | |
|---|---|---|---|---|---|---|---|---|---|---|---|---|
| 796 | Regulating synaptic membrane exocytosis protein 1 (Fragment) OS=Mus musculus GN=Rims1 PE=1 SV=3 | E9PWP6_MOUSE | Rims1 | 89 kDa | 0.0014 | KO Synap low, WT Synap high | 0 | 0 | 0 | 1 | 1 | 1 |
| 797 | Peptidyl-prolyl cis-trans isomerase FKBP8 OS=Mus musculus GN=Fkbp8 PE=1 SV=2 | FKBP8_MOUSE | Fkbp8 | 44 kDa | 0.027 | KO Synap low, WT Synap high | 0 | 0 | 0 | 2 | 1 | 1 |
| 798 | Radixin OS=Mus musculus GN=Rdx PE=1 SV=3 | RADI_MOUSE | Rdx | 69 kDa | 0.0085 | KO Synap low, WT Synap high | 0 | 0 | 0 | 1 | 1 | 1 |
| 799 | Leucine-rich repeat and immunoglobulin-like domain-containing nogo receptor-interacting protein 1 OS=Mus musculus GN=Lingo1 PE=1 SV=1 | A9DA50_MOUSE | Lingo1 | 70 kDa | 0.016 | KO Synap low, WT Synap high | 0 | 0 | 0 | 1 | 1 | 1 |
| 800 | Metabotropic glutamate receptor 2 OS=Mus musculus GN=Grm2 PE=1 SV=2 | GRM2_MOUSE | Grm2 | 96 kDa | < 0.00010 | KO Synap low, WT Synap high | 0 | 0 | 0 | 2 | 2 | 2 |
| 801 | Aspartyl/asparaginyl beta-hydroxylase OS=Mus musculus GN=Asph PE=1 SV=1 | A2AL78_MOUSE (+7) | Asph | 26 kDa | 0.0085 | KO Synap low, WT Synap high | 0 | 0 | 0 | 1 | 1 | 2 |
| 802 | Dehydrogenase/reductase SDR family member 1 OS=Mus musculus GN=Dhrs1 PE=1 SV=1 | DHRS1_MOUSE | Dhrs1 | 34 kDa | 0.028 | KO Synap low, WT Synap high | 0 | 0 | 0 | 2 | 3 | 1 |
| 803 | Ras-related protein Rab-3B OS=Mus musculus GN=Rab3b PE=1 SV=1 | RAB3B_MOUSE | Rab3b | 25 kDa | 0.0058 | KO Synap low, WT Synap high | 0 | 0 | 0 | 1 | 1 | 1 |
| 804 | Phosphatidylinositol 4-kinase alpha OS=Mus musculus GN=Pi4ka PE=1 SV=2 | PI4KA_MOUSE | Pi4ka | 237 kDa | 0.028 | KO Synap low, WT Synap high | 0 | 0 | 0 | 2 | 3 | 1 |
| 805 | Apoptosis-inducing factor 1, mitochondrial OS=Mus musculus GN=Aifm1 PE=1 SV=1 | AIFM1_MOUSE (+1) | Aifm1 | 67 kDa | < 0.00010 | KO Synap low, WT Synap high | 0 | 0 | 0 | 1 | 1 | 1 |
| 806 | Synaptogyrin-1 OS=Mus musculus GN=Syngr1 PE=1 SV=2 | SNG1_MOUSE | Syngr1 | 26 kDa | < 0.00010 | KO Synap low, WT Synap high | 0 | 0 | 0 | 1 | 1 | 1 |
| 807 | Bcl-2-like protein 13 OS=Mus musculus GN=Bcl2l13 PE=1 SV=2 | B2L13_MOUSE | Bcl2l13 | 47 kDa | < 0.00010 | KO Synap low, WT Synap high | 0 | 0 | 0 | 1 | 1 | 1 |
| 808 | Versican core protein OS=Mus musculus GN=Vcan PE=1 SV=2 | CSPG2_MOUSE (+5) | Vcan | 367 kDa | 0.027 | KO Synap low, WT Synap high | 0 | 0 | 0 | 2 | 1 | 1 |
| 809 | Tricarboxylate transport protein, mitochondrial OS=Mus musculus GN=Slc25a1 PE=1 SV=1 | TXTP_MOUSE | Slc25a1 | 34 kDa | < 0.00010 | KO Synap low, WT Synap high | 0 | 0 | 0 | 2 | 2 | 2 |
| 810 | 28S ribosomal protein S28, mitochondrial OS=Mus musculus GN=Mrps28 PE=1 SV=1 | RT28_MOUSE | Mrps28 | 21 kDa | 0.037 | KO Synap low, WT Synap high | 0 | 0 | 0 | 2 | 2 | 1 |
| 811 | Non-specific lipid-transfer protein OS=Mus musculus GN=Scp2 PE=1 SV=3 | NLTP_MOUSE | Scp2 | 59 kDa | < 0.00010 | KO Synap low, WT Synap high | 0 | 0 | 0 | 1 | 1 | 1 |
| 812 | Ras-related protein Rab-33B OS=Mus musculus GN=Rab33b PE=1 SV=1 | RB33B_MOUSE | Rab33b | 26 kDa | 0.00081 | KO Synap low, WT Synap high | 0 | 0 | 0 | 1 | 1 | 1 |
| 813 | Receptor-type tyrosine-protein phosphatase alpha OS=Mus musculus GN=Ptpra PE=1 SV=3 | PTPRA_MOUSE (+1) | Ptpra | 94 kDa | < 0.00010 | KO Synap low, WT Synap high | 0 | 0 | 0 | 1 | 1 | 1 |
| 814 | Acyl-coenzyme A thioesterase THEM4 OS=Mus musculus GN=Them4 PE=1 SV=1 | THEM4_MOUSE | Them4 | 26 kDa | < 0.00010 | KO Synap low, WT Synap high | 0 | 0 | 0 | 1 | 1 | 1 |
| 815 | Serine/threonine-protein phosphatase 2A catalytic subunit alpha isoform OS=Mus musculus GN=Ppp2ca PE=1 SV=1 | PP2AA_MOUSE (+1) | Ppp2ca | 36 kDa | 0.0085 | KO Synap low, WT Synap high | 0 | 0 | 0 | 1 | 1 | 2 |
| 816 | Voltage-dependent calcium channel subunit alpha-2/delta-2 OS=Mus musculus GN=Cacna2d2 PE=1 SV=2 | E9Q683_MOUSE (+1) | Cacna2d2 | 131 kDa | 0.027 | KO Synap low, WT Synap high | 0 | 0 | 0 | 2 | 1 | 1 |
| 817 | UPF0598 protein C8orf82 homolog OS=Mus musculus PE=1 SV=1 | CH082_MOUSE | | 24 kDa | < 0.00010 | KO Synap low, WT Synap high | 0 | 0 | 0 | 1 | 1 | 1 |
| 818 | Tumor protein D52 (Fragment) OS=Mus musculus GN=Tpd52 PE=1 SV=1 | D3Z125_MOUSE (+5) | Tpd52 | 19 kDa | < 0.00010 | KO Synap low, WT Synap high | 0 | 0 | 0 | 1 | 1 | 1 |
| 819 | Neuronal proto-oncogene tyrosine-protein kinase Src OS=Mus musculus GN=Src PE=1 SV=1 | F8WI90_MOUSE (+6) | Src | 60 kDa | < 0.00010 | KO Synap low, WT Synap high | 0 | 0 | 0 | 1 | 1 | 1 |
| 820 | Hydroxymethylglutaryl-CoA lyase, mitochondrial OS=Mus musculus GN=Hmgcl PE=1 SV=2 | HMGCL_MOUSE | Hmgcl | 34 kDa | < 0.00010 | KO Synap low, WT Synap high | 0 | 0 | 0 | 1 | 1 | 1 |
| 821 | Vacuolar protein sorting-associated protein 51 homolog OS=Mus musculus GN=Vps51 PE=1 SV=2 | VPS51_MOUSE | Vps51 | 86 kDa | < 0.00010 | KO Synap low, WT Synap high | 0 | 0 | 0 | 1 | 1 | 1 |
| 822 | Thioredoxin OS=Mus musculus GN=Txn PE=1 SV=3 | THIO_MOUSE | Txn | 12 kDa | 0.0085 | KO Synap low, WT Synap high | 0 | 0 | 0 | 1 | 1 | 2 |
| 823 | Cathepsin L1 OS=Mus musculus GN=Ctsl PE=1 SV=2 | CATL1_MOUSE | Ctsl | 38 kDa | 0.0087 | KO Synap low, WT Synap high | 0 | 0 | 0 | 1 | 1 | 1 |
| 824 | Protein phosphatase 1 regulatory subunit 21 OS=Mus musculus GN=Ppp1r21 PE=1 SV=2 | PPR21_MOUSE | Ppp1r21 | 88 kDa | < 0.00010 | KO Synap low, WT Synap high | 0 | 0 | 0 | 1 | 1 | 1 |
| 825 | CD166 antigen (Fragment) OS=Mus musculus GN=Alcam PE=1 SV=1 | F6QH25_MOUSE (+3) | Alcam | 38 kDa | < 0.00010 | KO Synap low, WT Synap high | 0 | 0 | 0 | 1 | 1 | 1 |
| 826 | Actin-related protein 3B OS=Mus musculus GN=Actr3b PE=1 SV=1 | ARP3B_MOUSE | Actr3b | 48 kDa | 0.024 | KO Synap low, WT Synap high | 0 | 0 | 0 | 1 | 1 | 1 |
| 827 | Ras-related protein Rab-8A OS=Mus musculus GN=Rab8a PE=1 SV=2 | RAB8A_MOUSE | Rab8a | 24 kDa | 0.00057 | KO Synap low, WT Synap high | 0 | 0 | 0 | 1 | 1 | 1 |
| 828 | SH2 domain-containing adapter protein F (Fragment) OS=Mus musculus GN=Shf PE=1 SV=1 | A2AQ87_MOUSE (+1) | Shf | 47 kDa | < 0.00010 | KO Synap low, WT Synap high | 0 | 0 | 0 | 1 | 1 | 1 |
| 829 | Methionine--tRNA ligase, mitochondrial OS=Mus musculus GN=Mars2 PE=1 SV=2 | SYMM_MOUSE | Mars2 | 66 kDa | < 0.00010 | KO Synap low, WT Synap high | 0 | 0 | 0 | 1 | 1 | 1 |
